# Identification of a *Prunus* MAX1 Homolog as a Unique Strigol Synthase from Carlactone Bypassing 5-Deoxystrigol

**DOI:** 10.1101/2022.10.24.513630

**Authors:** Sheng Wu, Anqi Zhou, Kozue Hiugano, Akiyoshi Yoda, Xiaonan Xie, Kenji Yamane, Kenji Miura, Takahito Nomura, Yanran Li

**Affiliations:** Department of Chemical and Environmental Engineering, University of California, Riverside, California 92521, USA; Center for Bioscience Research and Education, Utsunomiya University, Tochigi 321-8505, Japan; School of Agriculture, Utsunomiya University, Tochigi 321-8505, Japan; United Graduate School of Agricultural Science, Tokyo University of Agriculture and Technology, Tokyo 183-8509, Japan; Graduate School of Life and Environmental Sciences, University of Tsukuba, Tsukuba, 305-8572, Japan

**Author notes:** Correspondence should be addressed to Yanran Li, Takahito Nomura, E-mail:.

## Abstract

Strigol was the first strigolactone (SL) to be discovered, but the biosynthetic pathway remains elusive. Here, through rapid gene screening using a microbial SL-producing platform, we functionally identified a strigol synthase (PpMAX1c, a cytochrome P450 711A enzyme) in *Prunus* that synthesizes strigol directly from the SL precursor carlactone through catalyzing multi-step oxidations and C-ring cyclization, bypassing the synthesis of 5-deoxystrigol. The function of PpMAX1c was validated through reconstructing the biosynthesis of strigol in *Nicotiana benthamiana*. Additional genomic analysis and functional verification confirm that peach also encodes an orobanchol synthase (PpCYP722C, a cytochrome P450 722C enzyme), which hints at the presence of both strigol-type and orobanchol-type SLs in peach and was confirmed through metabolic analysis of peach seedlings. This work highlights the catalytic diversity of the largely unexplored family of CYP711A homologs and sets the foundation to characterize the roles of different types of SLs in the economically important *Prunus*.

## Introduction

Strigolactones (SLs) are a class of plant hormones that play essential roles in plant growth, development, and communication with soil microbes (1–3). To date, more than 30 natural SLs have been reported, and have been classified into two groups according to their ring scaffold: canonical and noncanonical SLs (4, 5). Canonical SLs have a four-ring structure with a butanolide ring (D ring) and a tricyclic lactone (ABC ring) connected through a enol ether bond (3), and can be further divided into strigol (*S*)-type and orobanchol (*O*)-type SLs according to the different stereo configuration of the BC ring (Fig. 1) (5). Most plants have been found to generally produce one type of canonical SLs, with quite a few exceptions reported to produce both types but inconclusively (5, 6).

**Fig. 1.**
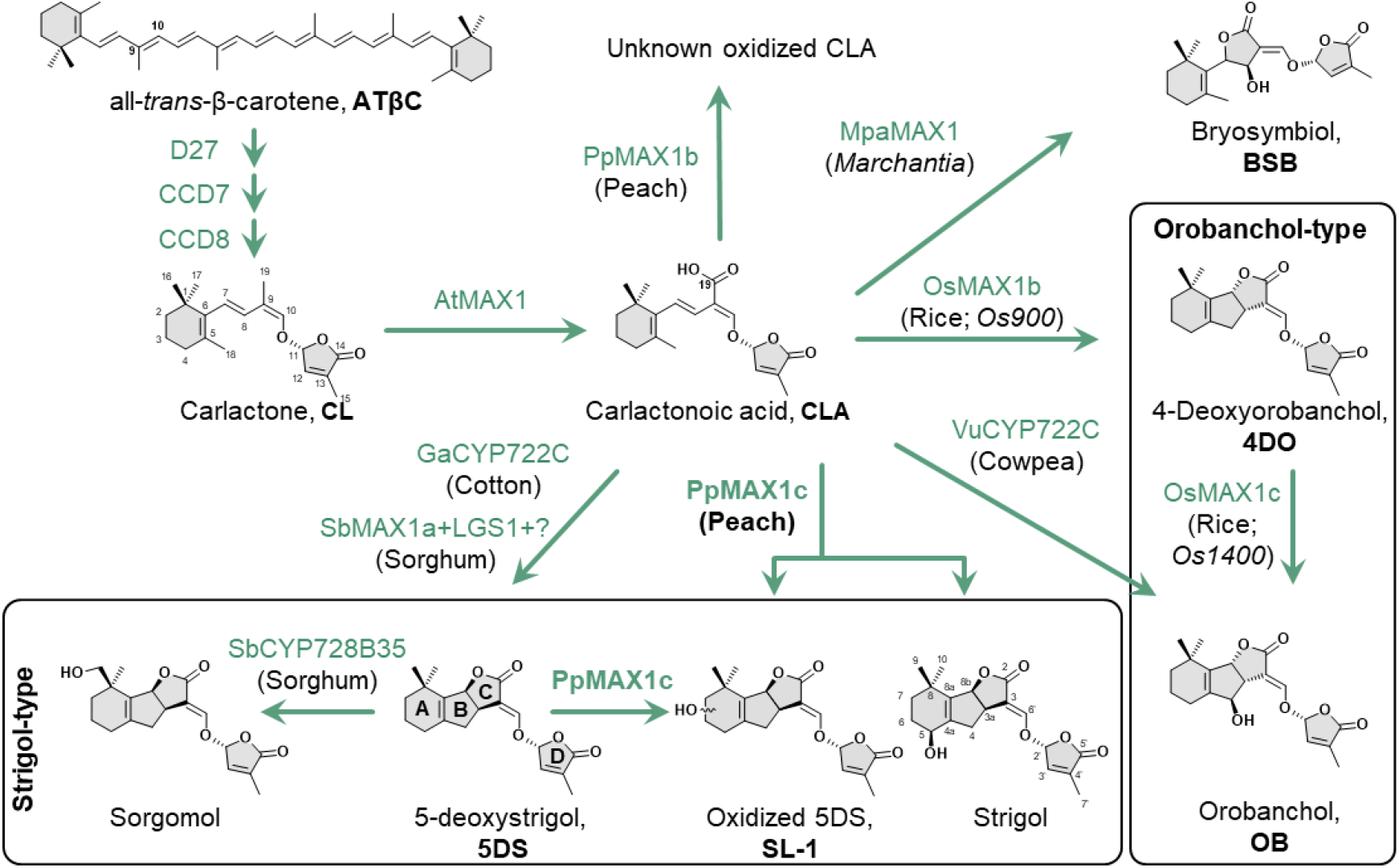
The biosynthetic pathway of strigol in *Prunus persica* and the other strigolactones in seed plants and bryophytes. Abbreviations: D27, DWARF27, a [2Fe-2S]-containing isomerase; CCD7, carotenoid cleavage dioxygenase 7; CCD8, carotenoid cleavage dioxygenase 8; AtMAX1, *MORE AXILLARY GROWTH 1* from *A. thaliana* (GenBank accession number: NP_565617); PpMAX1c, MAX1 analog c from *P. persica* (XP_007225050); PpMAX1b, MAX1 analog b from *P. persica* (XP_007224581); VuCYP722C, CYP722C from *V. unguiculata* (XP_027918387); GaCYP722C, CYP722C from *G. arboretum* (XP_016745621); OsMAX1b, MAX1 analog b (CYP711A2) from *O. sativa* (XP_015633367); OsMAX1c, MAX1 analog c (CYP711A3) from *O. sativa* (XP_015644699); MpaMAX1, MAX1 from *M. paleacea* (BCG55903); SbMAX1a, MAX1 analog a from *S. bicolor* (XP_002458367); LGS1, *LOW GERMINATION STIMULANT 1*, a sulfotransferase from *S. bicolor* (KAG0530922); SbCYP728B35, cytochrome P450 CYP728B subfamily from *S. bicolor* (XP_002443327).

SLs are apocarotenoids derived from β-carotene through a key intermediate carlactone (CL, Fig. 1) (7), which can then be converted to various SL structures with the function of cytochrome P450s and other accessory enzymes such as methyltransferases and 2-oxoglutarate-dependent dioxygenases (8–12). SL biosynthesis can be diverse and species-dependent (13, 14). Taking the biosynthesis of orobanchol (OB) as an example, rice has been identified to produce both 4-deoxyorobanchol (4DO) and OB (15), yet 4DO cannot be detected from many other OB-producing plants such as cowpea (16). One recent investigation on the cytochrome P450 MORE AXILLARY GROWTH1 (MAX1, belonging to CYP711A) homologs in rice indicates that OsCYP711A2 (Os900/OsMAX1b in Fig. 1, *Os01g0700900*) is responsible for converting CL to 4DO through carlactonoic acid (CLA) and OsCYP711A3 (Os1400/OsMAX1c in Fig. 1, *Os01g0701400*) subsequently oxidizes 4DO to afford the synthesis of OB (Fig. 1) (17). More recently, CYP722C from tomato and cowpea have been found to be involved in the conversion of CLA to OB (Fig. 1) (12). Meanwhile, CYP722C from cotton was identified to convert CLA to 5DS (Fig. 1) (18).

Strigol, the first identified strigolactone, was originally isolated and identified from cotton root exudates (*Gossypium hirsutum*) (19, 20) as a germination stimulant for root parasitic weeds (20). Correspondingly, strigol was later on found to exist widely in *Striga* hosts such as maize (*Zea mays*), sorghum (*Sorghum bicolor*), and proso millet (*Panicum miliaceum*) (21). Strigol and 5-deoxystrigol (5DS) are structurally very similar with the only difference in the hydroxyl group at the C5 position in the A ring (Fig. 1). The recent pioneering *in planta* study showed that when 5DS is fed to cotton, strigol can be detected (Fig. S1, S2) (16); while another strigol-accumulating plant moonseed (*Menispermum dauricum* DC., Fig. S1) cannot synthesize 5DS nor convert 5DS to strigol (16). A putative hydroxyl CL was detected in moonseed and proposed to be 4-hydroxy carlactone, which might be the precursor to strigol in moonseed (Fig. S1) (16). Thus, there has been a hypothesis that at least two distinct strigol biosynthetic pathways exist in nature, through 5DS or bypass 5DS, like the biosynthesis of OB (22).

The genus *Prunus* includes several economically important members such as peach, apricots, plums, and almond, and generally encode multiple MAX1 homologs (Fig. S3). Little is known about SL profiles, biosynthesis, and functions in *Prunus* trees. Here, harnessing the recently established SL-producing microbial consortium (23), we examined the functions of MAX1 homologs encoded by peach, and uncovered a special MAX1 homolog (PpMAX1c) that can directly convert CL into strigol and another oxygenated 5DS derivative *via* the intermediate 18-hydroxy-carlactonoic acid (18-OH-CLA), the function of which was further confirmed in *Nicotiana benthamiana*. In addition, the function of CYP722C encoded by peach was also confirmed to be involved in OB biosynthesis, which indicates that peach can produce both strigol and orobanchol. The SLs present in peach seedlings were analyzed and confirmed to be a mixture of both (*S*)- and (*O*)-type SLs. The identification of the strigol synthase PpMAX1c from *P. persica* expands our understanding of SL biosynthesis and the function of MAX1. We have also established a strigol-producing microbial consortium at the titer of 71.82±6.93 μg/L, providing a platform to facilitate the investigation on biosynthetic enzymes that functions subsequent to strigol.

## Results

### Peach encodes multiple MAX1 homologs distinct from previously identified MAX1s

MAX1 was first identified in *Arabidopsis* catalyzing the oxidation of CL at C-19 to produce CLA (24). MAX1 gene(s) are conserved among land plants (11, 25). MAX1 homologs are functionally diverse, especially when multiple MAX1 homologs are encoded by one plant species (11, 17, 18, 26, 27). For example, cereal crops generally encode more than one MAX1 homologs (28), with some identified to catalyze the conversion of CL to CLA (a step conserved in all the plants reported so far) (11, 24), some CL to 18-OH-CLA (26, 27, 29), some CL to 4DO (11, 17), and some 4DO to OB (Fig. 1) (11, 17). No MAX1 analog has been reported to convert CL to strigol-type SL. Through searching the Genome Database in Phytozome and GenBank, we found that many fruit crops encode multiple MAX1 homologs and are phylogenetically distinct from previously identified MAX1s (Fig. 2, Fig. S3, Table S1). For example, *P. persica* encodes three MAX1 homologs: PpMAX1a (GenBank accession number: XP_007222310.1), PpMAX1b (XP_007224581.1) and PpMAX1c (XP_007225050.2). Similarly, woodland strawberry (*Fragaria vesca*) encodes two: FveMAX1a1 (Gene IDs, FvH4_2g31680.t2) and FveMAX1a2 (FvH4_2g31660.t1); and apple (*Malus domestica*) encodes four MAX1 homologs: MdMAX1a1 (XP_008393629), MdMAX1a2 (XP_028955031), MdMAX1b1 (XP_008357300) and MdMAX1b2 (RXH72971). Their amino acid sequences are also similar to AtMAX1 from *Arabidopsis* (68-74% amino acid identity). According to the evolutionary analysis, the *Rosaceae* family encodes three distinct MAX1 subclades (group I, II and III). Peach encodes 3 MAX1 genes each fall in one of these three different subclades; apple encodes 4 MAX1 genes but only of two subclades (group I and II, not III); woodland strawberry encodes two highly similar MAX1 genes that belong to group I only (Fig. 2A). The third clade (group III) is quite unique and only represented in *Prunus* species, which is likely evolved from group I or II.

**Fig. 2.**
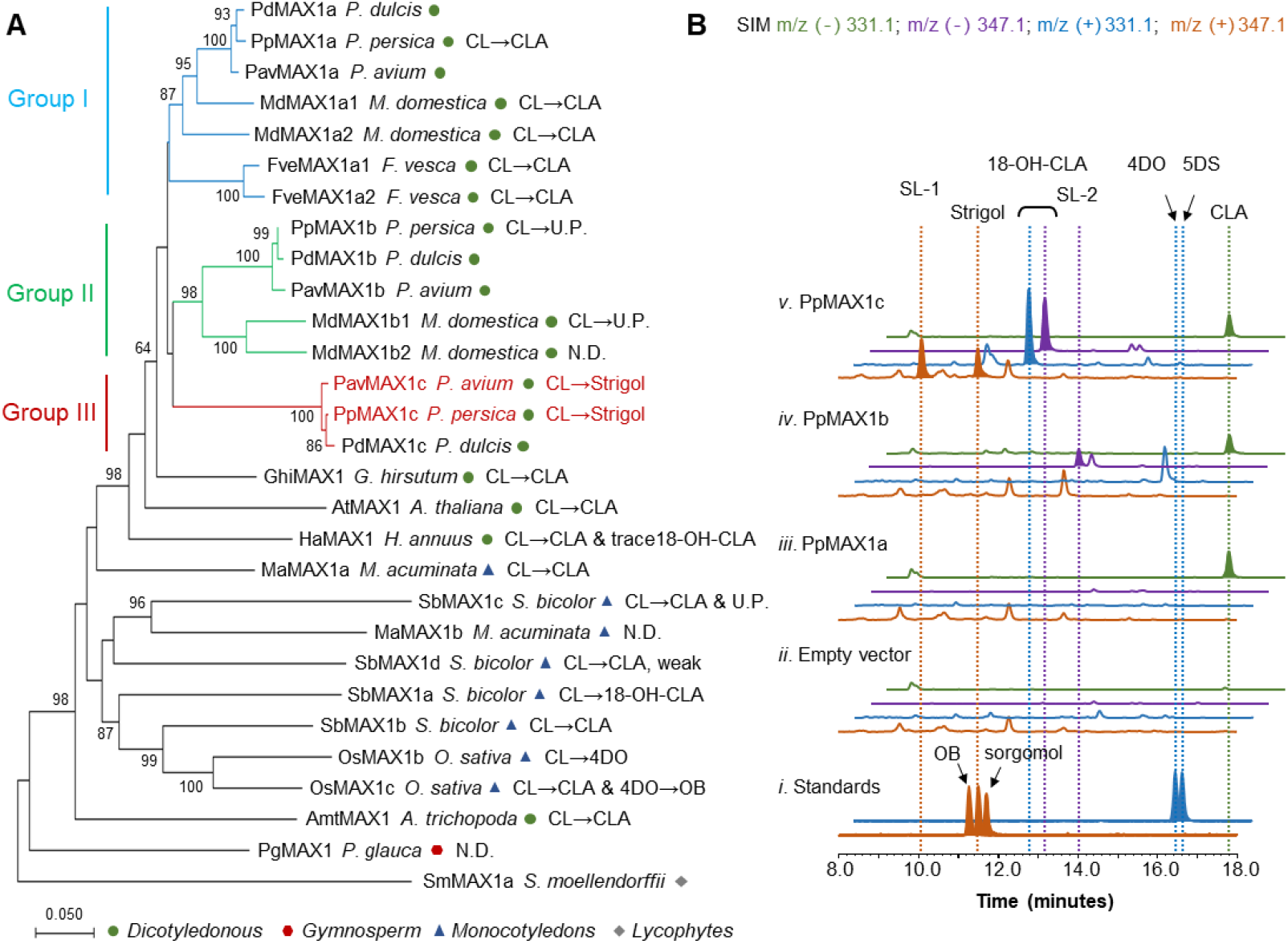
Functional characterization of PpMAX1c as a strigol synthase using SL-producing microbial consortium. (A) Phylogenetic analysis of MAX1 protein analogs. The phylogenetic tree was constructed by MEGA X using neighbor-joining method (90% partial deletion, 5000 bootstraps, p-distance mode, bootstrap values >60% are shown). A total of 29 MAX1 analogs from both monocotyledonous and dicotyledonous were selected for the analysis. The Genbank accession numbers can be found in Table S5. MAX1 with identified functions are marked with their functions. (B) Selected ion monitoring (SIM) extracted ion chromatogram (EIC) at m/z^-^ = 331.1 (green), m/z^-^ 347.1 (purple), m/z^+^ = 331.1 (blue), and m/z^+^ = 347.1 (orange) of (i) strigol, sorgomol, OB, 4DO, and 5DS standard; CL-producing *E. coli* co-cultured with yeast expressing ATR1 and (ii) an empty vector, (iii) PpMAX1a, (iv) PpMAX1b, (v) PpMAX1c. The characteristic m/z^+^ of strigol signal (*MW* = 346.38) is [C_19_H_22_O_6_ + H]^+^ = [C_19_H_23_O_6_]^+^ = 347.1. The characteristic m/z+ signal of 4DO and 5DS (*MW* = 330.38) is [C_19_H_22_O_5_ + H]^+^ = [C_19_H_23_O_5_] ^+^ = 331.1. Strain used for analysis: PpMAX1a-c (ECL/YSL1a–c; Table S3). Data are representative of at least three biological replicates.

### Functional identification of MAX1 homologs from peach using SL-producing microbial consortium

Recently, we have developed a series of SL-producing microbial consortium for the bioproduction of CL, CLA, 5DS, 4DO and OB, which also enables efficient functional identification of SL biosynthetic enzymes (23). Harnessing the CL-producing *Escherichia coli* strain (ECL; Table S2, S3), we tested the functions of the MAX1 homologs from peach. Each gene was co-expressed with the cytochrome P450 reductase 1 from *Arabidopsis thaliana* (ATR1) in *Saccharomyces cerevisiae* on a low copy number plasmid and expressed downstream of the strong, constitutive *GPD* promoter (resulting strain: YSL1a-c; Table S2, S3). The corresponding yeast strains were subsequently co-cultured with the CL-producing *E. coli* strain (ECL; Table S2, S3) (23).

Surprisingly, in the microbial consortium harboring PpMAX1c (ECL/YSL1c; Table S3), two new peaks (Fig. 2B) were detected in addition to the synthesis of CLA and 18-OH-CLA. The peak with retention time at 11.49 min was then confirmed to be strigol through comparison with the authentic standard, showing identical tandem mass spectrometry (MS/MS) fragmentation patterns (Fig. S4). The other peak with the retention time at 10.12 min and a positive mass/charge ratio (m/z^+^) = 347.1, which agrees with the mass of strigol, OB, or an oxygenated 5DS/4DO (Fig. 2B). Comparison with the available authentic standards, the unknown compound is excluded to be OB, strigol, or sorgomol (Fig. 2B). The unknown peak was then analyzed by high resolution mass spectrometry (HRMS) and MS/MS (Fig. S5). The appearance of the fragment m/z^+^=97 in MS/MS implies the presence of butenolide ring, and together with the molecular weight confirmed the identity of the unknown peak a SL compound (SL-1). Due to the lack of commercially available standard and the relatively low efficiency of the microbial bioproduction platform (at the level of ~92μg/L), we are currently not able to provide further elucidation of the structure of SL-1. CLA and 18-OH-CLA were detected from ECL/YSL1c (Table S3) but not 5DS (Fig. 2B). Thus, PpMAX1c likely first catalyzes four-step of oxidations to convert CL into 18-OH-CLA, which serves as the branch point and is oxidized to either strigol or the unknown SL-1(Fig. 1, Fig. S2). The titer of strigol in the consortium (ECL/YSL1c; Table S3) was 71.82±6.93 μg/L.

The other two MAX1 homologs encoded by peach, PpMAX1a and PpMAX1b, did not exhibit such activity as PpMAX1c. While only CLA was detected in the PpMAX1a-harboring microbial consortium (ECL/YSL1a; Fig. 2, Table S2, S3), PpMAX1b (ECL/YSL1b; Table S2, S3) synthesized both CLA and a trace amount of a new peak with m/z^-^ = 347.1 that agreed with the mass of a hydroxylated CLA. Likewise, due to the relatively low efficiency of the SL-producing microbial consortium, the identity of this compound remained to be identified and referred as SL-2 here (Fig. 2). To further examine the activities of PpMAX1a and PpMAX1b, the co-expression of PpMAX1a, PpMAX1b with PpMAX1c was also examined in the microbial consortium (ECL/YSL2a-c; Table S3), but no additional changes to the metabolite profiles were detected (Fig. S6).

### Identification of PpMAX1c-catalyzed strigol biosynthesis

Although 5DS was not detected from ECL/YSL1c, how strigol is synthesized by PpMAX1c from CL is unclear. To elucidate the biosynthetic route of strigol from CL by PpMAX1c, whole-cell biotransformation experiments were performed through incubating PpMAX1c-expressing yeast strain (YSL1c; Table S2, S3) with either the precursor CL or each of the possible intermediates (including CLA, 18-OH-CLA, and 5DS). Consistent with ECL/YSL1c co-culture assay, CL was converted into strigol and the unknown SL-1 (Fig. 3), with 18-OH-CLA detected as well (Fig. S7C). Likewise, a similar product profile was observed when using CLA and 18-OH-CLA as the substrate (Fig. 3, Fig. S7A) In comparison to the ECL/YSL1c co-culture assay, the whole-cell biotransformation experiments seem to be of lower efficiency, which is likely due to the instability of CL, CLA, and 18-OH-CLA (Fig. 3, Fig. S7B) (4, 30, 31). On the other hand, when YSL1c was incubated with the commercially available synthetic (±) 5DS, no strigol but only a peak of the same retention time and m/z^+^ signal as the unknown SL-1 was detected (Fig. 3, Fig. S7A). Further HR-MS/MS analysis was conducted to confirm that the identity of the 5DS-converted metabolite and the unknown SL-1 synthesized from ECL/YSL1c co-culture assay are the same compound (Fig. S5E, F, G). To examine whether SL-1 was the precursor to strigol, we also incubated YSL1c with SL-1 (prepared from PpMAX1c^C117S^ mutant, see *Material and Methods*), and no conversion from SL-1 to strigol was detected (Fig. S7D, E). Thus, strigol is unlikely to be converted from SL-1; and strigol and SL-1 are likely synthesized from 18-OH-CLA through independent routes catalyzed by PpMAX1c. According to previous investigations, strigol may be synthesized through two routes: 1) strigol is oxidized from intermediate 5DS; 2) BC ring of strigol is formed from 4,18-dihydroxy-CLA via 18-OH-CLA (Fig. S2). Since PpMAX1c cannot convert 5DS to strigol, likely strigol is synthesized from 4,18-dihydroxy-CLA bypassing 5DS though 4,18-dihydroxy-CLA cannot be detected in either ECL/YSL1c co-culture assay or the whole-cell biotransformation experiments, which may be due to the instability and low level of 4,18-dihydroxy-CLA. Additionally, we also examined the function of PpMAX1c with orobanchol-type SLs, 4DO and OB, using whole-cell biotransformation experiments. No conversions of 4DO or OB were detected (Fig. S7D, E), indicating that PpMAX1c is stereospecific towards the substrates. In sum, the whole-cell biotransformation experiments suggest that the BC ring closure of strigol likely occurs after the A-ring hydroxylation of 18-OH-CLA (Fig. 3A, Fig. S2), which is different from previously proposed strigol biosynthetic pathways in cotton (Fig. S1) (16, 22).

**Fig. 3.**
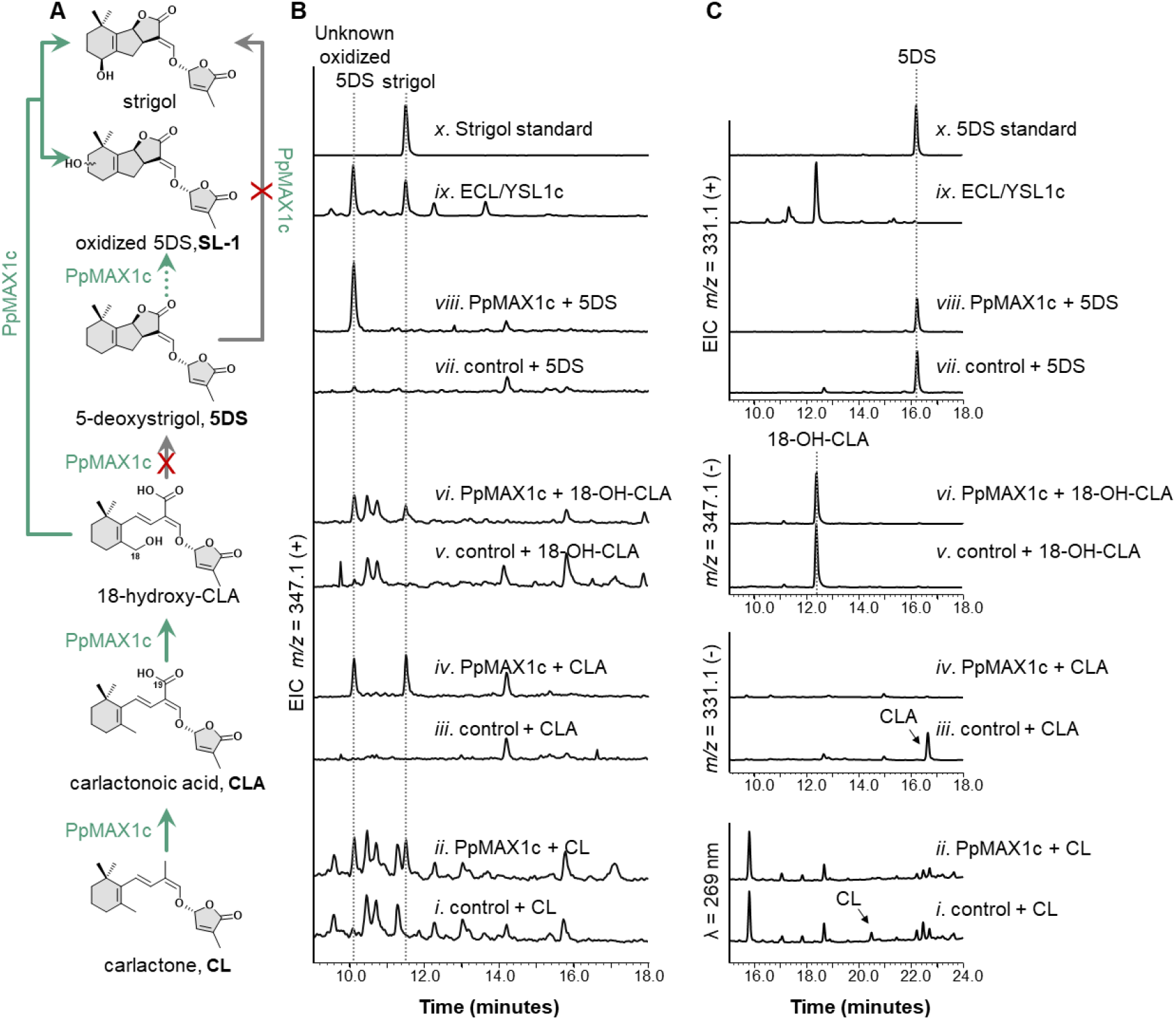
Whole-cell Biotransformation assays of PpMAX1c fed with CL, CLA, 18-OH-CLA, and 5DS as the substrate. (A) The proposed biosynthetic pathway of strigol and SL-1 in peach. (B) SIM EIC at strigol’s characteristic m/z^+^ = 347.1 of yeast harboring (i) an empty vector fed with CL (control), (ii) PpMAX1c fed with CL, (iii) an empty vector fed with CLA (control), (iv) PpMAX1c fed with CLA, (v) an empty vector fed with 18-OH-CLA (control), (vi) PpMAX1c fed with 18-OH-CLA, (vii) an empty vector fed with 5DS (control), (viii) PpMAX1c fed with 5DS; (ix) ECL/YSL1c; and (x) strigol standard. (C) High-performance liquid chromatography (HPLC) analysis at λ = 269 nm of yeast harboring (i) an empty vector fed with CL (control), and (ii) PpMAX1c fed with CL. SIM EIC at CLA’s characteristic m/z^-^ = 331.1 of yeast strain harboring (iii) an empty vector fed with CLA (control); and (iv) PpMAX1c fed with CLA. SIM EIC at 18-OH-CLA’s characteristic m/z^-^ =347.1 of yeast strain harboring (v) an empty vector fed with 18-OH-CLA (control); and (vi) PpMAX1c fed with 18-OH-CLA. SIM EIC at 5DS ’s characteristic m/z^+^ = 331.1 of yeast strain harboring (vii) an empty vector fed with 5DS; (viii) PpMAX1c fed with 5DS; (ix) ECL/YSL1c; and (x) 5DS standard. The characteristic m/z^+^ of strigol signal (*MW* = 346.38) is [C_19_H_22_O_6_ + H]^+^ = [C_19_H_23_O_6_]^+^ = 347.1. Data are representative of at least three biological replicates.

### Site-directed mutagenesis of PpMAX1c to study the strigol formation

To further investigate the biosynthetic route of strigol catalyzed by PpMAX1c and to understand the sequence-function correlation of MAX1, we conducted site-directed mutagenesis of PpMAX1c based on the multiple sequence alignment analysis of PpMAX1c with other MAX1 homologs (Fig. 4A, Fig. S8, Table S4). The sequence alignment analysis of MAX1s revealed that the heme-binding (FxxGxRxCxG) motif is conserved (32) (Fig. S8). To confirm the importance of the heme-binding motif, we made mutant PpMAX1c^G480A^, the catalytic activity of which was dramatically diminished with no strigol and SL-1 detected (ECL/YSL3; Table S3, Fig. 4). This confirmed that the conserved heme-binding motif is essential for MAX1 activity. In addition, we observed a number of residues that are only conserved within subclade group III (Fig. S8), which might be responsible for the unique catalytic activity of this subclade. To investigate the sequence-function correlation of subclade III MAX1s, we constructed PpMAX1c mutants on these residues and made PpMAX1c mutants including PpMAX1c^P402M^, PpMAX1c^C117S^. Most mutants showed no change in the catalytic activity when introduced to the microbial consortium (ECL/YSL3; Table S3, Fig. S9) except PpMAX1c^P402M^ and PpMAX1c^C117S^.The microbial consortium harboring PpMAX1c^P402M^ (ECL/YSL3M26; Table S2, S3) abolished the synthesis of all the products downstream of CLA (including 18-OH-CLA, strigol, and SL-1) (Fig. 4), suggesting that Pro402 is likely involved in the C18-hydroxylation. Interestingly, microbial consortium harboring PpMAX1c^C117S^ exhibited a dramatic decrease in the synthesis of strigol yet no obvious change in the synthesis of SL-1 (ECL/YSL3M4; Table S2, S3, Fig. 4). These results indicated that Cys117 is crucial for strigol formation but not the synthesis of SL-1, and further implied that the conversions of SL-1 and strigol from 18-OH-CLA are two independent routes. The activity of PpMAX1c^C117S^ may also provide a convenient strategy to alter SL profiles in *Prunus* in the future.

**Fig. 4.**
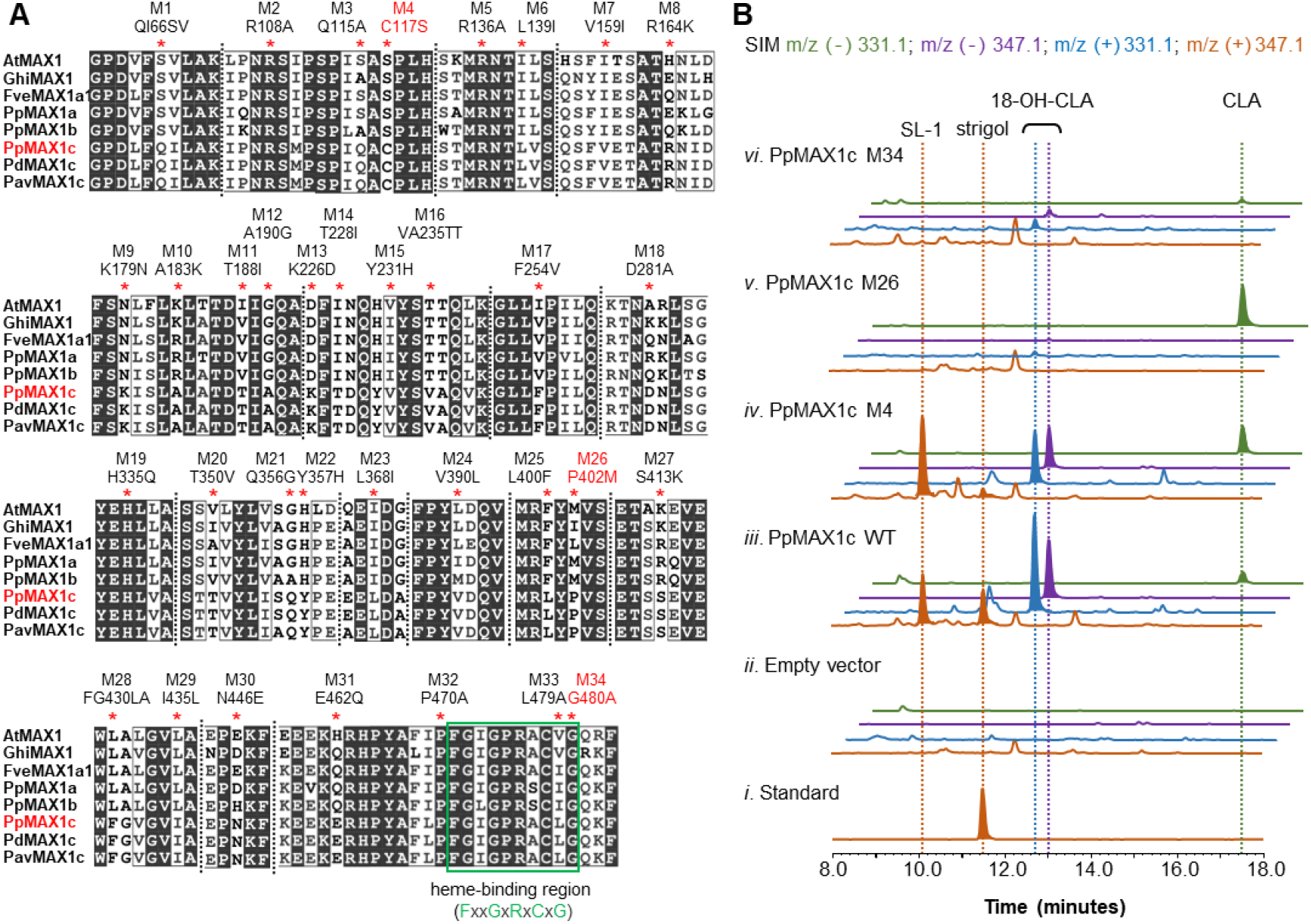
LC-MS analysis of products catalyzed by PpMAX1c mutants using SL-producing microbial consortium. (A) Multiple alignment of PpMAX1c with other MAX1 proteins. Genebank accession numbers can be found in Table S5. Amino acid sequence alignment was performed by using online software Clustal Omega with the default parameters. (https://www.ebi.ac.uk/Tools/msa/clustalo/). The mutated amino acid residues in this study have been marked with red asterisks. The heme-binding region (FxxGxRxCxG) are annotated in green box. (B) SIM EIC at m/z^-^ = 331.1 (green), m/z^-^ = 347.1 (purple), m/z^+^ = 331.1 (blue), and m/z^+^ = 347.1 (orange) of (i) strigol standard; CL-producing *E. coli* co-cultured with yeast expressing ATR1 and (ii) Empty vector, (iii) wild-type PpMAX1c, (iv) PpMAX1c Mutant 4 (C117S), (v) PpMAX1c Mutant 26 (P402M), and (vi) PpMAX1c Mutant 34 (G480A). Strain used for analysis: PpMAX1c (ECL/YSL1c); PpMAX1c Mutant 4 (C117S) (ECL/YSL3M4); PpMAX1c Mutant 26 (P402M) (ECL/YSL3M26); PpMAX1c Mutant 34 (G480A) (ECL/YSL3M34); Table S3. Data are representative of at least three biological replicates.

### PpMAX1c-mediated strigol biosynthesis is conserved and unique in *Prunus*

MAX1 genes are widely present in various plant species and play an important role in the diversity of SL (11, 18, 25–27). The discovery of PpMAX1c-mediated strigol biosynthesis further implies the functional diversity of MAX1s across different plants. All the *Prunus* plants included in this study contain three MAX1 (CYP711A) genes that span the three subclades Group I-III (Fig. 2B). A second member from group III subclade, PavMAX1c (*Prunus avium*, close relative of *P. persica*) of 98.69 % similarity to PpMAX1c, was assayed using the microbial consortium (ECL/YSL4, Table S3). PavMAX1c exhibited the same function as PpMAX1c and synthesized both SL-1 and strigol with the presence of CLA and 18-OH-CLA (Fig. S10). We also tested the functions of the MAX1 homologs from woodland strawberry (FveMAX1a1, FveMAX1a2, both of Group I), cotton (GhiMAX1), banana (MaMAX1a and MaMAX1b), apple (MdMAX1a1 and MdMAX1a2 of Group I, MdMAX1b1 and MdMAX1b2 of Group II), sunflowers (*Helianthus annuus;* HaMAX1) and ancestral angiosperm Amborella (*Amborella trichopoda*; AmtMAX1) (ECL/YSL5-11, Table S3, Fig. 2A). Likewise, FveMAX1a1, FveMAX1a2, GhiMAX1, MaMAX1a, MdMAX1a1, MdMAX1a2, HaMAX1 and AmtMAX1 exhibited the same function as PpMAX1a and AtMAX1, i.e. oxidation of CL to CLA; HaMAX1 can also produce a small amount of 18-OH-CLA; MdMAX1b1 from group II, same as PpMAX1b, oxidized CL to the unidentified hydroxylated CLA SL-2 likely *via* CLA (Table S1, Fig. S10). The functional identifications of these MAX1 homologs across the phylogenetic tree implies that likely the function of PpMAX1c as strigol synthase is conserved across Group III; the function of PpMAX1b synthesizing the unknown hydroxylated CLA SL-2 is also conserved across Group II; and MAX1s in Group I are likely CLA synthase.

### *In planta* identification of PpMAX1c as a strigol synthase

To validate PpMAX1c as a strigol synthase as detected in the microbial consortium, we need to confirm the activity of PpMAX1c in plant. *Sorghum bicolor* homologs of *D27 (SbD27), CCD7 (SbCCD7*), and *CCD8 (SbCCD8*) were used to produce CL by *Agrobacterium tumefaciens-mediated* infiltration in *N. benthamiana* leaves as reported previously (Fig. 5) (27). *PpMAX1a, PpMAX1b* and *PpMAX1c* were transiently co-expressed with *SbD27, SbCCD7, SbCCD8*, and a *S. bicolor* NADPH-P450 reductase (*SbCPR1*) in *N. benthamiana*. CL was decreased in leaves co-expressing *PpMAX1a*, *PpMAX1b* or *PpMAX1c*. As expected, strigol was detected in *PpMAX1c*-expressed leaves but not in *PpMAX1a*- and *PpMAX1b*-expressed leaves (Fig. 5). In addition, strigone (33), an oxidized metabolite of strigol, was detected, while the peak of 5DS was a trace in *PpMAX1c*-expressed leaves but none of them in *PpMAX1a*- and *PpMAX1b*-expressed leaves. These results validate the function of PpMAX1c as a strigol/strigone synthase synthesizing from CL *in planta*.

**Fig. 5.**
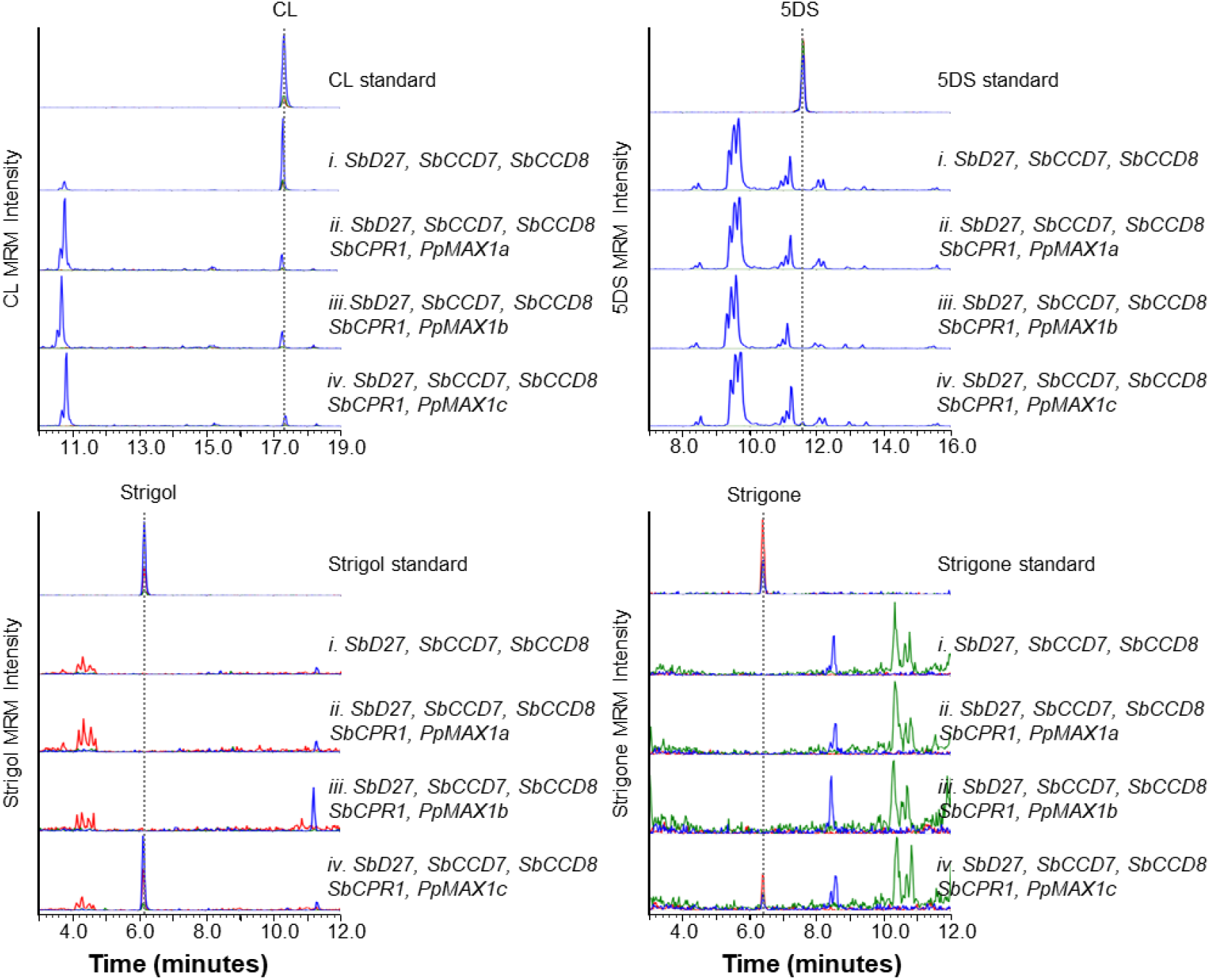
Functional identification of PpMAX1c as a strigol synthetase using *N. benthamiana* systems. Production of strigol in the *N. benthamiana* leaves co*-*expressing *SbD27, SbCCD7, SbCCD8, SbCPR1*, and *PpMAX1s*. (i) *SbD27+SbCCD7+SbCCD8;* (ii) *SbD27+SbCCD7+SbCCD8+SbCPR1+PpMAX1a;* (iii) *SbD27*+*SbCCD7*+*SbCCD8*+*SbCPR1*+*PpMAX1b*; (iv) *SbD27+SbCCD7+SbCCD8+SbCPR1+PpMAX1c*. Multiple reaction monitoring (MRM) chromatograms of CL (blue, 303.00/97.00; red, 303.00/189.00; green, 303.00/207.00; *m/z* in positive mode), 5DS (blue, 331.15/97.00; red, 331.15/216.00; green, 331.15/234.00; *m/z* in positive mode), strigol (blue, 329.00/215.00; red, 329.00/97.00; green, 347.00/215.00; *m/z* in positive mode) and strigone (blue, 345.00/203; red, 345.00/231.00; green, 345.00/97.00; *m/z* in positive mode) by LC-MS/MS are shown. All activities assays are representative of at least three biological replicates. Abbreviations: Sb, *Sorghum bicolor;* Pp, *Prunus persica*.

### Identification of PpCYP722C as Group I CYP722C responsible for OB biosynthesis

In addition to multiple MAX1 homologs, the *P. persica* genome also encodes one CYP722C gene (PpCYP722C) that is conserved in the *Prunus* family. Phylogenetic analysis of CYP722Cs including PpCYP722C indicates that PpCYP722C is a Group I CYP722C that generally converts CLA to OB (Fig. S11) (23), which implies that the *Prunus* family potentially can synthesize both strigol and OB. To test such a hypothesis, we introduced AtMAX1 and PpCYP722C into *S. cerevisiae* to yield strain YSL13 and co-cultured with CL-producing *E. coli* strain (ECL/YSL13; Table S2, S3). As expected, OB can be detected from ECL/YSL13 (Fig. 6A).

**Fig. 6.**
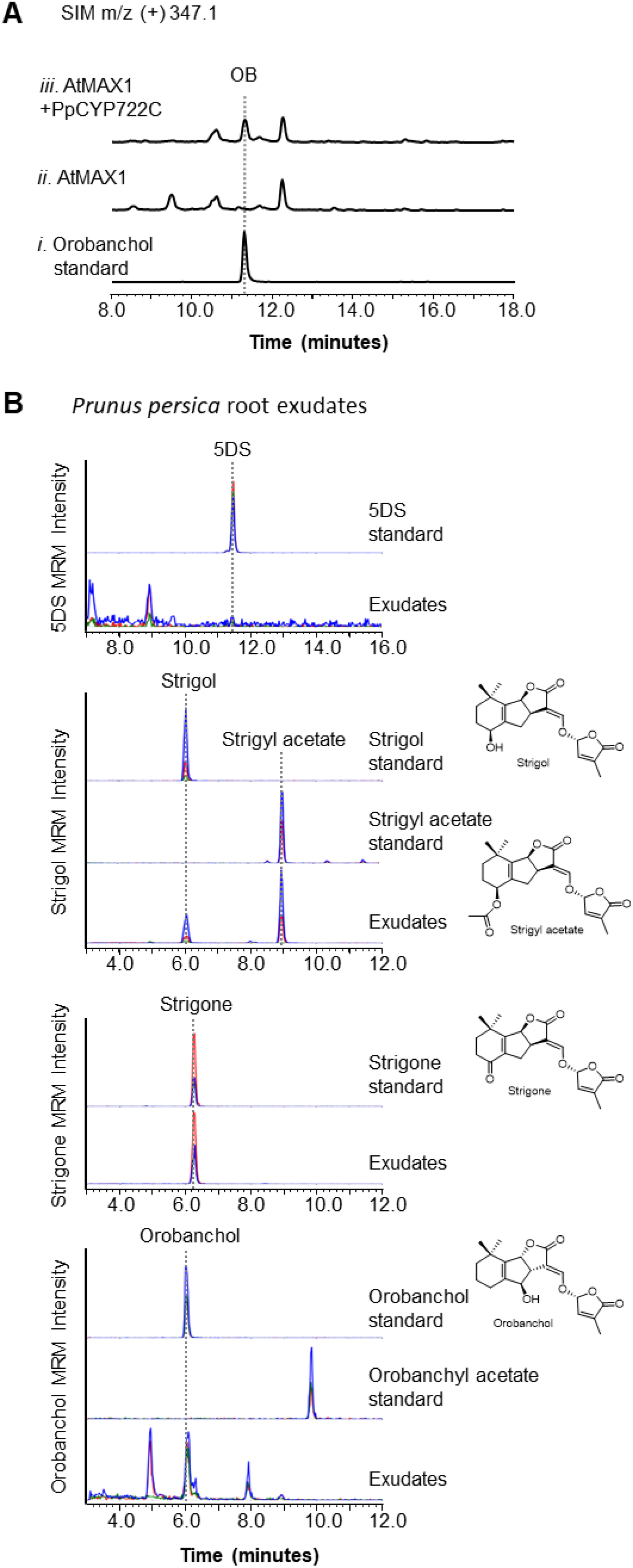
SL analysis in root exudates of *Prunus persica*. (A) Functional verification of CYP722C from *Prunus persica* as OB synthase. SIM EIC at OB’s characteristic m/z^+^ =347.1 of (i) OB standard; CL-producing *E. coli* co-cultured with yeast expressing ATR1, AtMAX1 and (ii) Empty vector (ECL/YSL13N; Table S2, S3); (iii) PpCYP722C (ECL/YSL13; Table S2, S3). (B) Detection of strigol, strigyl acetate, strigone and orobanchol in root exudates of *P. persica*. Multiple reaction monitoring (MRM) chromatograms of 5DS (blue, 331.15/97.00; red, 331.15/216.00; green, 331.15/234.00; *m/z* in positive mode), strigol/strigyl acetate (blue, 329.00/215.00; red, 329.00/97.00; green, 347.00/215.00; *m/z* in positive mode), strigone (blue, 345.00/203; red, 345.00/231.00; green, 345.00/97.00; *m/z* in positive mode) and orobanchol/orobanchyl acatate (blue, 347.00/233; red, 347.00/97.00; green, 347.00/205.00; *m/z* in positive mode) by LC-MS/MS are shown. Data are representative of three biological replicates.

### *P. persica* can produce both strigol and orobanchol

The functional identification of PpMAX1c and PpCYP722C indicates that *P. persica* has the catalytic capability to produce both strigol and OB, which is unusual and has not been observed to our knowledge. Thus, it is worthwhile to investigate the SL profiles in *P. persica*. We then examined the SL profiles of the seedlings of *P. persica*. LC-MS analysis confirmed the presence of strigol, OB, as well as strigol derivatives strigone and putative strigyl acetate, but not 5DS or 4DO (Fig. 6B). The relative amount of strigone/strigol in *P. persica* seedlings compared to *in vivo* conversion using *N. benthamiana* implies that strigone might be synthesized from strigol by other endogenous oxidase(s) in plants. One acetyltransferase is likely responsible for the conversion of strigol to strigyl acetate in *P. persica* and remains to be uncovered.

## Discussion

Different SL structures have been reported to exhibit distinct biological effects, and plants may evolve to produce more than one SLs to cope with complicated environments (5). However, much remains to be investigated on the SL structures that can be produced from different plants and the biosynthesis, the physiological functions of different SL structures in different plants, and the origin and evolution of SL biosynthesis and function. In this study, using the recently established SL-producing microbial consortium (23), we conducted functional screening of an array of MAX1 analogs, especially those in the rose family. A strigol synthase (PpMAX1c) was discovered, and the catalytic activity of PpMAX1c synthesizing strigol directly from CL bypassing 5DS was confirmed through substrate feed experiments and mutant analysis. The biosynthetic pathway of strigol was subsequently reconstituted and validated in *N. benthamiana* using transient expression. Meanwhile, the function of PpCYP722C as an OB synthase was confirmed, and the presence of both OB and strigol in peach was confirmed. This work serves as a good example of metabolite prediction based on gene function identification, which highlights the importance of deciphering the sequence-function correlation of plant biosynthetic enzymes to more accurately predicate plant metabolites without metabolic analysis.

Recent studies indicate the important role of CYP711As in the structural diversity of SLs (17, 18, 27). Although most plants encode only one copy of MAX1 gene with highly conserved function as CLA synthase (11), the presence of multiple copies of MAX1 orthologous genes is not an unusual phenomenon, especially in monocot plants (17, 27, 29), dicotyledonous *Leguminosae*, and *Rosaceae*. The activities of OsCYP711A2 (Os900/OsMAX1b in Fig. 1, *Os01g0700900*) in converting CL to 4DO (17), OsCYP711A3 (Os1400/OsMAX1c in Fig. 1, *Os01g07014000*) in converting 4DO to OB (17), one of four MAX1 homologs in sorghum SbMAX1a in converting CL to 18-OH-CLA (27, 29), and PpMAX1c characterized in this study as a strigol synthase are all unique functions that are only conserved in certain species or genus (Fig. 2A). To further investigate the functional diversity, more attention should be placed on the genus that encode multiple copies of MAX1 analogs.

Meanwhile, the MAX1 encoded by the primitive liverwort *Marchantia paleacea* (MpaMAX1) was very recently characterized to catalyze the conversion of CL to bryosymbiol (BSB, Fig. 1), a non-canonical SL (25). Unlike hydroxylation that is prototypically catalyzed by MAX1 in seed plants (11), MpaMAX1 catalyzes the epoxidation of the C7-C8 double bond, which facilitates a subsequent C-ring closure to afford the synthesis of the unique structure of BSB (25), the function of which remains to be investigated. The unique function of MpaMAX1 also implies the importance of functional investigations on MAX1s encoded by primitive land plants to uncover the diversity and evolution of the CYP711A family.

The PpMAX1c-mediated strigol biosynthesis seems to be only conserved in *Prunus* genus. 5DS was barely detected in the strigol-producing microbial consortium (ECL/YSL1c), *N. benthamiana* agroinfiltration-based assay expressing PpMAX1c, or in peach seedlings. In addition, 5DS was only converted to SL-1 but not strigol in the PpMAX1c whole cell transformation assay. All the results explicitly exclude the possibility that strigol is converted from 5DS in *Prunus*. By contrast, both strigol and 5DS were detected in cotton (16). In this study, we have also tested the activity of MAX1 from the Mexican cotton *Gossypium hirsutum* (GhiMAX1), which exhibited only CLA-producing activity (Fig. S10). Previous investigations confirmed the function of cotton CYP722C (GaCYP722C from *Gossypium arboreum*) as a 5DS synthase that function on CLA (18, 23). In addition, strigol can be efficiently converted from fed rac-5DS in cotton (16). All these results strongly suggest a distinct strigol biosynthetic pathway in cotton from 5DS synthesized by GaCYP722C. Recently, CYP728B35 in sorghum was found to oxidize 5DS to afford the synthesis of sorgomol (34). The missing enzyme in cotton catalyzing the conversion of strigol from 5DS is likely another cytochrome P450. The distinct biosynthetic pathways of strigol in peach and cotton that are catalyzed by a different set of cytochrome P450s (CYP711 in peach; CYP722 and unknown enzyme in cotton) suggests that 5DS and strigol may have emerged at the same time through divergent evolution from the occurrence of CLA catalyzed by CYP711s (Fig. S2). Although previous feeding experiments showed that CL or CLA can be converted to strigol without the detection of 5DS, and 5DS cannot be converted to strigol in moonseed (16), which is consistent with what we observed with PpMAX1c biotransformation assay; a putative hydroxyl-CL was also detected in moonseed when feeding CL (16) (Fig. S1), and whether there is a third mechanism to synthesize strigol from CL is thus inclusive and remains to be investigated (Fig. S1).

Interestingly, PpMAX1c can catalyze multistep oxidation at no less than three different carbon positions on CL: 3-step C19-oxidation from methyl group to carboxyl group, C4-hydroxylation, C18-hydroxylation (to afford synthesis of strigol), and one oxidation at unknown carbon positions (to afford synthesis of SL-1, Fig. 1). OsCYP711A2 can catalyze multistep oxidation at two different carbon positions: 3-step C19-oxidation from methyl group to carboxyl group and C18-hydroxylation (to afford synthesis of 4DO) (11, 17). Similarly, the recently characterized SbMAX1a also catalyzes multi-step oxidations at two different carbon positions: 3-step C19-oxidation from methyl group to carboxyl group and C18-hydroxylation (to afford synthesis of 18-OH-CLA, Fig. 1), and presumably further oxidation of C18-hydroxy (to afford synthesis of 18-oxo-CLA and OB) (27, 29). In addition, SbMAX1c, PpMAX1b, and MdMAX1b1 were found to synthesize unknown monohydroxylated-CLAs (structures to be elucidated), which also requires at least four-step oxidations at two different carbon positions (29). Such robust catalytic activity (multi-step oxidation at multiple carbons) is not commonly observed among plant cytochrome P450s involved in secondary metabolism but seems to be quite common among MAX1 analogs. Likely these MAX1 analogs retained the CLA-producing (C19-carboxylation) activity from the ancestral MAX1s and acquired additional catalytic activities during subsequent evolution. The driven force of evolution for the synthesis of these SLs (e.g., strigol, various hydroxylated-CLAs) by various MAX1 analogs in different plants should be correlated with the functions of the corresponding SLs in plant. Recently, various hydroxylated CLs and CLAs (including 3-HO-CL, 4-HO-CL, 16-HO-CL; 3-HO-CLA, 4-HO-CLA, 16-HO-CLA) were also identified from *Arabidopsis* (35), which brings up the questions on the physiological functions of these noncanonical SLs. It is also to be noted that *Arabidopsis* only encodes one MAX1 gene, and thus additional oxidoreductases such as cytochrome P450s are likely required for the synthesis of the different hydroxylated CLs and CLAs in *Arabidopsis*.

Unlike OsCYP711A2, which was previously identified to synthesize 4DO, an orobanchol-type canonical SL (17), PpMAX1c produced strigol-type SL from CL. OsCYP711A2 and PpMAX1c both belong to the CYP711A subfamily, and thus it raises an interesting question on how CYP711As control the stereospecificity of BC rings, which might be resolved through biochemical investigations aided with the structural comparison between OsCYP711A2 and PpMAX1c. In addition to strigol synthesis, a few other previously unknown activities of MAX1s (e.g., SbMAX1c, PpMAX1b, and MdMAX1b1) were also uncovered, but due to the low efficiency of the SL-producing microbial consortium, the structures of the unknown SLs cannot be elucidated, which also highlights the necessity of metabolic engineering to enhance the efficiency of this bioproduction platform as a more useful and versatile characterization tool.

Furthermore, despite years of studies, much remains to be investigated to understand the structure-function correlation of SLs. Most plants have been reported to produce a mixture of SL structures instead of a single SL structure (4–6), but why one plant needs to produce multiple SLs is unknown. Several plants have been reported to produce both strigol- and orobanchol-type SLs, but such phenomena are cultivar-, growth condition-, and development stage-dependent (5). Here, we have shown the production of both strigol and orobanchol in *Prunus*, and characterized the biosynthetic machineries that affords the synthesis of both types of SLs. The *Prunus* genus thus serves as a unique model to identify the function of different SLs.

In conclusion, we have identified PpMAX1c as a unique strigol synthase that converts CL into strigol and an unknown oxidized 5DS using microbial consortium-based assay, whole cell transformation assay fed with different substrates, *N. benthamiana* agroinfiltration-based pathway reconstitution, and mutagenesis analysis. This work highlights the high efficiency of using a SL-producing microbial consortium to study SL biosynthesis, demonstrates the functional diversity of MAX1 analogs, and validates *Prunus* as a unique model for future investigation on the structure-function correlation of SLs.

## Materials and Methods

### Reagents and general procedures

(±)5-deoxy-strigol (purity >98%), (±)strigol and (±)-orobanchol were purchased from Strigolab (Torino, Italy), (±)4-deoxyorobanchol [(±)-2’-epi-5-deoxystrigol] was purchased from Chempep (Wellington, FL, United States), the other chemicals were purchased from Sigma-Aldrich or Fisher Scientific. Sorgomol was obtained from Dr. Kaori Yoneyama (Ehime University, Japan) as a gift. The gateway entry vector pDONR221, BP clonase enzyme and LR recombinase were purchased from Invitrogen (Rockville, MD, United States). The *Saccharomyces cerevisiae* Advanced Gateway Destination Vector Kit was purchased from Addgene (Watertown, MA, United States). Phusion high-fidelity DNA polymerase (New England Biolabs) was used for PCR amplification. High-resolution mass (HRMS) analysis was performed on a Synapt G2-Si quadrupole time-of-flight mass spectrometer (Waters, Milford, MA, United States) coupled to an I-class ultra-performance liquid chromatography (UPLC) system (Waters, Milford, MA, United States). Liquid chromatography–mass (LC-MS) analysis was performed on a Shimadzu LC-MS 2020 (Kyoto, Japan) with Optima LC-MS grade solvent (Fisher Scientific). Liquid chromatography-tandem mass spectrometry (LC-MS/MS) analysis was performed on a triple quadrupole/linear ion trap instrument (QTRAP5500; AB Sciex, MA, USA) coupled to a UPLC (Nexera X2; Shimazu, Japan). The synthetic genes were synthesized by Integrated DNA Technologies (Coralville, IA, United States) and Oligonucleotide primers were purchased from Life Technologies (Pleasanton, CA, United States). Sanger sequencing was performed at Azenta US, Inc. (South Plainfield, NJ). The plasmids used in this study are listed in Table S2. The gene sequences and GenBank accession numbers of the proteins were obtained from Tables S7 and S5, respectively.

### Plant material and growth conditions

*P. persica* (ornamental peaches, cv. Yaguchi) seeds were collected from a peach flower garden (Koga, Ibaraki, Japan) (36). *N. benthamiana* seeds were propagated in our laboratory (11).

### Plasmid construction and yeast transformation

The full-length synthetic CYP genes (such as PpMAX1c) were codon optimized for *Saccharomyces cerevisiae* and introduced into the entry vector (pDONR221) (Invitrogen, Rockville, MD, United States) by Gateway BP reactions. Subsequently, they were cloned into yeast expression vectors (such as pAG416GPD-ccdB) by Gateway LR reactions (Resulting pAG416GPD-PpMAX1c). Then these constructs were co-transformed with ATR1 (NADPH-CYP reductase 1 from *A. thaliana*) expression vector (Addgene, Catalog # 178288) into the *Saccharomyces cerevisiae* wild-type strain CEN.PK2-1D using the Frozen-EZ yeast transformation II kit (Zymo Research) according to according to the manufacturer’s instructions.

### Phylogenetic analysis of MAX1 or CYP722C

Phylogenetic trees was conducted in MEGA X program using neighbor-joining method (90% partial deletion, 5000 bootstraps, p-distance mode) (37). GenBank accession numbers are summarized in the Tables S5 and S6.

### Site-directed mutagenesis

To generate the PpMAX1c mutant, the plasmid pENTR-PpMAX1c (pYL859) was employed as a template and was amplified using phusion high-fidelity DNA polymerase (New England Biolabs, Ipswich, MA, USA). The PCR products was purified, recovered and used to perform Gibson assembly reaction, then the reaction mixture was directly transformed into TOP 10 competent cells. The correctly sequenced plasmids were cloned into yeast expression vectors pAG416GPD-ccdB, resulting pAG416GPD-PpMAX1c mutant, which were used for activity testing. The primers are listed in Table S8. The underlined bases are mutations introduced by PCR. In vivo functional validation of the PpMAX1c mutant was carried out as described above.

### Production of SL by engineered *E. coli-yeast* consortium

The protocol for fermentation is the same as described before with some optimization (23). XY medium was employed for coculture fermentations, XY medium [13.3 g l^-1^ monopotassium phosphate (KH_2_PO_4_), 4 g l^-1^ diammonium phosphate [(NH_4_)_2_HPO_4_], 1.7 g l^-1^ citric acid, 0.0025 g l^-1^ cobalt(II) chloride (CoCl_2_), 0.015 g l^-1^ manganese(II) chloride (MnCl_2_), 0.0015 g l^-1^ copper(II) chloride (CuCl_2_), 0.003 g l^-1^ boric acid (H_3_BO_3_), 0.0025 g l^-1^ sodium molybdate (Na_2_MoO_4_), 0.008 g l^-1^ zinc acetate [Zn(CH_3_COO)_2_], 0.03 g l^-1^ iron(III) citrate, 0.03 g l^-1^ ferrous sulphate, 0.0045 g l^-1^ thiamine, 1.3 g l^-1^ magnesium sulfate (MgSO_4_), 5 g l^-1^ yeast extract, and 40 g l^-1^ xylose, pH 7.0] was prepared as described previously (23). For *E. coli* inducible expression, a yellow single colony of the *E. coli* BL21(DE3) harboring pAC-BETAipi (Addgene # 53277), pCDFDuet-trAtCCD7-OsD27 (Addgene # 178285) and pET21a-trAtCCD8 (Addgene # 178286) (generating strain ECL, Table S3) was cultured at 37 °C overnight in 3 ml of lysogeny broth (LB) medium with ampicillin (100 μg ml^-1^), spectinomycin (50 μg ml^-1^) and chloramphenicol (34 μg ml^-1^). 100 μl of the overnight grown seed culture was grown at 37 °C and 220 r.p.m. When OD_600_ ≈0.6, after cooling, isopropyl β-D-1-thiogalactopyranoside (IPTG) (final concentration of 0.2 mM) and ferrous sulphate (10 mg l^-1^) was added to the culture. After IPTG induction, the strain were cultured at 18°C and 220 r.p.m. for another 16 h. For yeast ectopic expression, a single colony of yeast (*S. cerevisiae*) cells were cultured overnight at 30 °C and 220 r.p.m in 2 ml synthetic drop-out (SD) medium [0.425 g yeast nitrogen base (YNB) (BD Biosciences, San Jose, CA, United States), 1.25 g ammonium sulfate [(NH_4_)_2_SO_4_], 25 ml glucose solution (200 g l^-1^), 25 ml amino acid drop-out mix (20 g l^-1^)]. 100 μl of overnight grown yeast seed culture were inoculated into 5 ml SD medium in a 50 ml Erlenmeyer flask and grown at 30°C with shaking at 220 r.p.m for 15 h. Next day, the bacteria and yeast cells were harvested by centrifugation at 3,500 r.p.m. for 3 min and re-suspended together in 5 ml XY media and grown at 25°C for 48 h.

### SL analysis from *E. coli-yeast* consortium

SL was extracted and detected as described previously (23). For the extraction of extracellular metabolites. 5 ml cultures were centrifuged at 3500 r.p.m. for 10 min. The medium was extracted with 4 ml ethyl acetate (EtOAc). For better separation of the organic and aqueous phases, the solution were centrifuged at 4000 r.p.m. for 20 min. Then the EtOAc layer was transferred to a new 1.7 ml microcentrifuge tube and concentrated in vacuo (Eppendorf concentrator plus, Enfield, CT, United States). The residues was redissolved in 120 μL acetone, centrifuged at 13,000 r.p.m. for 10 min and then 15 μl of sample was subjected to LC-MS analysis (Shimadzu, LC-MS 2020). LC-MS parameters: C18 column (Kinetex® C18, 100 mm × 2.1 mm, 100 Å, particle size 2.6 μm; Phenomex, Torrance, CA, United States); flow rate of 0.4 ml/min; column temperature 40 °C; mobile phase A: water containing 0.1% (v/v) formic acid; mobile phase B: acetonitrile containing 0.1% (v/v) formic acid. 0–28 min, 5–100% B; 28–35 min, 100% B; 35–40 min, 5% B.

### *S. cerevisiae-based* whole-cell biotransformation

The yeast strains expressing ATR1 with PpMAX1c (YSL1c, Table S3), or harboring an empty plasmid (negative control, YSL1N, Table S3) were used for the whole-cell biotransformation. Taking YSL1c as an example, 1mL SD medium is first inoculated with a fresh colony and cultured overnight at 30°C and 200 r.p.m. 20 μL of the overnight culture was used to inoculate 1 mL fresh SD medium in a test tube and grown at 30°C with shaking at 220 r.p.m for 16 h. The cells were then harvested by centrifugation at 3000 r.p.m. and resuspended in 1 ml YNB medium. 5DS, 4DO, OB, or strigol were fed to the 1mL YSL1c biotransformation matrix at a final concentration of 0.08 mg/L, incubated at 25 °C and 220 r.p.m. for 12 hours. On the other hand, CL, CLA, 18-OH-CLA, and the oxidized 5DS SL-1 were extracted from ECL/1D, ECL/YSL13N’, ECL/YSL12 and ECL/YSL3M4, respectively (Table S2, S3), using similar method as described above in *SL analysis from E. coli-yeast consortium*. Then, Bond Elut DEA cartridge column was utilized to separate each SL mixture into acidic fraction (containing CLA and/or 18-OH-CLA) and neutral fraction (containing CL and/or SL-1) to enrich the corresponding target SLs (27). The semi-purified SLs were then dissolved in 10 μl of acetone, added into the 1mL YSL1c biotransformation matrix, and incubated at 25 °C and 220 r.p.m. for 12 hours. Next, 1 ml EtOAc was added into each biotransformation matrix, vortexed vigorously, and applied to centrifugation. The EtOAc phase was then collected and concentrated in vacuo. The residue was redissolved in 50 μl of acetone. The solution was centrifuged at 13,000 r.p.m. for 10 min and then 15 μl samples were subjected to LC-MS analysis as described above.

### HR-MS analysis of strigolactones

Before HR-LC–MS/MS, the target components were purified by HPLC (Shimadzu, LC-MS 2020). HPLC parameters: C18 column (Kinetex® C18, 100 mm × 2.1 mm, 100 Å, particle size 2.6 μm; Phenomex, Torrance, CA, United States); flow rate of 0.4 ml/min; column temperature 40 °C; mobile phase A: water; mobile phase B: acetonitrile. The eluting program is: 0–12 min, 5–45.7% B; 12–16 min, 45.7-100% B; 16–18 min, 100% B; 18–18.5 min, 100-5% B. 18.5–20 min, 5% B. The eluates (SL-1 elutes at about 10 min) were concentrated and subjected to LC–MS/MS system as described below.

HR-MS analysis of strigolactones were performed on an I-class UPLC system (Waters) coupled to a Synapt G2-Si Q-TOF MS (Waters). The separation was conducted on the C18 column (Kinetex C18, 100 mm by 2.1 mm, 100 Å, particle size 2.6 μm; Phenomex, Torrance, CA, USA). Separation Method: the mobile phase A was 0.1% (v/v) formic acid in water, the mobile phase B was 0.1% (v/v) formic acid in acetonitrile, and gradient elution was 0.4 ml/min. The gradient was as follows: 0 to 18 min, 5 to 100 %B; 18 to 20 min, 100 %B; 20 to 23 min, 100% to 5% B; and 23 to 25 min, 5 %B. Total Run Time was 25.00 min; Column temperature was 40°C; The injection volume was 5 μl. The MS were obtained using the positive ion mode, the scan range of m/z was from 50 to 1200 with a 0.2 s scan time. The MS/MS conditions were as follows: collision energy was 25 eV. Source temperature was 150°C and desolvation temperature was 600°C. Capillary voltage was 1 KV; precursor ion m/z, 331.15 for 5DS, 347.15 for orobanchol, strigol and SL-1. Desolvation gas flow: 1100 liter/hour. All gases were nitrogen except the collision gas. The analysis of the data was performed using MassLynx 4.1 software (Waters).

### Heterologous reconstitution of strigol biosynthesis in *N. benthamiana*

The full-length synthetic *PpMAX1a, PpMAX1b* and *PpMAX1c* genes were subcloned into the *SalI* site of pBYR2HS (38) using GeneArt Seamless Cloning and Assembly Enzyme Mix (Invitrogen). The plasmids of *SbD27, SbCCD7, SbCCD8*, and *SbCPR1* cloned previously in pBYR2HS (27) were used in this study. Co-expression by *Agrobacterium tumefaciens* (GV3101)-mediated infiltration was performed as reported previously (27). To prevent necrosis, sodium ascorbate solution (200 mM) was treated to the infiltrated leaves using a spray (39). After 5 days, 1 g of agro-infiltrated leaves were cut into 1-cm squares and extracted in 40 ml of acetone. The filtrates were evaporated to aqueous residues and added 1 ml of distilled water. The aqueous residues were extracted twice with 2 ml of EtOAc. The EtOAc phase was purified using Bond Elut DEA cartridge column (100 mg; Agilent Technologies, USA) and analyzed by LC-MS/MS (QTRAP 5500; AB Sciex, USA) as reported previously (11, 24, 27).

### Metabolite analysis of peach seedlings

10 seedlings of *P. persica* were grown in pots filled with a mold for 9 months, then transplanted in 10 cm-diameter pots filled with vermiculite and further grown using tap water for 1 month in a glasshouse. 300 ml of water was poured and collected from the bottom of pot. The collected water was extracted with EtOAc. The EtOAc phase was dried over anhydrous sodium sulfate and evaporated to dryness. SLs were analyzed by LC-MS/MS.

## Acknowledgments

We thank Prof. Kaori Yoneyama (Ehime University, Japan) for her generous gift of sorgomol. We thank Prof. David Nelson (University of California, Riverside) and Prof. Kang Zhou (National University of Singapore) for the helpful discussion. We thank the Metabolomics Core Facility at UC Riverside and Dr. Amancio De Souza for instrument access, training, and data analysis. T. Chiu for valuable feedback in the preparation of the manuscript. pAC-BETAipi was a gift from F. X. Cunningham Jr. (Addgene plasmid no. 53277; http://n2t.net/addgene:53277; RRID: Addgene_53277). This work is supported by the Cancer Research Coordinating Committee Research Award (grant to Y.L., CRN-20-634571), NSF CAREER Award (grant to Y.L., 2144626), and JSPS Grant-in-Aid for Scientific Research (grant numbers 19K05838 and 22H02269 to T.N.).

## Author contributions

S.W. and Y.L. conceived of the project; S.W., A.Z., T. N., and Y.L. designed the experiments; S.W., A.Z., and T. N. performed the experiments and analyzed the results; K.H. and K.M. performed a transient expression; A.Y. and X.X. analyzed SLs *in planta;* K.Y. prepared plant materials; S.W., T. N., and Y.L. wrote the manuscript.

## Additional information

Y.L. and S.W. are inventors on a provisional patent application related to this work filed by University of California, Riverside (U.S. Provisional Application No. 63/142,801, filed 28 January 2021, patent pending). The authors declare that they have no other competing interests.

