## Supplementary material for "Identification of a *Prunus* MAX1 Homolog as a Unique Strigol Synthase from Carlactone Bypassing 5-Deoxystrigol": SI

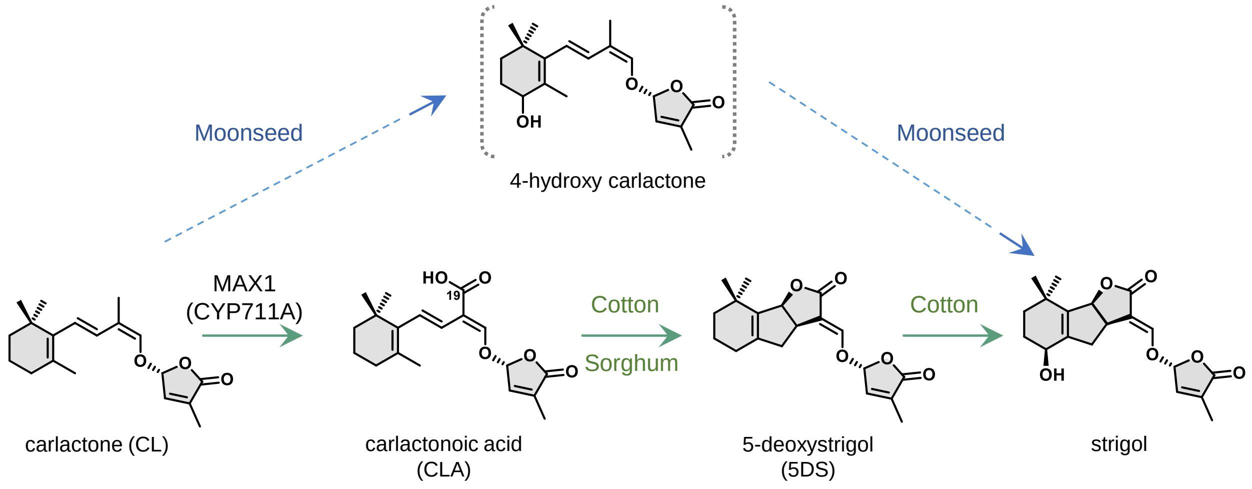


**Fig. S1**: **The proposed biosynthetic pathways of strigol in cotton and moonseed.** Moonseed can convert CL into strigol bypassing the 5DS likely via unidentified intermediate 4-hydroxy carlactone, and cannot oxidize 5DS to strigol, which suggests that 5DS is not a precursor to strigol in moonseed (1). On the other hand, cotton can oxidize 5DS to afford the synthesis of strigol and thus the synthesis of strigol is proposed to be derived from 5DS in cotton (1). The enzymes involved in strigol biosynthesis for both plants have not been identified.

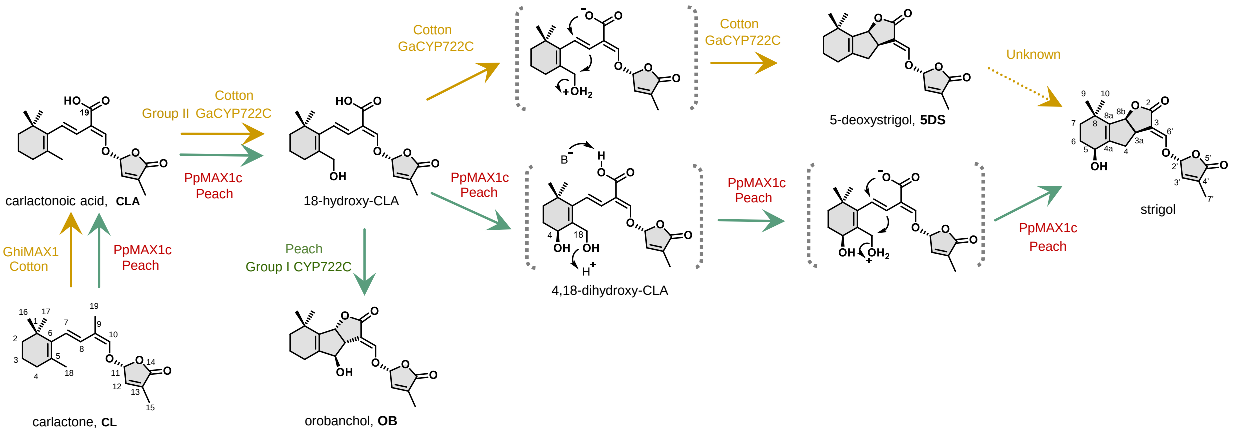


**Fig. S2**: **Proposed enzymic mechanism for the biosynthesis of strigol catalyzed by PpMAX1c.** In peach, PpMAX1c can catalyze the multistep oxidations at three different positions, i.e., the direct oxidation from methyl group to carboxyl group at C19 and the introduction of hydroxyl group at C4 and C18 positions respectively to afford the synthesis of intermediate 4,18-dihydroxy-CLA. The C19 carboxyl group then deprotonates to form a nucleophile and attacks C7, followed by nucleophilic substitution between C8 and C18 with water as the leaving group to afford the BC ring closure. In contrast, strigol in cotton is likely oxidized from 5DS by an unidentified mechanism, while the synthesis of 5DS from CL is well characterized and catalyzed by GhiMAX1 (CL to CLA) and GaCYP722C (CLA to 5DS). At least three enzymes are involved in the conversion of CL to strigol in cotton.

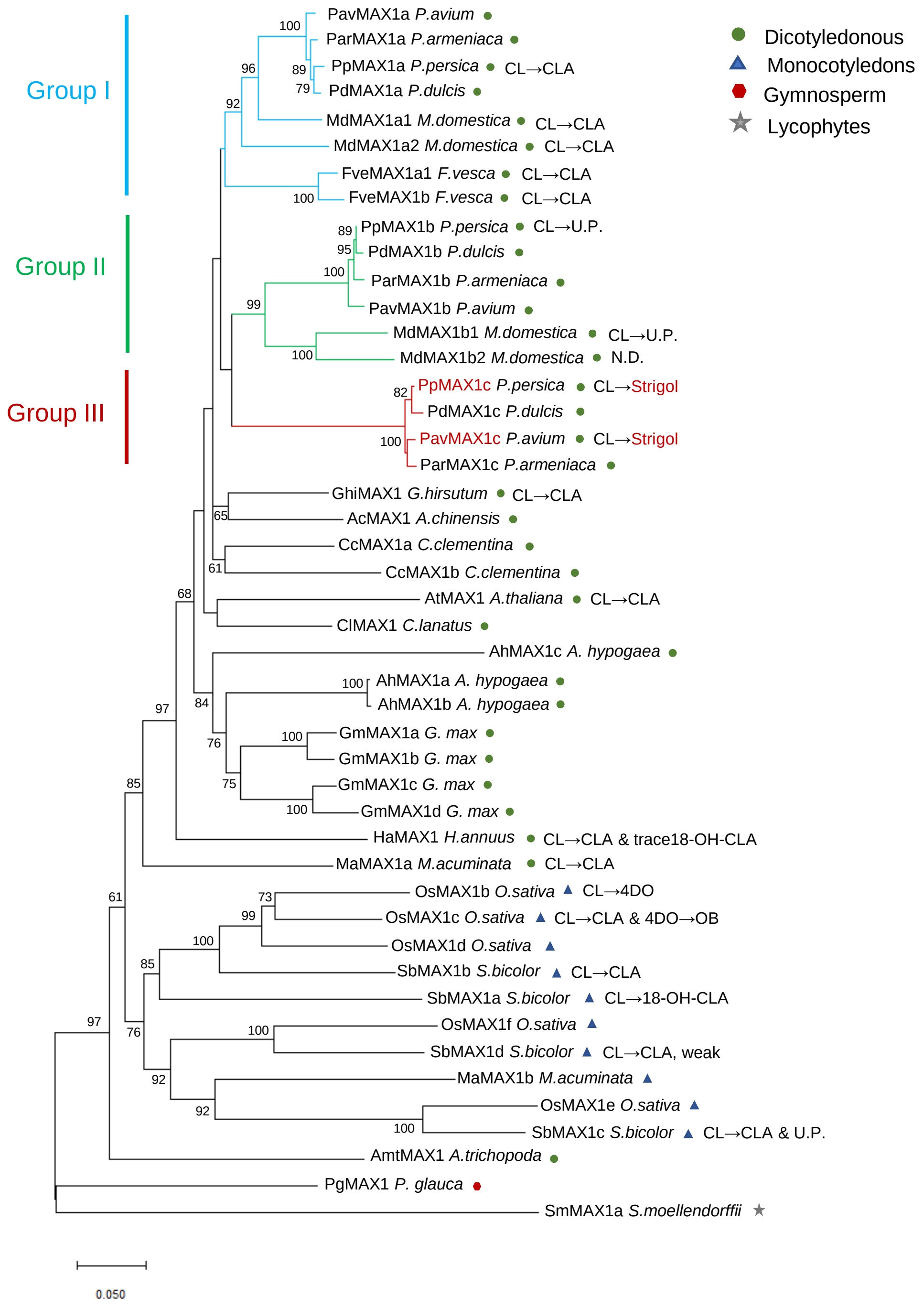


**Fig. S3**: **Extended phylogenetic analysis of MAX1 (CYP711A) genes.** The phylogenetic analysis was conducted in MEGA X using neighbor-joining method (90% partial deletion, 5000 replicates, p-distance mode). A total of 46 amino acid sequences are included and the Genebank accession numbers can be found in Table S5. Groups I, II and III are distinguished by blue, green and red, respectively.

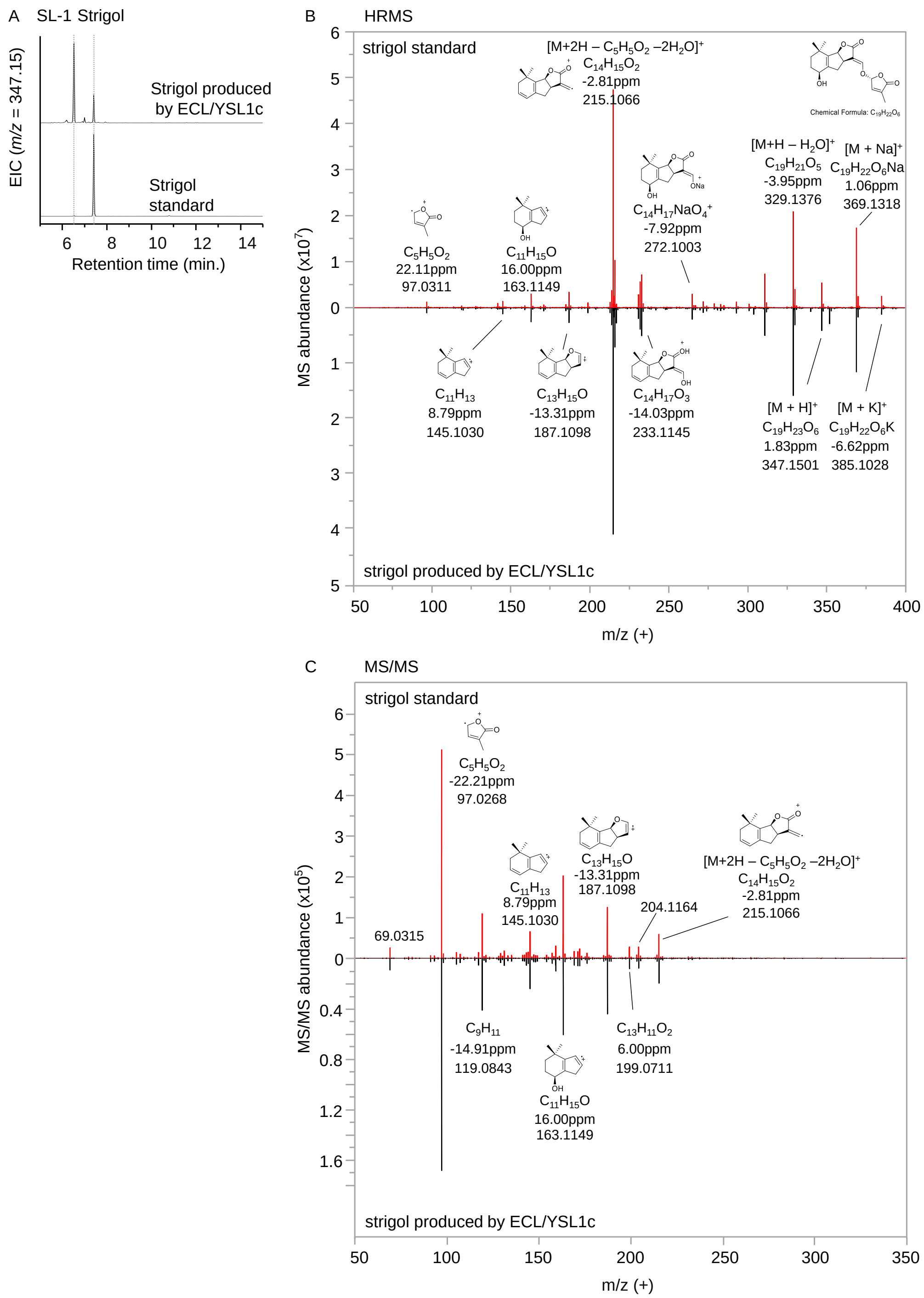


**Fig. S4**: **HRMS analysis of strigol in PpMAX1c assay.** (A) The HR-LC-MS chromatogram of strigol produced by strigol-producing consortia ECL/YSL1c (Table S3) and strigol standard. (B) The HRMS spectrum of strigol produced by ECL/YSL1c and strigol standard. HRMS analysis revealed $[M+H]^+$ of m/z 347.1501, corresponding to a chemical formula of $C_{19}H_{22}O_6$. (C) Tandem mass spectrometry (MS/MS) fragmentation patterns of strigol produced by ECL/YSL1c and strigol standard. They had the identical MS/MS fragmentation patterns, which confirms the identity of the compound produced by ECL/YSL1c strigol. HRMS data was recorded in positive ionization mode. The full-scan range is 50-1200 m/z.

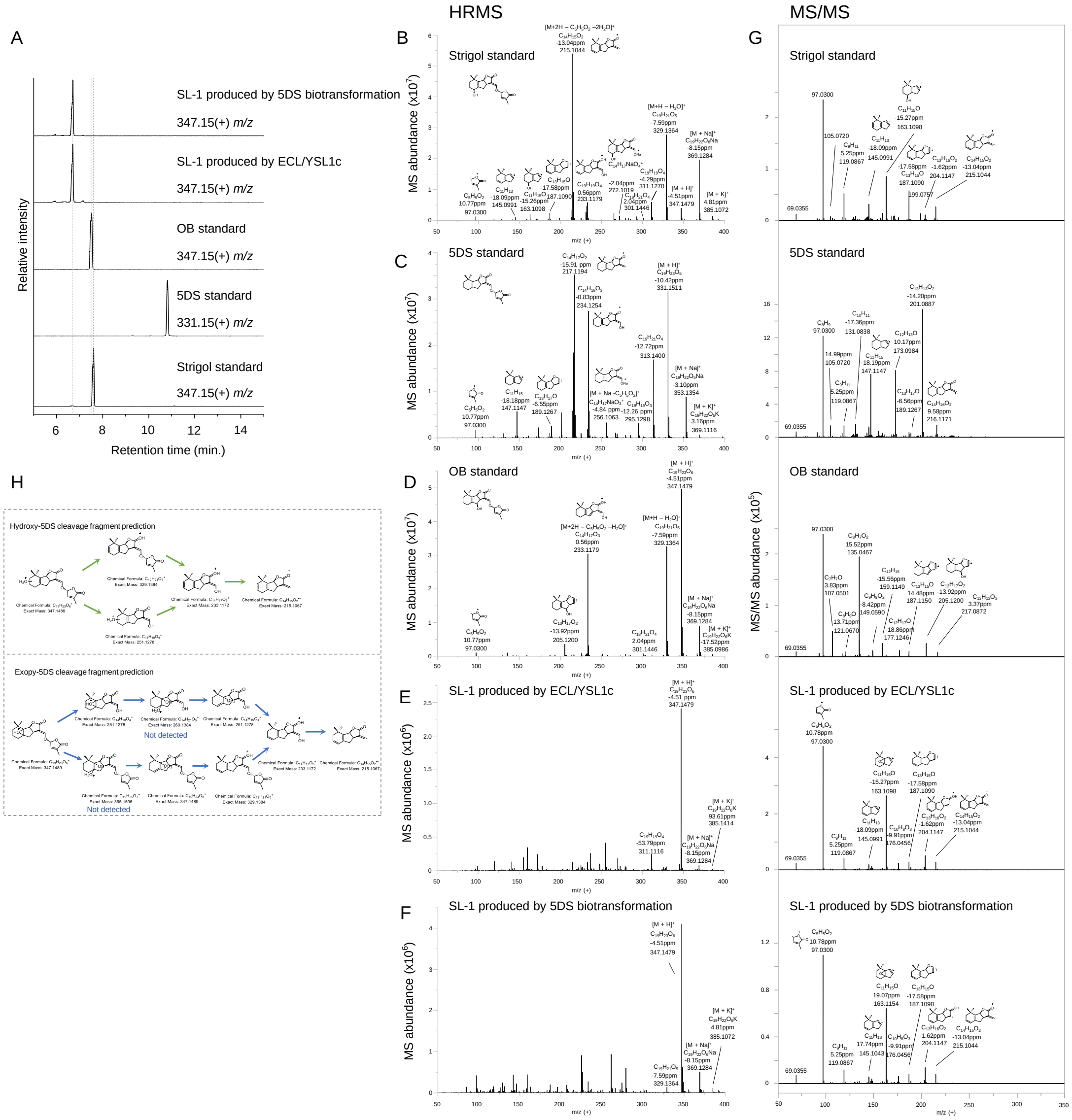


**Fig. S5**: **HRMS analysis of SL-1 produced by ECL/YSL1c.** (A) The HR-LC-MS chromatogram of OB, 5DS, and strigol standards; and SL-1 produced by ECL/YSL1c (Table S3), and PpMAX1c whole cell transformation fed with 5DS. The HRMS spectrum of (B) strigol standard, (C) 5DS standard, and (D) OB standard. (E) The HRMS spectrum of SL-1 produced by ECL/YSL1c (Table S3). (F) The HRMS spectrum of SL-1 produced by PpMAX1c whole cell transformation fed with 5DS. HRMS analysis revealed $[M+H]^+$ of m/z 347.1479, corresponding to a chemical formula of $C_{19}H_{22}O_6$. (G) The comparison of the MS/MS fragmentation patterns of OB, 5DS and strigol standard; and SL-1 produced by ECL/YSL1c, and PpMAX1c whole cell transformation fed with 5DS. MS/MS analysis shows that the compound produced from PpMAX1c whole cell transformation fed with 5DS is identical to SL-1 synthesized from ECL/YSL1c. HRMS data were recorded in positive ionization mode, the full-scan range is 50-1200 m/z. (H) We speculate that unknown oxidized 5DS (SL-1) may be epoxidized 5DS or hydroxylated 5DS (less likely due to the retention time). We predicted the fragmentation pattern of hydroxylated 5DS and epoxidized 5DS respectively, and most of the predicted fragmentation signals are consistent with signals observed in the HRMS and MS/MS analysis.

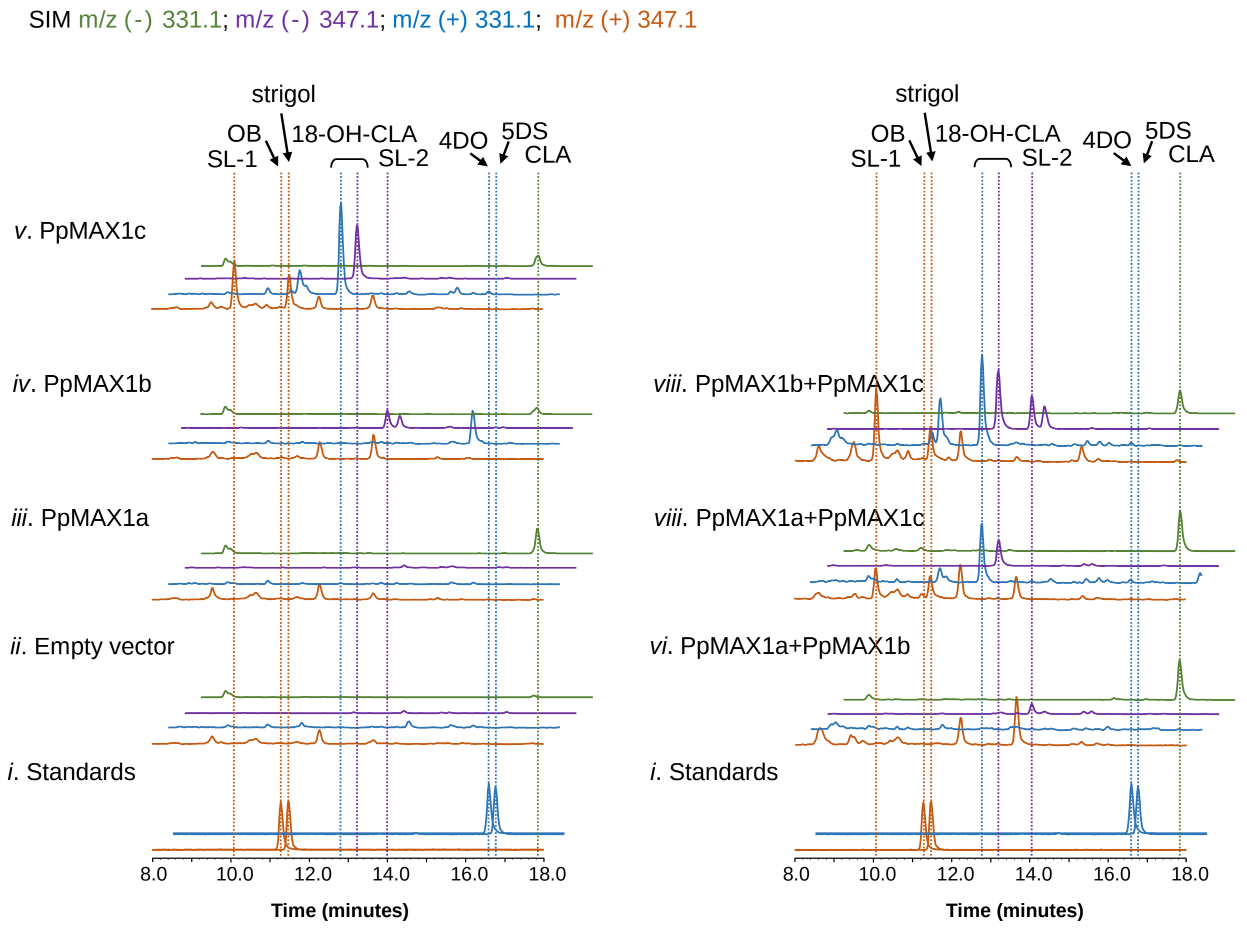


**Fig. S6**: **Introduction of PpMAX1c to PpMAX1a- or PpMAX1b-harboring consortium did not change the product profile.** Selected ion monitoring SIM (SIM) extracted ion chromatogram (EIC) of $m/z^- = 331.1$ (green), $m/z^- = 347.1$ (purple), $m/z^+ = 331.1$ (blue), and $m/z^+ = 347.1$ (orange) of (i) Strigol, 4DO and 5DS standard; CL-producing *E. coli* co-cultured with yeast expressing ATR1 and (ii) Empty vector (ECL/YSL1N), (iii) PpMAX1a (ECL/YSL1a), (iv) PpMAX1b (ECL/YSL1b), (v) PpMAX1c (ECL/YSL1c), (vi) PpMAX1a+PpMAX1b (ECL/YSL2a), (vii) PpMAX1a+PpMAX1c (ECL/YSL2b), and (viii) PpMAX1b+PpMAX1c (ECL/YSL2c). Detailed information of each strain can be found in Table S3. The data is representative of at least three biological replicates.

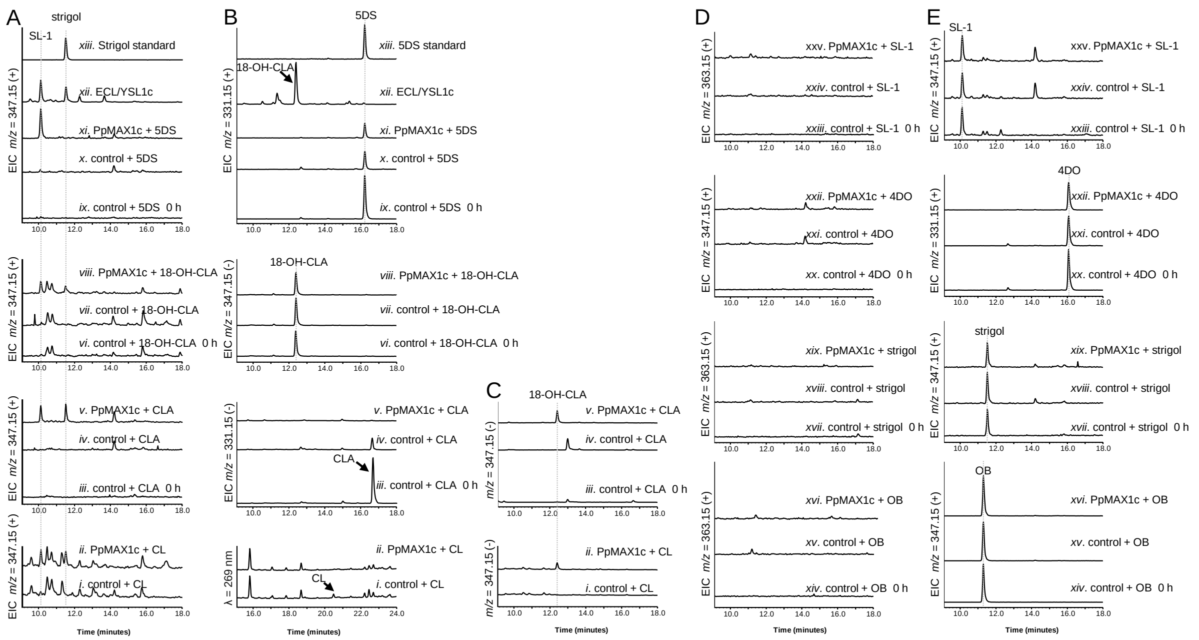


**Fig. S7**: **Whole-cell Biotransformation assays of PpMAX1c fed with CL, CLA, 18-OH-CLA, 4DO, 5DS, OB, or strigol as the substrate.** LC-MS analysis of PpMAX1c whole-cell biotransformation assays. Heterologously expressed PpMAX1c was incubated with substrate for 0 h or 12 h. SIM EIC using strigol′s characteristic $m/z^+$ = 347.1 of whole cell assay of yeast harboring (i) an empty vector fed with CL and quenched at 12h; (ii) PpMAX1c fed with CL and quenched at 12h; (iii) an empty vector fed with CLA and quenched at 0 h; (iv) an empty vector fed with CLA and quenched at 12h; (v) PpMAX1c fed with CLA and quenched at 12h; (vi) an empty vector fed with 18-OH-CLA and quenched at 0 h; (vii) an empty vector fed with 18-OH-CLA and quenched at 12h; (viii) PpMAX1c fed with 18-OH-CLA and quenched at 12h; (ix) an empty vector fed with 5DS and quenched at 0 h; (x) an empty vector fed with 5DS and quenched at 12h; (xi) PpMAX1c fed with 5DS and quenched at 12h; (xii) ECL/YSL1c; and (xiii) strigol standard. (B) High-performance liquid chromatography (HPLC) analysis at $\lambda$ = 269 nm of yeast harboring (i) an empty vector control fed with CL (control), and (ii) PpMAX1c fed with CL. SIM EIC at CLA′s characteristic $m/z^-$ = 331.1 of yeast strain harboring (iii) an empty vector fed with CLA and quenched at 0 h; (iv) an empty vector fed with CLA and quenched at 12h; (v) PpMAX1c fed with CLA and quenched at 12h. SIM EIC at 18-OH-CLA′s characteristic $m/z^-$ =347.1 of yeast strain harboring (vi) an empty vector fed with 18-OH-CLA and quenched at 0 h; (vii) an empty vector fed with 18-OH-CLA and quenched at 12h; (viii) PpMAX1c fed with 18-OH-CLA and quenched at 12h. SIM EIC at 5DS ′s characteristic $m/z^+$ = 331.1 of yeast strain harboring (ix) an empty vector fed with 5DS and quenched at 0 h; (x) an empty vector fed with 5DS and quenched at 12h; (xi) PpMAX1c fed with 5DS and quenched at 12h; (xii) ECL/YSL1c; and (xiii) 5DS standard. (C) SIM EIC at 18-OH-CLA′s characteristic $m/z^-$ =347.1 of yeast strain harboring (i) an empty vector fed with CL and quenched at 12h; (ii) PpMAX1c fed with CL and quenched at 12h; (iii) an empty vector fed with CLA and quenched at 0 h; (iv) an empty vector fed with CLA and quenched at 12h; (v) PpMAX1c fed with CLA and quenched at 12h; (D) SIM EIC at hydroxylated OB′s characteristic $m/z^-$ = 363.1 of yeast strain harboring (xiv) an empty vector fed with OB and quenched at 0 h; (xv) an empty vector fed with OB and quenched at 12 h; (xvi) PpMAX1c fed with OB and quenched at 12h. SIM EIC at hydroxylated strigol′s characteristic $m/z^-$ = 363.1 of yeast strain harboring (xvii) an empty vector fed with strigol and quenched at 0 h; (xviii) an empty vector fed with strigol and quenched at 12 h; (xix) PpMAX1c fed with strigol and quenched at 12h; SIM EIC at hydroxylated 4DO′s characteristic $m/z^-$ = 347.1 of yeast strain harboring (xx) an empty vector fed with 4DO and quenched at 0 h; (xxi) an empty vector fed with 4DO and quenched at 12 h; (xxii) PpMAX1c fed with 4DO and quenched at 12h. SIM EIC at hydroxylated SL-1′s characteristic $m/z^-$ = 363.1 of yeast strain harboring (xxiii) an empty vector fed with SL-1 and quenched at 0 h; (xxiv) an empty vector fed with SL-1 and quenched at 12 h; (xxv) PpMAX1c fed with SL-1 and quenched at 12h. (E) SIM EIC at OB′s characteristic $m/z^+$ = 347.1 of yeast strain harboring (xiv) an empty vector fed with OB and quenched at 0 h; (xv) an empty vector fed with OB and quenched at 12 h; (xvi) PpMAX1c fed with OB and quenched at 12h. SIM EIC at strigol′s characteristic $m/z^+$ = 347.1 of yeast strain harboring (xvii) an empty vector fed with strigol and quenched at 0 h; (xviii) an empty vector fed with strigol and quenched at 12 h; (xix) PpMAX1c fed with strigol and quenched at 12h; SIM EIC at 4DO ′s characteristic $m/z^+$ = 331.1 of yeast strain harboring (xx) an empty vector fed with 4DO and quenched at 0 h; (xxi) an empty vector fed with 4DO and quenched at 12 h; (xxii) an empty vector fed with 4DO and quenched at 12 h; SIM EIC at SL-1′s characteristic $m/z^+$ = 347.1 of yeast strain harboring (xxiii) an empty vector fed with SL-1 and quenched at 0 h; (xxiv) an empty vector fed with SL-1 and quenched at 12 h; (xxv) PpMAX1c fed with SL-1 and quenched at 12h. The characteristic $m/z^+$ of strigol signal (*MW* = 346.38) is $[C_{19}H_{22}O_6 + H]^+ = [C_{19}H_{23}O_6]^+$ = 347.1. The characteristic m/z+ signal of 4DO and 5DS (*MW* = 330.38) is $[C_{19}H_{22}O_5 + H]^+ = [C_{19}H_{23}O_5]^+$ = 331.1. All activities assays are representative of at least three biological replicates. Strigol and unknown oxidized 5DS (SL-1) can be converted from CL, CLA, or 18-OH-CLA. SL-1 can be converted from 5DS, but not strigol. Strigol cannot be converted from 5DS or SL-1. (*O*)-type 4DO and OB cannot be consumed by PpMAX1c, suggesting that PpMAX1c is stereospecific towards the substrates. Strigol cannot be consumed by PpMAX1c, suggesting that strigol is the final product of PpMAX1c. It is also noted that the substrates are not stable and were degraded after 12h. The data is representative of at least three biological replicates.

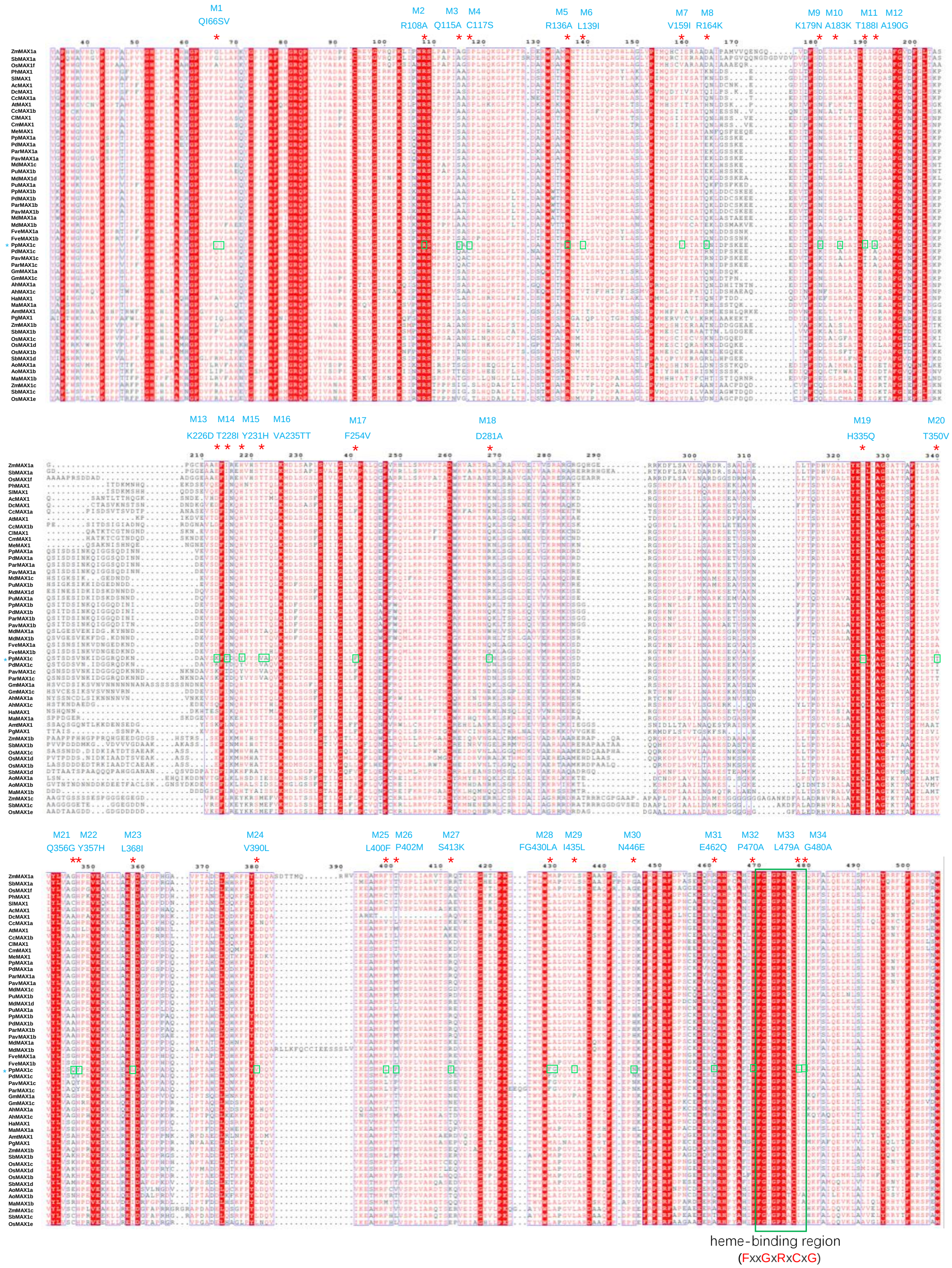


**Fig. S8**: **Multiple alignment of PpMAX1c with other MAX1 analogs.** Genebank accession numbers of amino acid sequences used here can be found in Table S5**.** Amino acid sequence alignment was performed by using online software Clustal Omega with the default parameters. (https://www.ebi.ac.uk/Tools/msa/clustalo/). The mutated amino acid residues in this study have been marked with blue asterisks. The heme-binding region (FxxGxRxCxG) is annotated in green box.

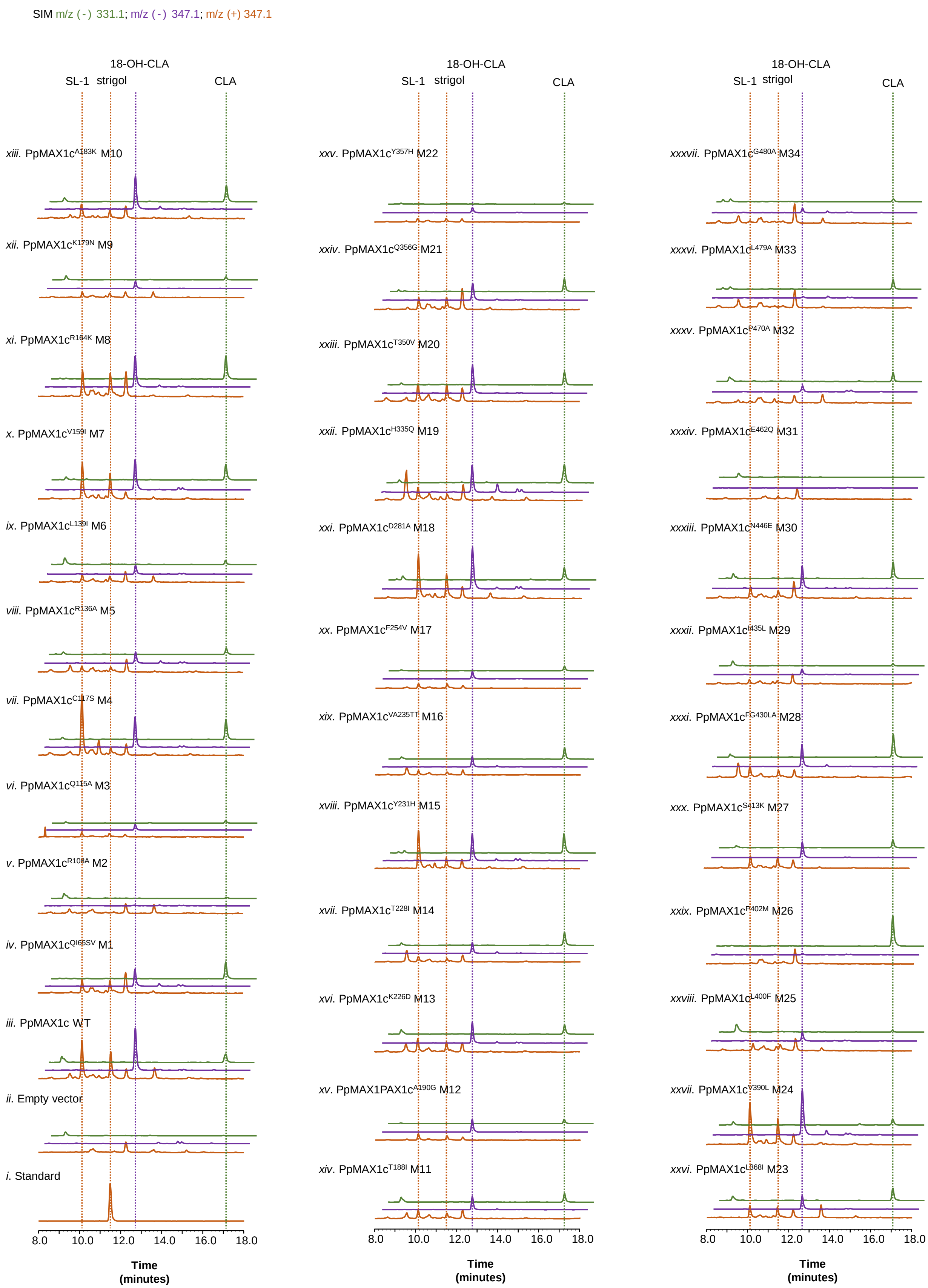
SIM m/z (-) 331.1; m/z (-) 347.1; m/z (+) 347.1
18-OH-CLA
SL-1
strigol
CLA
xiii. PpMAX1c$^{A183K}$ M10
xii. PpMAX1c$^{K179N}$ M9
xi. PpMAX1c$^{R164K}$ M8
x. PpMAX1c$^{V159I}$ M7
ix. PpMAX1c$^{L139I}$ M6
viii. PpMAX1c$^{R136A}$ M5
vii. PpMAX1c$^{C117S}$ M4
vi. PpMAX1c$^{Q115A}$ M3
v. PpMAX1c$^{R108A}$ M2
iv. PpMAX1c$^{Q66SV}$ M1
iii. PpMAX1c WT
ii. Empty vector
i. Standard
xxv. PpMAX1c$^{Y357H}$ M22
xxiv. PpMAX1c$^{Q356G}$ M21
xxiii. PpMAX1c$^{T350V}$ M20
xxii. PpMAX1c$^{H335Q}$ M19
xxi. PpMAX1c$^{D281A}$ M18
xx. PpMAX1c$^{F254V}$ M17
xix. PpMAX1c$^{VA235TT}$ M16
xviii. PpMAX1c$^{Y231H}$ M15
xvii. PpMAX1c$^{T228I}$ M14
xvi. PpMAX1c$^{K226D}$ M13
xv. PpMAX1PAX1c$^{A190G}$ M12
xiv. PpMAX1c$^{T188I}$ M11
xxxvii. PpMAX1c$^{G480A}$ M34
xxxvi. PpMAX1c$^{L479A}$ M33
xxxv. PpMAX1c$^{P470A}$ M32
xxxiv. PpMAX1c$^{E462Q}$ M31
xxxiii. PpMAX1c$^{N446E}$ M30
xxxii. PpMAX1c$^{I435L}$ M29
xxxi. PpMAX1c$^{FG430LA}$ M28
xxx. PpMAX1c$^{S413K}$ M27
xxix. PpMAX1c$^{F402M}$ M26
xxviii. PpMAX1c$^{L400F}$ M25
xxvii. PpMAX1c$^{V390L}$ M24
xxvi. PpMAX1c$^{L368I}$ M23
8.0
10.0
12.0
14.0
16.0
18.0
Time
(minutes)

**Fig. S9**: **LC-MS analysis of PpMAX1c mutants using SL-producing microbial consortium.** SIM EIC at $m/z^- = 331.1$ (green), $m/z^- = 347.1$ (purple), $m/z^+ = 347.1$ (orange) of (i) strigol standard; ECL (Table S3) cocultured with yeast expressing ATR1 and (ii) empty vector (YSL1N); (iii) PpMAX1c wild type (YSL1c); (iv) PpMAX1c$^{QI66S}$ (YSL3M1); (v) PpMAX1c$^{R108A}$ (YSL3M2); (vi) PpMAX1c$^{Q115A}$ (YSL3M3); (vii) PpMAX1c$^{C117S}$ (YSL3M4); (viii) PpMAX1c$^{R136A}$ (YSL3M5); (ix) PpMAX1c$^{L139I}$ (YSL3M6); (x) PpMAX1c$^{V159I}$ (YSL3M7); (xi) PpMAX1c$^{R164K}$ (YSL3M8); (xii) PpMAX1c$^{K179N}$ (YSL3M9); (xiii) PpMAX1c$^{A183K}$ (YSL3M10); (xiv) PpMAX1c$^{T188I}$ (YSL3M11); (xv) PpMAX1c$^{A190G}$ (YSL3M12); (xvi) PpMAX1c$^{K226D}$ (YSL3M13); (xvii) PpMAX1c$^{T228I}$ (YSL3M14); (xviii) PpMAX1c$^{Y231H}$ (YSL3M15); (xix) PpMAX1c$^{VA235TT}$ (YSL3M16); (xx) PpMAX1c$^{F254V}$ (YSL3M17); xxi) PpMAX1c$^{D281A}$ (YSL3M18); (xxii) PpMAX1c$^{H335Q}$ (YSL3M19); (xxiii) PpMAX1c$^{T350V}$ (YSL3M20); (xxiv) PpMAX1c$^{Q356G}$ (YSL3M21); (xxv) PpMAX1c$^{Y357H}$ (YSL3M22); (xxvi) PpMAX1c$^{L368I}$ (YSL3M23); (xxvii) PpMAX1c$^{V390L}$ (YSL3M24); (xxviii) PpMAX1c$^{L400F}$ (YSL3M25); (xxix) PpMAX1c$^{P402M}$ (YSL3M26); (xxx) PpMAX1c$^{S413K}$ (YSL3M27); (xxxi) PpMAX1c$^{FG430LA}$ (YSL3M28); (xxxii) PpMAX1c$^{I435L}$ (YSL3M29); (xxxiii) PpMAX1c$^{N446E}$ (YSL3M30); (xxxiv) PpMAX1c$^{E462Q}$ (YSL3M31); (xxxv) PpMAX1c$^{P470A}$ (YSL3M32); (xxxvi) PpMAX1c$^{L479A}$ (YSL3M33); (xxxvii) PpMAX1c$^{G480A}$ (YSL3M34). Detailed strain information are available in Table S3. PpMAX1c harboring M4 (C117S) mutation almost abolished the synthesis of strigol but showed no change in the synthesis of the oxidized 5DS (SL-1), indicating that this residue is specifically involved in the production of SL-1. PpMAX1c harboring M26 (P402M) mutation interrupts the production of downstream products of CLA (including 18-OH-CLA, strigol, and SL-1) and accumulates more CLA, suggesting that Pro402 is likely involved in the C18-hydroxylation. PpMAX1c harboring M2 (R108A), M31 (E462Q), M32 (P470A), M33 (L479A), or M34 (G480A) mutations exhibit significant decrease in the overall catalytic activity towards downstream SLs. These residues may be involved in substrate binding or reside in the catalytic site. While PpMAX1c harboring M1 (QI66SV), M3 (Q115A), M5 (R136A), M6 (L139I), M7 (V159I), M8 (R164K), M9 (K179N), M10 (A183K), M11 (S115C), M12 (A190G), M13 (K226D), M14 (T228I), M15 (Y231H), M16 (VA235TT), M17 (F254V), M18 (D281A), M19 (H335Q), M20 (T350V), M21 (Q356G), M22 (Y357H), M23 (L368I), M24 (V390L), M25 (L400F), M27 (S413K), M28 (FG430LA), M29 (I435L), or M30 (N446E) mutations showed no change in its catalytic activity, which implies that the corresponding residues are not likely involved in substrate binding or catalysis. The data is representative of at least three biological replicates.

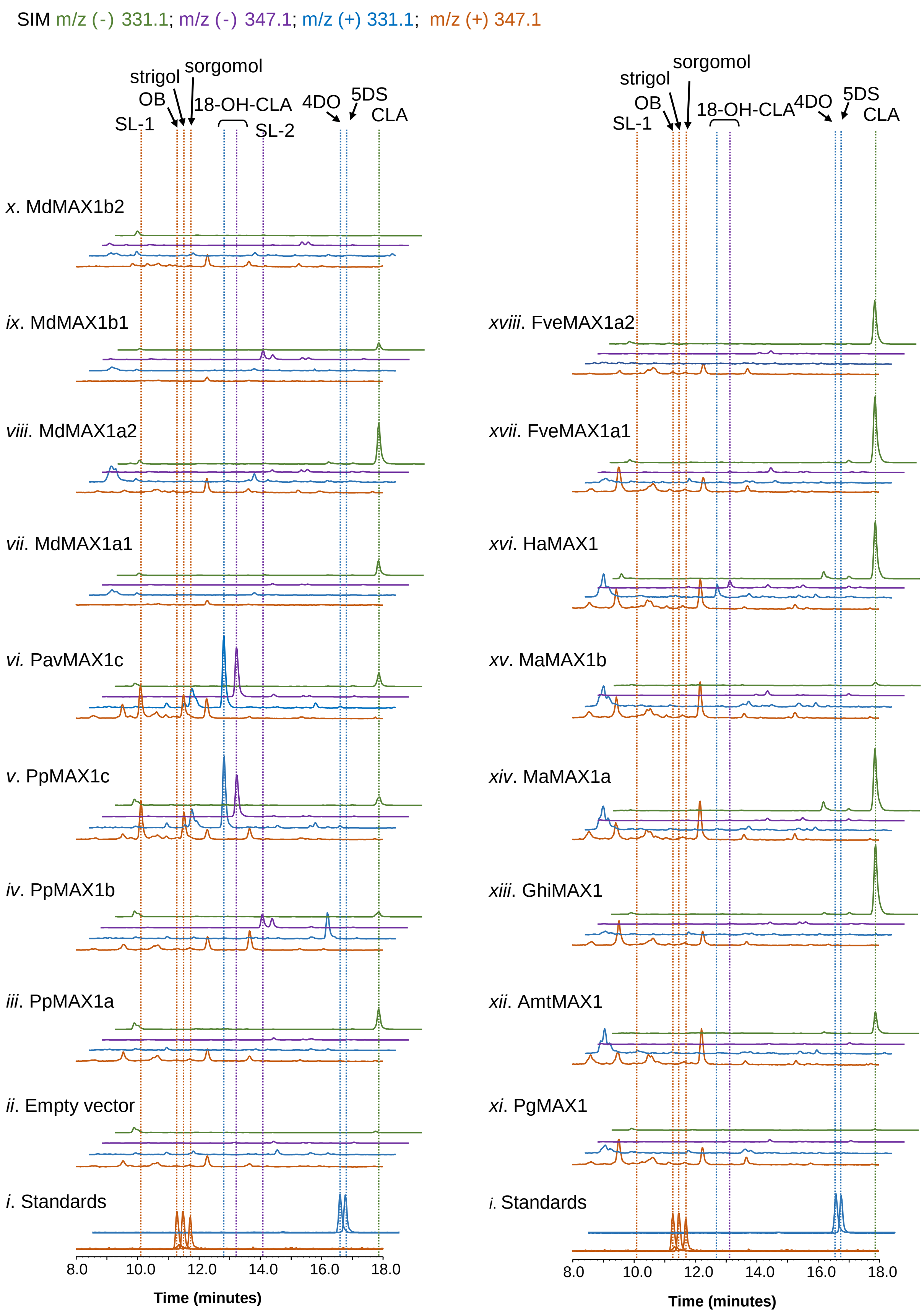

SIM m/z (-) 331.1; m/z (-) 347.1; m/z (+) 331.1; m/z (+) 347.1
strigol
sorgomol
OB
18-OH-CLA
4DO
5DS
SL-1
SL-2
CLA
x. MdMAX1b2
ix. MdMAX1b1
viii. MdMAX1a2
vii. MdMAX1a1
vi. PavMAX1c
v. PpMAX1c
iv. PpMAX1b
iii. PpMAX1a
ii. Empty vector
i. Standards
xviii. FveMAX1a2
xvii. FveMAX1a1
xvi. HaMAX1
xv. MaMAX1b
xiv. MaMAX1a
xiii. GhiMAX1
xii. AmtMAX1
xi. PgMAX1
i. Standards
8.0
10.0
12.0
14.0
16.0
18.0
Time (minutes)

**Fig. S10**: **Functional characterization of MAX1 (CYP711A) genes from strawberry, apple, cotton, banana, Amborella, white Spruce, and sunflower using SL-producing microbial consortium.** LC-MS analysis of metabolites from the co-culture strain. SIM EIC at $m/z^- = 331.1$ (green), $m/z^-$ 347.1 (purple), $m/z^+ = 331.1$ (blue), and $m/z^+ = 347.1$ (orange) of (i) strigol, OB, 4DO, and 5DS standards; CL-producing *E. coli* co-cultured (ECL) with yeast expressing ATR1 and (ii) empty vector (YSL1N); (iii) PpMAX1a (YSL1a); (iv) PpMAX1b (YSL1b); (v) PpMAX1c (YSL1c); (vi) PavMAX1c (YSL4); (vii) MdMAX1a1 (YSL6a); (viii) MdMAX1a2 (YSL6b); (ix) MdMAX1b1 (YSL6c); (x) MdMAX1b2 (YSL6d);. (xi) PgMAX1 (YSL10); (xii) AmtMAX1 (YSL9); (xiii) GhiMAX1 (YSL11); (xiv) MaMAX1a (YSL8a); (xv) MaMAX1b (YSL8b); (xvi) HaMAX1(YSL7); (xvii) FveMAX1a1 (YSL5a); and (xviii) FveMAX1a2 (YSL5b). Strigol and OB show characteristic $m/z^+$ signal ($MW = 346.38$, $[C_{19}H_{22}O_6 + H]^+ = [C_{19}H_{23}O_6]^+ = 347.1$). 4DO and 5DS show characteristic $m/z^+$ signal ($MW = 330.38$, $[C_{19}H_{22}O_5 + H]^+ = [C_{19}H_{23}O_5]^+ = 331.1$) . CLA shows characteristic $m/z^- = 331.1$ ($MW = 332.40$, $[C_{19}H_{24}O_5\text{-}H]^- = [C_{19}H_{23}O_5]^- = 331.1$); 18-OH-CLA shows characteristic $m/z^- = 347.1$ and $m/z^+ = 331.1$ ($MW = 348.40$, $[C_{19}H_{24}O_6\text{-}H]^- = [C_{19}H_{23}O_6]^- = 347.1$, $[C_{19}H_{24}O_6\text{-}H_2O\text{+}H]^+ = [C_{19}H_{23}O_5]^+ = 331.1$). Strain information are detailed in Table S3. The data is representative of at least three biological replicates.

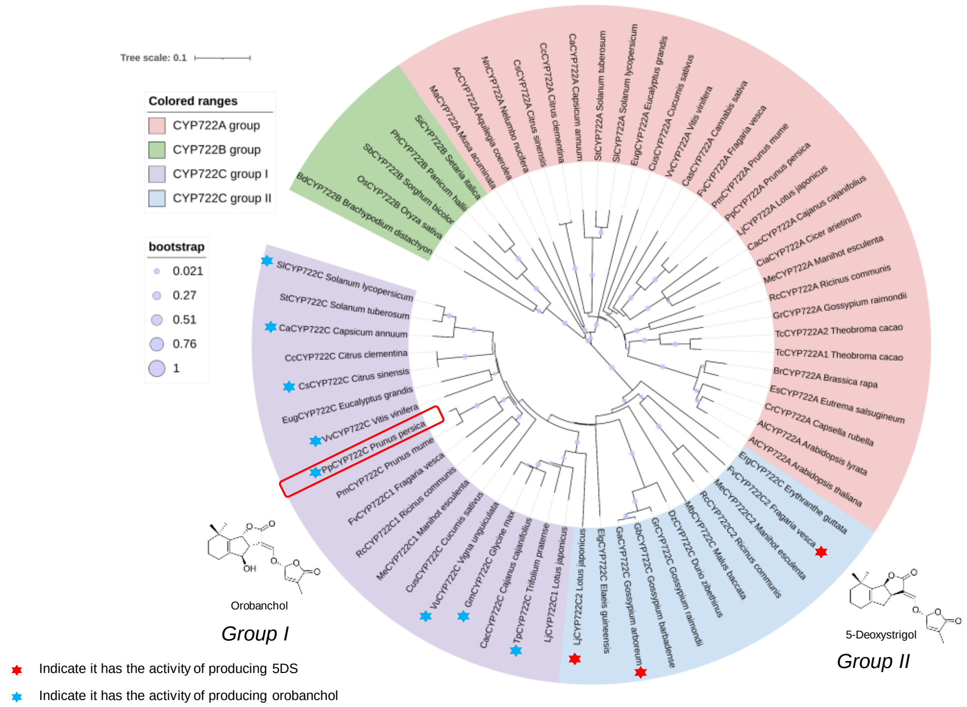


**Fig. S11**: **Extended phylogenetic analysis of CYP722 genes.** The phylogenetic analysis was conducted in MEGA X using neighbor-joining method (complete deletion, 1000 replicates, p-distance mode). A total of 61 amino acid sequences are included and the Genebank accession numbers can be found in Table S6.

**Table S1.** CYPs used in this study and summary of results. The gene IDs were extracted from Phytozome database (https://phytozome-next.jgi.doe.gov/), the accession numbers of amino acid sequences are from NCBI.

| Gene name/ ID | Accession numbers | Species | Experimental results/product |
|---|---|---|---|
| *PpMAX1a* (*Prupe.1G410300*) | XP_007222310 | *Prunus persica* | CL → CLA |
| *PpMAX1b* | XP_007224581 | *Prunus persica* | CL → CLA & U.P. |
| *PpMAX1c* (*Prupe.1G410100*) | XP_007225050 | *Prunus persica* | CL →18-OH-CLA → strigol & 5DS-derivative |
| *PavMAX1c* | XP_021823488 | *Prunus avium* | CL →18-OH-CLA → strigol & 5DS-derivative |
| *MdMAX1b1* (*MDP0000215198*) | XP_008357300 | *Malus domestica* | CL → CLA & U.P. |
| *MdMAX1b2* (*MDP0000215198*) | RXH72971 | *Malus domestica* | N.D. |
| *MdMAX1a1* (*MDP0000130133*) | XP_008393629 | *Malus domestica* | CL → CLA |
| *MdMAX1a2* (*MDP0000148030*) | XP_028955031 | *Malus domestica* | CL → CLA |
| *FveMAX1a1* (*FvH4_2g31680*) | XP_004291053 | *Fragaria vesca* | CL → CLA |
| *FveMAX1a2* (*FvH4_2g31660*) | | *Fragaria vesca* | CL → CLA |
| GhiMAX1 (*Gohir.D05G042900*) | XP_016689695 | *Gossypium hirsutum* | CL → CLA |
| AmtMAX1 (*evm_27.TU.AmTr_v1.0_scaffold00036.59*) | XP_011626843 | *Amborella trichopoda* | CL → CLA |
| PgMAX1 | AGI65359 | *Picea glauca* | N.D. |
| HaMAX1 (*HanXRQChr07g0207181*) | OTG21718 | *Helianthus annuus* | CL → CLA, trace 18-OH-CLA |
| MaMAX1a (*GSMUA_Achr10P21290_001*) | XP_009380454 | *Musa acuminata* | CL → CLA |
| MaMAX1b | XP_009408870 | *Musa acuminata* | N.D. |

U.P. Unknown product; N.D. No detected;

**Table S2.** Plasmids used in the study.

| Plasmids | Description | Reference/ Source |
|---|---|---|
| pAC-BETAipi | Contains ctrE, crtB, crtI, crtY, and idi genes of *Erwinia herbicola* (Pantoea agglomerans) Eho10 and thereby produces beta-carotene in Escherichia coli; Derived from pACYCDuet-1, Replicon P15A (pACYC184); Resistance, Chloramphenicol; | Addgene #53277 (2) |
| pYL726 (pCDFDuet-trAtCCD7-OsD27) | pCDFDuet-1 carrying D27 from *Oryza sativa* and N-terminus 31 amino-acid truncated CCD7 from *Arabidopsis* | Addgene #178285(3) |
| pYL735 (pET21a-trAtCCD8) | pET21a carrying N-terminus 56 amino-acid truncated CCD8 from *Arabidopsis* | Addgene #178286(3) |
| pAG414GPD-ccdB | *Centromeric* TRP, attR1-$P_{GPD}$-*ccdB*-$T_{CYC1}$-attR2 | (4) |
| pAG415GPD-ccdB | *Centromeric* LEU, attR1-$P_{GPD}$-*ccdB*-$T_{CYC1}$-attR2 | (4) |
| pAG416GPD-ccdB | *Centromeric* URA, attR1-$P_{GPD}$-*ccdB*-$T_{CYC1}$-attR2 | (4) |
| pYL573 | *Centromeric* HIS, $P_{TEF1}$-*ATR1*-$T_{CYC1}$ | Addgene #178288(5) |
| pYL759 | *Centromeric* LEU, $P_{PGK1}$-*AtMAX1*-$T_{pho5}$ | Addgene #178289(3) |
| pYL859 | *attL1-PpMAX1c-attL2* | This study |
| pYL864 | *Centromeric* URA, $P_{GPD}$-*PpMAX1c*-$T_{CYC1}$ | This study |
| pYL865 | *Centromeric* TRP, $P_{GPD}$-*PpMAX1b*-$T_{CYC1}$ | This study |
| pYL1081 | *Centromeric* URA, $P_{GPD}$-*PpMAX1b*-$T_{CYC1}$ | This study |
| pYL1267 | *Centromeric* URA, $P_{GPD}$-*PpMAX1cM4*-$T_{CYC1}$ | This study |
| pYL1268 | *Centromeric* URA, $P_{GPD}$-*PpMAX1cM11*-$T_{CYC1}$ | This study |
| pYL1269 | *Centromeric* URA, $P_{GPD}$-*PpMAX1cM12*-$T_{CYC1}$ | This study |
| pYL1270 | *Centromeric* URA, $P_{GPD}$-*PpMAX1cM13*-$T_{CYC1}$ | This study |
| pYL1271 | *Centromeric* URA, $P_{GPD}$-*PpMAX1cM14*-$T_{CYC1}$ | This study |
| pYL1272 | *Centromeric* URA, $P_{GPD}$-*PpMAX1cM16*-$T_{CYC1}$ | This study |
| pYL1273 | *Centromeric* URA, $P_{GPD}$-*PpMAX1cM17*-$T_{CYC1}$ | This study |
| pYL1274 | *Centromeric* URA, $P_{GPD}$-*PpMAX1cM20*-$T_{CYC1}$ | This study |
| pYL1275 | *Centromeric* URA, $P_{GPD}$-*PpMAX1cM22*-$T_{CYC1}$ | This study |
| pYL1276 | *Centromeric* URA, $P_{GPD}$-*PpMAX1cM28*-$T_{CYC1}$ | This study |
| pYL1308 | *Centromeric* URA, $P_{GPD}$-*PpMAX1cM15*-$T_{CYC1}$ | This study |
| pYL1309 | *Centromeric* URA, $P_{GPD}$-*PpMAX1cM19*-$T_{CYC1}$ | This study |
| pYL1310 | *Centromeric* URA, $P_{GPD}$-*PpMAX1a*-$T_{CYC1}$ | This study |
| pYL1318 | *Centromeric* TRP, $P_{GPD}$-*PpCYP722C*-$T_{CYC1}$ | This study |
| pYL1319 | *Centromeric* URA, $P_{GPD}$-*PavMAX1c*-$T_{CYC1}$ | This study |
| pYL1333 | *Centromeric* URA, $P_{GPD}$-*PpMAX1cM2*-$T_{CYC1}$ | This study |
| pYL1334 | *Centromeric* URA, $P_{GPD}$-*PpMAX1cM5*-$T_{CYC1}$ | This study |
| pYL1335 | *Centromeric* URA, $P_{GPD}$-*PpMAX1cM32*-$T_{CYC1}$ | This study |
| pYL1336 | *Centromeric* URA, $P_{GPD}$-*PpMAX1cM33*-$T_{CYC1}$ | This study |
| pYL1337 | *Centromeric* URA, $P_{GPD}$-*PpMAX1cM34*-$T_{CYC1}$ | This study |
| pYL1360 | *Centromeric* URA, $P_{GPD}$-*MdMAX1b2*-$T_{CYC1}$ | This study |
| pYL1365 | *Centromeric* URA, $P_{GPD}$-*PpMAX1cM1*-$T_{CYC1}$ | This study |
| pYL1366 | *Centromeric* URA, $P_{GPD}$-*PpMAX1cM3*-$T_{CYC1}$ | This study |
| pYL1367 | *Centromeric* URA, $P_{GPD}$-*PpMAX1cM6*-$T_{CYC1}$ | This study |
| pYL1368 | *Centromeric* URA, $P_{GPD}$-*PpMAX1cM7*-$T_{CYC1}$ | This study |
| pYL1369 | *Centromeric* URA, $P_{GPD}$-*PpMAX1cM8*-$T_{CYC1}$ | This study |
| pYL1370 | *Centromeric* URA, $P_{GPD}$-*PpMAX1cM9*-$T_{CYC1}$ | This study |
| pYL1371 | *Centromeric* URA, $P_{GPD}$-*PpMAX1cM10*-$T_{CYC1}$ | This study |
| pYL1373 | *Centromeric* URA, $P_{GPD}$-*PpMAX1cM18*-$T_{CYC1}$ | This study |

| | | |
|---|---|---|
| pYL1374 | *Centromeric* URA, $P_{GPD}$-*PpMAX1cM21*-$T_{CYC1}$ | This study |
| pYL1375 | *Centromeric* URA, $P_{GPD}$-*PpMAX1cM23*-$T_{CYC1}$ | This study |
| pYL1376 | *Centromeric* URA, $P_{GPD}$-*PpMAX1cM24*-$T_{CYC1}$ | This study |
| pYL1377 | *Centromeric* URA, $P_{GPD}$-*PpMAX1cM25*-$T_{CYC1}$ | This study |
| pYL1378 | *Centromeric* URA, $P_{GPD}$-*PpMAX1cM26*-$T_{CYC1}$ | This study |
| pYL1379 | *Centromeric* URA, $P_{GPD}$-*PpMAX1cM27*-$T_{CYC1}$ | This study |
| pYL1380 | *Centromeric* URA, $P_{GPD}$-*PpMAX1cM29*-$T_{CYC1}$ | This study |
| pYL1381 | *Centromeric* URA, $P_{GPD}$-*PpMAX1cM30*-$T_{CYC1}$ | This study |
| pYL1382 | *Centromeric* URA, $P_{GPD}$-*PpMAX1cM31*-$T_{CYC1}$ | This study |
| pYL1385 | *Centromeric* URA, $P_{GPD}$-*MdMAX1b1*-$T_{CYC1}$ | This study |
| pYL1387 | *Centromeric* URA, $P_{GPD}$-*MdMAX1a2*-$T_{CYC1}$ | This study |
| pYL1427 | *Centromeric* URA, $P_{GPD}$-*MdMAX1a1*-$T_{CYC1}$ | This study |
| pYL1578 | *Centromeric* LEU, $P_{GPD}$-*PpCYP722C*-$T_{CYC1}$ | This study |
| pYL1580 | *Centromeric* URA, $P_{GPD}$-*FveMAX1a1*-$T_{CYC1}$ | This study |
| pYL1581 | *Centromeric* URA, $P_{GPD}$-*FveMAX1a2*-$T_{CYC1}$ | This study |
| pYL1585 | *Centromeric* LEU, $P_{GPD}$-*PpMAX1a*-$T_{CYC1}$ | This study |
| pYL1591 | *Centromeric* LEU, $P_{GPD}$-*HaMAX1*-$T_{CYC1}$ | This study |
| pYL1592 | *Centromeric* LEU, $P_{GPD}$-*MaMAX1a*-$T_{CYC1}$ | This study |
| pYL1593 | *Centromeric* LEU, $P_{GPD}$-*MaMAX1b*-$T_{CYC1}$ | This study |
| pYL1612 | *Centromeric* LEU, $P_{GPD}$-*AmtMAX1*-$T_{CYC1}$ | This study |
| pYL1622 | *Centromeric* LEU, $P_{GPD}$-*PgMAX1*-$T_{CYC1}$ | This study |
| pYL1623 | *Centromeric* LEU, $P_{GPD}$-*GhiMAX1*-$T_{CYC1}$ | This study |

*P, promoter; T, terminator*

**Table S3.** Strains used in the study.

| Yeast Strain | Description | Genotype | Function | Reference |
|---|---|---|---|---|
| CEN.PK2-1D | | *MATα; his3D1; leu2-3_112; ura3-52; trp1-289; MAL2-8c; SUC2* | Wild type yeast strain | (6) |
| YSL1N | CEN.PK2-1D carrying pYL573 and pAG416GPD-ccdB | Centromeric HIS, $P_{TEF1}$-*ATR1*-$T_{CYC1}$, Centromeric URA, $P_{GPD}$-*ccdB*-$T_{CYC1}$ | Negative control for YSL1 and YSL3 | This study |
| YSL1a | CEN.PK2-1D carrying pYL573 and pYL1310 | Centromeric HIS, $P_{TEF1}$-*ATR1*-$T_{CYC1}$, Centromeric URA, $P_{GPD}$-*PpMAX1a*-$T_{CYC1}$ | CLA production | This study |
| YSL1b | CEN.PK2-1D carrying pYL573 and pYL1081 | Centromeric HIS, $P_{TEF1}$-*ATR1*-$T_{CYC1}$, Centromeric URA, $P_{GPD}$-*PpMAX1b*-$T_{CYC1}$ | CLA and unknown production | This study |
| YSL1c | CEN.PK2-1D carrying pYL573 and pYL864 | Centromeric HIS, $P_{TEF1}$-*ATR1*-$T_{CYC1}$, Centromeric URA, $P_{GPD}$-*PpMAX1c*-$T_{CYC1}$ | Strigol & 5DS-derivative production | This study |
| YSL2a | CEN.PK2-1D carrying pYL573, pYL1585 and pYL865 | Centromeric HIS, $P_{TEF1}$-*ATR1*-$T_{CYC1}$, Centromeric LEU, $P_{GPD}$-*PpMAX1a*-$T_{CYC1}$ Centromeric TRP, $P_{GPD}$-*PpMAX1b*-$T_{CYC1}$ | CLA and unknown production | This study |
| YSL2b | CEN.PK2-1D carrying pYL573, pYL864 and pYL1585 | Centromeric HIS, $P_{TEF1}$-*ATR1*-$T_{CYC1}$, Centromeric URA, $P_{GPD}$-*PpMAX1c*-$T_{CYC1}$ Centromeric LEU, $P_{GPD}$-*PpMAX1a*-$T_{CYC1}$ | Strigol & 5DS-derivative production | This study |
| YSL2c | CEN.PK2-1D carrying pYL573, pYL864 and pYL865 | Centromeric HIS, $P_{TEF1}$-*ATR1*-$T_{CYC1}$, Centromeric URA, $P_{GPD}$-*PpMAX1c*-$T_{CYC1}$ Centromeric TRP, $P_{GPD}$-*PpMAX1b*-$T_{CYC1}$ | Strigol & 5DS-derivative and unknown production | This study |
| YSL3M1 | CEN.PK2-1D carrying pYL573 and pYL1365 | Centromeric HIS, $P_{TEF1}$-*ATR1*-$T_{CYC1}$, Centromeric URA, $P_{GPD}$-*PpMAX1cM1*-$T_{CYC1}$ | Strigol & 5DS-derivative production | This study |
| YSL3M2 | CEN.PK2-1D carrying pYL573 and pYL1333 | Centromeric HIS, $P_{TEF1}$-*ATR1*-$T_{CYC1}$, Centromeric URA, $P_{GPD}$-*PpMAX1cM2*-$T_{CYC1}$ | No strigol & 5DS-derivative production | This study |
| YSL3M3 | CEN.PK2-1D carrying pYL573 and pYL1366 | Centromeric HIS, $P_{TEF1}$-*ATR1*-$T_{CYC1}$, Centromeric URA, $P_{GPD}$-*PpMAX1cM3*-$T_{CYC1}$ | Strigol & 5DS-derivative production | This study |
| YSL3M4 | CEN.PK2-1D carrying pYL573 and pYL1267 | Centromeric HIS, $P_{TEF1}$-*ATR1*-$T_{CYC1}$, Centromeric URA, $P_{GPD}$-*PpMAX1cM4*-$T_{CYC1}$ | Almost abolished the synthesis of strigol but showed no change in the synthesis of the 5DS-derivative | This study |
| YSL3M5 | CEN.PK2-1D carrying pYL573 and pYL1334 | Centromeric HIS, $P_{TEF1}$-*ATR1*-$T_{CYC1}$, Centromeric URA, $P_{GPD}$-*PpMAX1cM5*-$T_{CYC1}$ | Strigol & 5DS-derivative production | This study |
| YS3M6 | CEN.PK2-1D carrying pYL573 and pYL1367 | Centromeric HIS, $P_{TEF1}$-*ATR1*-$T_{CYC1}$, Centromeric URA, $P_{GPD}$-*PpMAX1cM6*-$T_{CYC1}$ | Strigol & 5DS-derivative production | This study |

| | | | | |
|---|---|---|---|---|
| YSL3M7 | CEN.PK2-1D carrying pYL573 and pYL1368 | Centromeric HIS, $P_{TEF1}$-*ATR1*-$T_{CYC1}$, Centromeric URA, $P_{GPD}$-*PpMAX1cM7*-$T_{CYC1}$ | Strigol & 5DS-derivative production | This study |
| YSL3M8 | CEN.PK2-1D carrying pYL573 and pYL1369 | Centromeric HIS, $P_{TEF1}$-*ATR1*-$T_{CYC1}$, Centromeric URA, $P_{GPD}$-*PpMAX1cM8*-$T_{CYC1}$ | Strigol & 5DS-derivative production | This study |
| YSL3M9 | CEN.PK2-1D carrying pYL573 and pYL1370 | Centromeric HIS, $P_{TEF1}$-*ATR1*-$T_{CYC1}$, Centromeric URA, $P_{GPD}$-*PpMAX1cM9*-$T_{CYC1}$ | Strigol & 5DS-derivative production | This study |
| YSL3M10 | CEN.PK2-1D carrying pYL573 and pYL1371 | Centromeric HIS, $P_{TEF1}$-*ATR1*-$T_{CYC1}$, Centromeric URA, $P_{GPD}$-*PpMAX1cM10*-$T_{CYC1}$ | Strigol & 5DS-derivative production | This study |
| YSL3M11 | CEN.PK2-1D carrying pYL573 and pYL1268 | Centromeric HIS, $P_{TEF1}$-*ATR1*-$T_{CYC1}$, Centromeric URA, $P_{GPD}$-*PpMAX1cM11*-$T_{CYC1}$ | Strigol & 5DS-derivative production | This study |
| YSL3M12 | CEN.PK2-1D carrying pYL573 and pYL1269 | Centromeric HIS, $P_{TEF1}$-*ATR1*-$T_{CYC1}$, Centromeric URA, $P_{GPD}$-*PpMAX1cM12*-$T_{CYC1}$ | Strigol & 5DS-derivative production | This study |
| YSL3M13 | CEN.PK2-1D carrying pYL573 and pYL1270 | Centromeric HIS, $P_{TEF1}$-*ATR1*-$T_{CYC1}$, Centromeric URA, $P_{GPD}$-*PpMAX1cM13*-$T_{CYC1}$ | Strigol & 5DS-derivative production | This study |
| YSL3M14 | CEN.PK2-1D carrying pYL573 and pYL1271 | Centromeric HIS, $P_{TEF1}$-*ATR1*-$T_{CYC1}$, Centromeric URA, $P_{GPD}$-*PpMAX1cM14*-$T_{CYC1}$ | Strigol & 5DS-derivative production | This study |
| YSL3M15 | CEN.PK2-1D carrying pYL573 and pYL1308 | Centromeric HIS, $P_{TEF1}$-*ATR1*-$T_{CYC1}$, Centromeric URA, $P_{GPD}$-*PpMAX1cM15*-$T_{CYC1}$ | Strigol & 5DS-derivative production | This study |
| YSL3M16 | CEN.PK2-1D carrying pYL573 and pYL1272 | Centromeric HIS, $P_{TEF1}$-*ATR1*-$T_{CYC1}$, Centromeric URA, $P_{GPD}$-*PpMAX1cM16*-$T_{CYC1}$ | Strigol & 5DS-derivative production | This study |
| YSL3M17 | CEN.PK2-1D carrying pYL573 and pYL1273 | Centromeric HIS, $P_{TEF1}$-*ATR1*-$T_{CYC1}$, Centromeric URA, $P_{GPD}$-*PpMAX1cM17*-$T_{CYC1}$ | Strigol & 5DS-derivative production | This study |
| YSL3M18 | CEN.PK2-1D carrying pYL573 and pYL1373 | Centromeric HIS, $P_{TEF1}$-*ATR1*-$T_{CYC1}$, Centromeric URA, $P_{GPD}$-*PpMAX1cM18*-$T_{CYC1}$ | Strigol & 5DS-derivative production | This study |
| YSL3M19 | CEN.PK2-1D carrying pYL573 and pYL1337 | Centromeric HIS, $P_{TEF1}$-*ATR1*-$T_{CYC1}$, Centromeric URA, $P_{GPD}$-*PpMAX1cM19*-$T_{CYC1}$ | Strigol & 5DS-derivative production | This study |
| YSL3M20 | CEN.PK2-1D carrying pYL573 and pYL1274 | Centromeric HIS, $P_{TEF1}$-*ATR1*-$T_{CYC1}$, Centromeric URA, $P_{GPD}$-*PpMAX1cM20*-$T_{CYC1}$ | Strigol & 5DS-derivative production | This study |
| YSL3M21 | CEN.PK2-1D carrying pYL573 and pYL1374 | Centromeric HIS, $P_{TEF1}$-*ATR1*-$T_{CYC1}$, Centromeric URA, $P_{GPD}$-*PpMAX1cM21*-$T_{CYC1}$ | Strigol & 5DS-derivative production | This study |
| YSL3M22 | CEN.PK2-1D carrying pYL573 and pYL1275 | Centromeric HIS, $P_{TEF1}$-*ATR1*-$T_{CYC1}$, Centromeric URA, $P_{GPD}$-*PpMAX1cM22*-$T_{CYC1}$ | Strigol & 5DS-derivative production | This study |
| YSL3M23 | CEN.PK2-1D carrying pYL573 and pYL1375 | Centromeric HIS, $P_{TEF1}$-*ATR1*-$T_{CYC1}$, Centromeric URA, $P_{GPD}$-*PpMAX1cM23*-$T_{CYC1}$ | Strigol & 5DS-derivative production | This study |
| YSL3M24 | CEN.PK2-1D carrying pYL573 and pYL1376 | Centromeric HIS, $P_{TEF1}$-*ATR1*-$T_{CYC1}$, Centromeric URA, $P_{GPD}$-*PpMAX1cM24*-$T_{CYC1}$ | Strigol & 5DS-derivative production | This study |
| YSL3M25 | CEN.PK2-1D carrying pYL573 and pYL1377 | Centromeric HIS, $P_{TEF1}$-*ATR1*-$T_{CYC1}$, Centromeric URA, $P_{GPD}$-*PpMAX1cM25*-$T_{CYC1}$ | Strigol & 5DS-derivative production | This study |

| | | | | |
|---|---|---|---|---|
| YSL3M26 | CEN.PK2-1D carrying pYL573 and pYL1378 | Centromeric HIS, $P_{TEF1}$-*ATR1*-$T_{CYC1}$, Centromeric URA, $P_{GPD}$-*PpMAX1cM26*-$T_{CYC1}$ | interruption of production of downstream products of CLA | This study |
| YSL3M27 | CEN.PK2-1D carrying pYL573 and pYL1379 | Centromeric HIS, $P_{TEF1}$-*ATR1*-$T_{CYC1}$, Centromeric URA, $P_{GPD}$-*PpMAX1cM27*-$T_{CYC1}$ | Strigol & 5DS-derivative production | This study |
| YSL3M28 | CEN.PK2-1D carrying pYL573 and pYL1276 | Centromeric HIS, $P_{TEF1}$-*ATR1*-$T_{CYC1}$, Centromeric URA, $P_{GPD}$-*PpMAX1cM28*-$T_{CYC1}$ | Strigol & 5DS-derivative production | This study |
| YSL3M29 | CEN.PK2-1D carrying pYL573 and pYL1380 | Centromeric HIS, $P_{TEF1}$-*ATR1*-$T_{CYC1}$, Centromeric URA, $P_{GPD}$-*PpMAX1cM29*-$T_{CYC1}$ | Strigol & 5DS-derivative production | This study |
| YSL3M30 | CEN.PK2-1D carrying pYL573 and pYL1381 | Centromeric HIS, $P_{TEF1}$-*ATR1*-$T_{CYC1}$, Centromeric URA, $P_{GPD}$-*PpMAX1cM30*-$T_{CYC1}$ | Strigol & 5DS-derivative production | This study |
| YSL3M31 | CEN.PK2-1D carrying pYL573 and pYL1382 | Centromeric HIS, $P_{TEF1}$-*ATR1*-$T_{CYC1}$, Centromeric URA, $P_{GPD}$-*PpMAX1cM31*-$T_{CYC1}$ | No strigol & 5DS-derivative production | This study |
| YSL3M32 | CEN.PK2-1D carrying pYL573 and pYL1335 | Centromeric HIS, $P_{TEF1}$-*ATR1*-$T_{CYC1}$, Centromeric URA, $P_{GPD}$-*PpMAX1cM32*-$T_{CYC1}$ | No strigol & 5DS-derivative production | This study |
| YSL3M33 | CEN.PK2-1D carrying pYL573 and pYL1336 | Centromeric HIS, $P_{TEF1}$-*ATR1*-$T_{CYC1}$, Centromeric URA, $P_{GPD}$-*PpMAX1cM33*-$T_{CYC1}$ | No strigol & 5DS-derivative production | This study |
| YSL3M34 | CEN.PK2-1D carrying pYL573 and pYL1337 | Centromeric HIS, $P_{TEF1}$-*ATR1*-$T_{CYC1}$, Centromeric URA, $P_{GPD}$-*PpMAX1cM34*-$T_{CYC1}$ | No strigol & 5DS-derivative production | This study |
| YSL4 | CEN.PK2-1D carrying pYL573 and pYL1319 | Centromeric HIS, $P_{TEF1}$-*ATR1*-$T_{CYC1}$, Centromeric URA, $P_{GPD}$-*PavMAX1c*-$T_{CYC1}$ | Strigol & 5DS-derivative production | This study |
| YSL5a | CEN.PK2-1D carrying pYL573 and pYL1580 | Centromeric HIS, $P_{TEF1}$-*ATR1*-$T_{CYC1}$, Centromeric URA, $P_{GPD}$-*FveMAX1a1*-$T_{CYC1}$ | CLA production | This study |
| YSL5b | CEN.PK2-1D carrying pYL573 and pYL1581 | Centromeric HIS, $P_{TEF1}$-*ATR1*-$T_{CYC1}$, Centromeric URA, $P_{GPD}$-*FveMAX1a2*-$T_{CYC1}$ | CLA production | This study |
| YSL6a | CEN.PK2-1D carrying pYL573 and pYL1427 | Centromeric HIS, $P_{TEF1}$-*ATR1*-$T_{CYC1}$, Centromeric URA, $P_{GPD}$-*MdMAX1a1*-$T_{CYC1}$ | CLA production | This study |
| YSL6b | CEN.PK2-1D carrying pYL573 and pYL1387 | Centromeric HIS, $P_{TEF1}$-*ATR1*-$T_{CYC1}$, Centromeric URA, $P_{GPD}$-*MdMAX1a2*-$T_{CYC1}$ | CLA production | This study |
| YSL6c | CEN.PK2-1D carrying pYL573 and pYL1385 | Centromeric HIS, $P_{TEF1}$-*ATR1*-$T_{CYC1}$, Centromeric URA, $P_{GPD}$-*MdMAX1b1*-$T_{CYC1}$ | CLA and unknown production | This study |
| YSL6d | CEN.PK2-1D carrying pYL573 and pYL1360 | Centromeric HIS, $P_{TEF1}$-*ATR1*-$T_{CYC1}$, Centromeric URA, $P_{GPD}$-*MdMAX1b2*-$T_{CYC1}$ | Failed functional characterization of MdMAX1b2 | This study |
| YSL7N | CEN.PK2-1D carrying pYL573 and pAG415GPD-ccdB | Centromeric HIS, $P_{TEF1}$-*ATR1*-$T_{CYC1}$, Centromeric LEU, $P_{GPD}$-*ccdB*-$T_{CYC1}$ | Negative control for YSL7-12 | This study |
| YSL7 | CEN.PK2-1D carrying pYL573 and pYL1591 | Centromeric HIS, $P_{TEF1}$-*ATR1*-$T_{CYC1}$, Centromeric LEU, $P_{GPD}$-*HaMAX1*-$T_{CYC1}$ | CLA production | This study |

| | | | | |
|---|---|---|---|---|
| YSL8a | CEN.PK2-1D carrying pYL573 and pYL1592 | Centromeric HIS, $P_{TEF1}$-*ATR1*-$T_{CYC1}$, Centromeric LEU, $P_{GPD}$-*MaMAX1a*-$T_{CYC1}$ | CLA production | This study |
| YSL8b | CEN.PK2-1D carrying pYL573 and pYL1593 | Centromeric HIS, $P_{TEF1}$-*ATR1*-$T_{CYC1}$, Centromeric LEU, $P_{GPD}$-*MaMAX1b*-$T_{CYC1}$ | Failed functional characterization of MaMAX1b | This study |
| YSL9 | CEN.PK2-1D carrying pYL573 and pYL1612 | Centromeric HIS, $P_{TEF1}$-*ATR1*-$T_{CYC1}$, Centromeric LEU, $P_{GPD}$-*AmtMAX1*-$T_{CYC1}$ | CLA production | This study |
| YSL10 | CEN.PK2-1D carrying pYL573 and pYL1622 | Centromeric HIS, $P_{TEF1}$-*ATR1*-$T_{CYC1}$, Centromeric LEU, $P_{GPD}$-*PgMAX1*-$T_{CYC1}$ | Failed functional characterization of PgMAX1 | This study |
| YSL11 | CEN.PK2-1D carrying pYL573 and pYL1623 | Centromeric HIS, $P_{TEF1}$-*ATR1*-$T_{CYC1}$, Centromeric LEU, $P_{GPD}$-*GhiMAX1*-$T_{CYC1}$ | CLA production | This study |
| YSL12 | CEN.PK2-1D carrying pYL573 and pYL891 | Centromeric HIS, $P_{TEF1}$-*ATR1*-$T_{CYC1}$, Centromeric LEU, $P_{GPD}$-*SbMAX1a*-$T_{CYC1}$ | OB and 18-OH-CLA production | (7) |
| YSL13 | CEN.PK2-1D carrying pYL573, pYL759, and pYL1318 | Centromeric HIS, $P_{TEF1}$-*ATR1*-$T_{CYC1}$, Centromeric LEU, $P_{PGK1}$-*AtMAX1*-$T_{PHO5}$, Centromeric TRP, $P_{GPD}$-*PpCYP722C*-$T_{CYC1}$ | OB production | This study |
| YSL13N | CEN.PK2-1D carrying pYL573, pYL759 and pAG416GPD-ccdB | Centromeric HIS, $P_{TEF1}$-*ATR1*-$T_{CYC1}$, Centromeric LEU, $P_{PGK1}$-*AtMAX1*-$T_{PHO5}$, Centromeric TRP, $P_{GPD}$-*ccdB*-$T_{CYC1}$ | Negative control for YSL13 CLA production | This study |
| YSL13N' | CEN.PK2-1D carrying pYL573 and pYL759 | Centromeric HIS, $P_{TEF1}$-*ATR1*-$T_{CYC1}$, Centromeric LEU, $P_{PGK1}$-*AtMAX1*-$T_{PHO5}$ | CLA production | This study |
| YSL14a | CEN.PK2-1D carrying pYL573, pYL1578 and pYL1310 | Centromeric HIS, $P_{TEF1}$-*ATR1*-$T_{CYC1}$, Centromeric LEU, $P_{GPD}$-*PpCYP722C*-$T_{CYC1}$ Centromeric URA, $P_{GPD}$-*PpMAX1a*-$T_{CYC1}$ | CLA and OB production | This study |
| YSL14b | CEN.PK2-1D carrying pYL573, pYL1578 and pYL1081 | Centromeric HIS, $P_{TEF1}$-*ATR1*-$T_{CYC1}$, Centromeric LEU, $P_{GPD}$-*PpCYP722C*-$T_{CYC1}$ Centromeric URA, $P_{GPD}$-*PpMAX1b*-$T_{CYC1}$ | OB, CLA and unknown production | This study |
| YSL14c | CEN.PK2-1D carrying pYL573, pYL1578 and pYL864 | Centromeric HIS, $P_{TEF1}$-*ATR1*-$T_{CYC1}$, Centromeric LEU, $P_{GPD}$-*PpCYP722C*-$T_{CYC1}$ Centromeric URA, $P_{GPD}$-*PpMAX1c*-$T_{CYC1}$ | Strigol & 5DS-derivative and OB production | This study |
| ***E. coli* Strain** | **Base Strain** | **Plasmid** | **Function** | |
| ECL | BL21(DE3) | pAC-BETAipi; pYL726 (pCDFDuet-trAtCCD7-OsD27); pYL735 (pET21a-trAtCCD8) | CL production | (3) |

**Table S4.** PpMAX1c mutants used in this study and summary of results.

| Strain name | Mutation | CLA | 18-OH-CLA | 5DS-derivative | strigol |
|---|---|---|---|---|---|
| PpMAX1c WT | Wild type | √ | √ | √ | √ |
| PpMAX1cM1 | QI66SV | √ | √ | √ | √ |
| PpMAX1cM2 | R108A | × | × | × | × |
| PpMAX1cM3 | Q115A | √ | √ | √ | √ |
| PpMAX1cM4 | C117S | √ | √ | √ trace | √ |
| PpMAX1cM5 | R136A | √ | √ | √ | √ |
| PpMAX1cM6 | L139I | √ | √ | √ | √ |
| PpMAX1cM7 | V159I | √ | √ | √ | √ |
| PpMAX1cM8 | R164K | √ | √ | √ | √ |
| PpMAX1cM9 | K179N | √ | √ | √ | √ |
| PpMAX1cM10 | A183K | √ | √ | √ | √ |
| PpMAX1cM11 | T188I | √ | √ | √ | √ |
| PpMAX1cM12 | A190G | √ | √ | √ | √ |
| PpMAX1cM13 | K226D | √ | √ | √ | √ |
| PpMAX1cM14 | T228I | √ | √ | √ | √ |
| PpMAX1cM15 | Y231H | √ | √ | √ | √ |
| PpMAX1cM16 | VA235TT | √ | √ | √ | √ |
| PpMAX1cM17 | F254V | √ | √ | √ | √ |
| PpMAX1cM18 | D281A | √ | √ | √ | √ |
| PpMAX1cM19 | H335Q | √ | √ | √ | √ |
| PpMAX1cM20 | T350V | √ | √ | √ | √ |
| PpMAX1cM21 | Q356G | √ | √ | √ | √ |
| PpMAX1cM22 | Y357H | √ | √ | √ | √ |
| PpMAX1cM23 | L368I | √ | √ | √ | √ |
| PpMAX1cM24 | V390L | √ | √ | √ | √ |
| PpMAX1cM25 | L400F | √ | √ | √ | √ |
| PpMAX1cM26 | P402M | √ | × | × | × |
| PpMAX1cM27 | S413K | √ | √ | √ | √ |
| PpMAX1cM28 | FG430LA | √ | √ | √ | √ |
| PpMAX1cM29 | I435L | √ | √ | √ | √ |
| PpMAX1cM30 | N446E | √ | √ | √ | √ |
| PpMAX1cM31 | E462Q | × | × | × | × |
| PpMAX1cM32 | P470**A** | √ | √ | × | × |
| PpMAX1cM33 | L479A | √ | √ | × | × |
| PpMAX1cM34 | G480A | √ | √ | × | × |

**Table S5.** Accession numbers of MAX1 analogs used for the phylogenetic tree analysis in Fig. 2 and multiple alignment in Fig. 4 and Fig. S8. The amino acid sequences are downloadable from NCBI or Phytozom (https://phytozome.jgi.doe.gov/pz/portal.html).

| Gene | Species | Size (a.a.) | Accession numbers |
|---|---|---|---|
| *AtMAX1* | *Arabidopsis thaliana* | 522 | NP_565617 |
| *SmMAX1a* | *Selaginella moellendorffii* | 512 | AGI65366 |
| *OsMAX1b* (*Os01g0700900*) | *Oryza sativa* (Rice) | 539 | XP_015633367 |
| *OsMAX1c* (*Os01g0701400*) | *Oryza sativa* (Rice) | 541 | XP_015644699 |
| *OsMAX1d* (*Os01g0701500*) | *Oryza sativa* (Rice) | 516 | XP_015642272 |
| *OsMAX1e* (*Os02g0221900*) | *Oryza sativa* (Rice) | 548 | XP_015626073 |
| *OsMAX1f* (*Os06g0565100*) | *Oryza sativa* (Rice) | 540 | XP_015644019 |
| *SlMAX1* | *Solanum lycopersicum* (Tomato) | 519 | XP_004245085 |
| *ZmMAX1a* | *Zea mays* (Corn) | 537 | PWZ07057 |
| *ZmMAX1b* | *Zea mays* (Corn) | 543 | ONM29770 |
| *ZmMAX1c* | *Zea mays* (Corn) | 560 | XP_020407074 |
| *AmtMAX1* | *Amborella trichopoda* | 555 | XP_011626843 |
| *PpMAX1a* (*Prupe.1G410300*) | *Prunus persica* (Peach) | 538 | XP_007222310 |
| *PpMAX1b* | *Prunus persica* (Peach) | 533 | XP_007224581 |
| *PpMAX1c* (*Prupe.1G410100*) | *Prunus persica* (Peach) | 536 | XP_007225050 |
| *MdMAX1a1* (*MDP0000130133*) | *Malus domestica* (Apple) | 540 | XP_008393629 |
| *MdMAX1a2* (*MDP0000148030*) | *Malus domestica* (Apple) | 532 | XP_028955031 |
| *MdMAX1b1* (*MDP0000215198*) | *Malus domestica* (Peach) | 526 | XP_008357300 |
| *MdMAX1b2* (*MDP0000231714*) | *Malus domestica* (Apple) | 542 | RXH72971 |
| *FveMAX1a1* (*FvH4_2g31680*) | *Fragaria vesca* (Woodland strawberry) | 531 | XP_004291053 |
| *FveMAX1a2* (*FvH4_2g31660*) | *Fragaria vesca* (Woodland strawberry) | 531 | FvH4_2g31660.t1 |
| *AcMAX1* | *Actinidia chinensis* (Chinese kiwifruit) | 537 | PSR91738 |
| *ClMAX1* | *Citrullus lanatus* (Watermelon) | 526 | Cla97C05G099260.1 |
| *CcMAX1a* | *Citrus clementina* (Clementine) | 538 | ESR64106 |
| *CcMAX1b* | *Citrus clementina* (Clementine) | 547 | ESR38800 |
| *HaMAX1* | *Helianthus annuus* (Common sunflower) | 521 | OTG21718 |
| *MeMAX1* | *Manihot esculenta* (Cassava) | 527 | OAY26871 |
| *MaMAX1a* | *Musa acuminata* (Banana) | 529 | XP_009380454 |
| *MaMAX1b* | *Musa acuminata* (Banana) | 526 | XP_009408870 |
| *PdMAX1a* | *Prunus dulcis* (Almond) | 538 | XP_034198777 |
| *PdMAX1b* | *Prunus dulcis* (Almond) | 533 | XP_034200540 |
| *PdMAX1c* | *Prunus dulcis* (Almond) | 535 | XP_034203783 |

| | | | |
|---|---|---|---|
| *PavMAX1a* | *Prunus avium* (Sweet Cherry) | 533 | XP_021823617 |
| *PavMAX1b* | *Prunus avium* (Sweet Cherry) | 535 | XP_021823330 |
| *PavMAX1c* | *Prunus avium* (Sweet Cherry) | 541 | XP_021823488 |
| *ParMAX1a* | *Prunus armeniaca* (Apricot) | 538 | CAB4266066 |
| *ParMAX1b* | *Prunus armeniaca* (Apricot) | 533 | CAB4296645 |
| *ParMAX1c* | *Prunus armeniaca* (Apricot) | 544 | CAB4296646 |
| *GhiMAX1* | *Gossypium hirsutum* (Cotton) | 539 | XP_016689695 |
| *AhMAX1a* | *Arachis hypogaea* (Peanut) | 539 | XP_025606701 |
| *AhMAX1b* | *Arachis hypogaea* (Peanut) | 539 | XP_025660007 |
| *AhMAX1c* | *Arachis hypogaea* (Peanut) | 528 | XP_025613410 |
| *GmMAX1a* | *Glycine max* (Soybean) | 551 | AQY54419 |
| *GmMAX1b* | *Glycine max* (Soybean) | 548 | AQY54420 |
| *GmMAX1c* | *Glycine max* (Soybean) | 532 | XP_003549345 |
| *GmMAX1d* | *Glycine max* (Soybean) | 538 | XP_003544542 |
| *PgMAX1* | *Picea glauca* (White Spruce) | 544 | AGI65359 |
| *SbMAX1a* | *Sorghum bicolor* (Sorghum) | 547 | XP_002458367 |
| *SbMAX1b* | *Sorghum bicolor* (Sorghum) | 545 | XP_002456213 |
| *SbMAX1c* | *Sorghum bicolor* (Sorghum) | 545 | XP_002453551 |
| *SbMAX1d* | *Sorghum bicolor* (Sorghum) | 540 | XP_002438586 |
| *DcMAX1* | *Daucus carota subsp. sativus* (Carrot) | 526 | KZN06895 |
| *CmMAX1* | *Cucumis melo* (Muskmelon) | 531 | XP_008452735 |
| *AoMAX1a* | *Asparagus officinalis* (Asparagus) | 552 | XP_020251236 |
| *AoMAX1b* | *Asparagus officinalis* (Asparagus) | 549 | XP_020251248 |

**Table S6.** Accession numbers of CYP722Cs used for the phylogenetic tree construction in Fig. S11. The amino acid sequences are downloadable from NCBI, except for TpCYP722C, which is downloaded from https://plants.ensembl.org/index.html. LjCYP722C1 is downloaded from https://lotus.au.dk/

| Gene | Species | Size (a.a.) | Accession numbers |
|---|---|---|---|
| GaCYP722C | *Gossypium arboreum* (Tree cotton) | 491 | XP_016745621 |
| VuCYP722C | *Vigna unguiculata* (Cowpea) | 494 | XP_027918387 |
| SlCYP722C | *Solanum lycopersicum* (Tomato) | 486 | XP_004232430 |
| GmCYP722C | *Glycine max* (Soybean) | 494 | XP_003548874 |
| TpCYP722C | *Trifolium pratense* (Red clover) | 492 | Tp57577_TGAC_v2_mRNA22267 |
| GbCYP722C | *Gossypium barbadense* (Egyptian cotton) | 491 | KAB2051573 |
| DzCYP722C | *Durio zibethinus* (Durian) | 490 | XP_022741804 |
| StCYP722C | *Solanum tuberosum* (Patato) | 495 | XP_006340657 |
| CaCYP722C | *Capsicum annuum* (Red bell pepper) | 490 | XP_016560669 |
| ErgCYP722C | *Erythranthe guttata* (Seep monkeyflower) | 483 | EYU24120 |
| GrCYP722C | *Gossypium raimondii* (New world cotton) | 491 | XP_012436859 |
| CcCYP722C | *Citrus clementina* (Clementine) | 485 | ESR38866 |
| CsCYP722C | *Citrus sinensis* (Sweet orange) | 497 | XP_006467149 |
| EugCYP722C | *Eucalyptus grandis* (Flooded gum) | 486 | XP_010029758 |
| CusCYP722C | *Cucumis sativus* (Cucumber) | 484 | XP_004150006 |
| PpCYP722C | *Prunus persica* (Peach) | 491 | XP_020424098 |
| PmCYP722C | *Prunus mume* (Japanese apricot) | 491 | XP_008241894 |
| MbCYP722C | *Malus baccata* (Siberian crab apple) | 497 | TQE02546 |
| FvCYP722C1 | *Fragaria vesca* (Woodland strawberry) | 481 | XP_004287873 |
| FvCYP722C2 | *Fragaria vesca* (Woodland strawberry) | 491 | XP_004303111 |
| LjCYP722C1 | *Lotus japonicus* (Birdsfoot trefoil) | 489 | LotjaGi4g1v0120600 |
| LjCYP722C2(DSD) | *Lotus japonicus* (Birdsfoot trefoil) | 484 | BBO93647 |
| CacCYP722C | *Cajanus cajanifolius* | 493 | XP_020238265 |
| RcCYP722C1 | *Ricinus communis* (Castor bean) | 492 | XP_002521537 |
| RcCYP722C2 | *Ricinus communis* (Castor bean) | 491 | XP_002524333 |
| MeCYP722C1 | *Manihot esculenta* (Cassava) | 506 | XP_021622147 |
| MeCYP722C2 | *Manihot esculenta* (Cassava) | 500 | XP_021600819 |
| VvCYP722C | *Vitis vinifera* (Grape) | 491 | XP_002269279 |
| ElgCYP722C | *Elaeis guineensis* (Oil palm) | 564 | XP_029117801 |
| OsCYP722B | *Oryza sativa* (Rice) | 518 | XP_015639511 |
| BdCYP722B | *Brachypodium distachyon* (Purple false brome) | 510 | XP_010230530 |
| SiCYP722B | *Setaria italica* (Foxtail millet) | 542 | XP_004960948 |
| PhCYP722B | *Panicum hallii* (Hall's panicgrass) | 551 | XP_025809022 |
| SbCYP722B | *Sorghum bicolor* (Sorghum) | 512 | OQU77193 |
| SlCYP722A | *Solanum lycopersicum* (Tomato) | 481 | XP_004238286 |
| AtCYP722A | *Arabidopsis thaliana* (Thale cress) | 476 | NP_173393 |
| StCYP722A | *Solanum tuberosum* (Potato) | 481 | XP_006342223 |
| CaCYP722A | *Capsicum annuum* (Red bell pepper) | 490 | XP_016568530 |
| AlCYP722A | *Arabidopsis lyrata* | 477 | XP_020869333 |

| | | | |
|---|---|---|---|
| | (Lyre-leaved rock cress) | | |
| EsCYP722A | *Eutrema salsugineum* (Saltwater cress) | 489 | XP_006416485 |
| CrCYP722A | *Capsella rubella* (Red shepherd's purse) | 481 | XP_006307378 |
| BrCYP722A | *Brassica rapa* (Field mustard) | 489 | RID51716 |
| TcCYP722A1 | *Theobroma cacao* (Cacao) | 488 | EOY14821 |
| TcCYP722A2 | *Theobroma cacao* (Cacao) | 488 | EOY14823 |
| GrCYP722A | *Gossypium raimondii* (New world cotton) | 498 | XP_012452669 |
| CcCYP722A | *Citrus clementina* (Clementine) | 523 | ESR38694 |
| CsCYP722A | *Citrus sinensis* (Sweet orange) | 498 | XP_006467169 |
| EugCYP722A | *Eucalyptus grandis* (Flooded gum) | 500 | XP_010060975 |
| CusCYP722A | *Cucumis sativus* (Cucumber) | 483 | XP_011649632 |
| PpCYP722A | *Prunus persica* (Peach) | 496 | XP_007226877 |
| PmCYP722A | *Prunus mume* (Japanese apricot) | 525 | XP_008220838 |
| FvCYP722A | *Fragaria vesca* (Woodland strawberry) | 493 | XP_011459540 |
| CasCYP722A | *Cannabis sativa* (Hemp) | 533 | XP_030484637 |
| LjCYP722A | *Lotus japonicus* (Birdsfoot trefoil) | 499 | AFK39117 |
| CiaCYP722A | *Cicer arietinum* (Chickpea) | 488 | XP_004501123 |
| CacCYP722A | *Cajanus cajanifolius* | 485 | XP_020219964 |
| RcCYP722A | *Ricinus communis* (Castor bean) | 489 | XP_002510423 |
| MeCYP722A | *Manihot esculenta* (Cassava) | 508 | XP_021600571 |
| VvCYP722A | *Vitis vinifera* (Grape) | 496 | CAN64558 |
| NnCYP722A | *Nelumbo nucifera* (Sacred lotus) | 481 | XP_010263599 |
| MaCYP722A | *Musa acuminata* (Banana) | 491 | XP_009390176 |

**Table S7.** Sequences of genes used in this study

| Gene | Sequence (5′–3′) |
|---|---|
| *PpMAX1a* | ATGGGTGTTTTGGAAGTTCTGTTGTTCACCAACGTTTCTTTGTTGTCTACCATCT<br>TCACCGTTTTGGCTATTTTGGCTGGTGTTTTAGGTTACTTGTATGGTCCATATTG<br>GGGTGTTAGAAGAGTTCCATCTCCACCAACTATTCCATTGGTTGGTCATTTGCC<br>ATTATTGGCTGAATATGGTCCAGATGTTTTCTCTGTTTTGGCAAAGCAATACGGT<br>CCAATCTTCAGATTTCATATGGGTAGACAGCCATTGATTATCGTTGCTGATGCTG<br>AATTGTGTAGAGAAGTTGGTATCAAGAAGTTCAAGGACATCCAGAACAGATCTA<br>TCCCATCTCCAATTTCTGCTTCTCCATTGCATCAAAAGGGTTTGTTCTTTACCAG<br>AGATGCTAGATGGTCTGCTATGAGAAACACCATTTTGTCTGTTTACCAACCATCT<br>CATTTGGCCTCATTGGTTCCAACTATGCAATCCTTTATTGAATCCGCTACTGAAA<br>AGCTGGGTTCCTCTAAAGAAGAAGATATCACCTTCTCCAACCTGTCTTTGAGATT<br>GACTACTGATGTTATTGGTCAAGCTGCTTTCGGTGTTAATTTCGGTTTGTCTAAG<br>CCACAATCCATCTCCGATTCTATCAACAAACAAATCGGTGGTTCCCAAGATATCA<br>ACAACGTTGAAGTTTCCGATTTCATCAACCAGCATATCTACTCTACTACCCAATT<br>GAAGATGGACTTGTCTGGTTCCTTGTCCATTATTTTGGGTTTGTTGGTTCCAGTG<br>TTGCAAGAACCATTCAGACAAATCTTGAAGAGAATCCCAGGTACTATGGATTGG<br>AAGGTTGAAAGAACTAACAGAAAGTTGTCCGGTAGATTGGATGAATTGGTCGGT<br>AAAAAGATGAGAGATAGAGACAGAGGTTCCAAGGATTTCTTGTCCTTGATTATG<br>AACGCCAGAGATTCTGAAACCGTTTCCAAGTCTGTTTTCACCCCAGATTACATTT<br>CTGCTGTTACCTACGAACATTTGTTGGCTGGTTCTGCTACTACTGCTTTTACTTT<br>GTCATCCATCGTTTACTTGGTTGCTGGTCATCCAGAAGTTGAAGAAAAGTTGTT<br>GGCCGAAATTGATGGTTTTGGTCCACCAGATCAAATGCCAACTGCTCATGACTT<br>GCAACATAAGTTCCCTTACATCGACCAAGTTATCAAAGAAGCCATGAGGTTCTAT<br>ATGGTTTCACCTTTGGTTGCCAGAGAAACCTCTAGACAAGTTGAAATTGGTGGC<br>TACTTGTTGCCAAAAGGTACTTGGGTTTGGTTAGCTTTGGGTGTATTGGCTAAA<br>GATCCAAAGAATTTTCCAGAGCCAGATAAGTTCAGACCAGAAAGATTTGATCCA<br>TCCTGCAAAGAAGTTAAGCAAAGACATCCATACGCCTTCATTCCATTTGGTATTG<br>GTCCAAGATCATGCATCGGTCAAAAGTTCTCATTGCAAGAGTTGAAGTTGAGCT<br>TGATCCACTTGTACAGAAACTACGTTTTCAGACATTCTCCAGGTATGGAAAAGC<br>CATTGCAATTGGAATATGGTATCGTGCTGAATTTCAAGAACGGTGTTAAGTTGA<br>GAGTCATCAAGAGAACCTTGTTCCAACATGCTTGA |
| *PpMAX1b* | ATGGAACTGTTGATGTTGTCCGATGTTTGTTTGGTTTCTACCATCTTCACTTTGT<br>TGGCTATGTTGGCTGGTGTTGTTTTGGGTTACTTTTACAAACCATACTGGGGTG<br>TTAGAAGAGTTCCAGGTCCACCAGCTATTCCATTGATTGGTCATATTCCACTGTT<br>GGCTAAACATGGTCCAGATGTTTTTTCTGTTTTGGCCAAGCAATACGGTCCAAT<br>CTTTAGATTTCATATGGGTAGACAGCCATTGATCATAGTTGCTGATGCTGAATTG<br>TGTAGAGAAGTCGGTATTAAGAAGTTCAAGGACATCAAGAACAGGTCCATTCCA<br>TCTCCATTGGCTGCTTCTCCATTGCATCAAAAAGGTTTGTTCTTGACCAGAGATG<br>CTAGATGGTGGACTATGAGAAACACCATTTTGTCTGTTTACCAGCCATCTCATTT<br>GGCTTCATTGGTTCCAACTATGCAGTCCTATATTGAATCTGCTACCCAAAAGTTG<br>GACGACTGCTCTAAAGAAGAAGAGGAAGTTACCTTCTCCAACATCTCTTTGAGA<br>TTGGCTACTGATGTTATTGGTCAAGCTGCTTTTGGTGTTAACTTCGGTTTGTCTA<br>AGCCACAATCTATCACCGATTCCATCAACAAACAAATCGGTGGTCAAGATATCA<br>ACATCGATGAAGTTTCCGACTTCATCAACCAGCATATCTACTCTACTACCCAATT<br>GAAGTTGGACTTCTCTGGTTCCTTGTCCATTATTTTGGGTTTGTTGGTTCCTATC<br>TTGCAAGCTCCATTTTGGCAAATCTTGAAGAGAATTCCAGGTACGATGGACAGA<br>AAGATCGAAAGAAACAATCAGAAGTTGACCTCCAGATTGGATCAAATCGTTGAA<br>AAGAGAATGAAGGACTCTGGTGGTAGAGGTTCTAAGAATTTCTTGTCCTTGATTT<br>TGAACGCCAGGGAATCTGAAACTGTCTCCAAGAATTTTTTCACCCCAGATTACAT<br>TTCCGCCTTGACTTATGAACATTTGTTAGCTGGTTCTGCTACCACCTCTTTTACT<br>TTGTCATCTGTTGTTTACTTGGTTGCTGCTCATCCTGAAGTCGAAAAAAAGTTGT<br>TGGCTGAAATTGATGGTTTTGGTCCTCCAGCTCAAATGCCAACTGCTCATGACT<br>TGCAACATAAGTTCCCTTACATGGACCAAGTTATCAAAGAAGCCATGAGGTTCT<br>ATATGGTTTCACCATTGGTTGCTAGAGAAACCTCTAGACAAGTTGAAATAGGTG |

| | |
|---|---|
| | GTTACTTGTTGCCAAAAGGTACTTGGGTTTGGTTAGCTTTGGGTGTTTTGGCTAA<br>AGACCCAAAGAATTTTCCAGAACCACACAAGTTTAAGCCAGAAAGATTCGATCC<br>ATCCTGCAAAGAAGAAAAGCAAAGACATCCATACGCCTTCATTCCATTTGGTTTG<br>GGTCCAAGATCATGCATCGGTCAAAAGTTTTCATTGCAAGAGATCAAGCTGTCG<br>TTGATCCACTTGTACAGAAAGTATGTTTTCAGACACACCCCAAACATGGAAATCC<br>CATTGGAATTGGAATTCGGTATCGTCCTGAAATTCAAGTACGGTATCAAGTTGA<br>GAGTCATCAAGAGAACCTGA |
| *PpMAX1c* | ATGATCTCCATGGAACTGTTCACCAACGTTTCTTTGGTTTCTGCTCCAACTATTT<br>TCACCGTTTTGGCTATGTTGGTTGGTGTTTTGGGTTACTTGTATGGTCCATATTG<br>GGGTGTTAGAAAAGTTCCAGGTCCACCAGTTATTCCATTGGTTGGTCATTTGCC<br>ATTGATGGCTAAATATGGTCCAGACTTGTTCCAAATTTTGGCCAAGCAATACGGT<br>CCAATCTTCAGATTTCATATGGGTAGACAACCATTGGTTATCGTTGCTGATGCTG<br>AATTGTGTAGAGAAGTTGGTATCAAGAAGTTCAAGGACATCCCAAATAGATCAAT<br>GCCATCTCCAATTCAAGCTTGCCCATTGCATCAAAAGGGTTTGTTTTTTACCAGA<br>GATGCTAGATGGTCTACCATGAGAAACACCTTGGTTTCATTATACCAACCATCTC<br>ATTTGGCCTCATTGGTTCCAACTATGCAATCTTTTGTTGAAACCGCCACCAGAAA<br>CATCGATCCATCTAAAGAAGAAGAGGACATCACCTTCTCCAAGATTTCTTTGGCT<br>TTGGCTACTGATACAATTGCTCAAGCTGCTTTTGGTGTTAACTTCGGTTTGTCTA<br>AACCACAATCTACCTCCGATTCCGTTAACAAAATTGATGGTGGTAGACAAGATAA<br>GAACGACGCTGTTTCTAAGTTCACCGATCAATACGTTTACTCTGTTGCCCAAGTT<br>AAGATGGATTTGACTGGTTCCTTGTCCATTATCTTGGGTTTGTTGTTTCCAATCC<br>TGCAAGAACCATTCAGACAGATCTTGAAAAGAATCCCAGGTACTATGGATTGGA<br>AGATTGAAAGAACTAACGACAACTTGTCTGGTAGGTTGGACGAAATCGTTGAAA<br>AGAGAATGAAGGATTCCGACAGAGGTTCTAAGGATTTCTTGTCCTTGATTTTGAA<br>CGCCAGGGAATCTGAAAAGGTTTCCAAGAAAGTTTTCACCCCAGATTACATTTC<br>TGCCTTGACTTACGAACATTTGGTTGCTGGTTCTGCTACTACTGCTTTTACTTTG<br>TCTACTACCGTCTACTTGATCTCTCAATATCCAGAAGTCGAGAAGAAGTTGTTGG<br>AAGAATTGGATGCATTTGGTCCAGCAGATCAAATGCCAACTGCTCATGACTTGC<br>AAAACAAGTTTCCATACGTTGACCAGGTCATCAAAGAATCTATGAGGTTGTATCC<br>AGTCTCTCCATTGATTGCTAGAGAAACCTCTTCCGAAGTTGAAATTGGTGGTTAC<br>GTTTTGCCAAAAGGTACATGGGTTTGGTTCGGTGTTGGTGTTATTGCTAAAGAT<br>CCAAAGAATTTCCCAGAGCCAAACAAGTTCAAGCCAGAAAGATTTGATCCAAAC<br>TGCAAAGAAGAGAAAGAAAGACATCCATACGCTTTCTTGCCATTTGGTATTGGT<br>CCAAGAGCTTGTTTGGGTCAAAAGTTTGCCTTGCAAGAAATCAAGTTGGCCTTG<br>ATTCACCTGTACCGTAAATACGTTTTTAGGCACTCTCCAAACATGGAAAAGCCAT<br>TGGAATTTGAGTTCGGTATCATCTTGGATTTCAAGAACGGTGTTAAGTTGAGAG<br>CTTTGAAGAGAACCCAAAAGTTCTAA |
| *PavMAX1c* | ATGATCTCCATGGAACTGGTTACCAACGTTTCTTTGGTTTCTGCTCCAACTATTT<br>TCACCGTTTTGGCTATGTTGGTTGGTGTTTTGGGTTACTTGTATGGTCCATATTG<br>GGGTGTTAGAAAAGTTCCAGGTCCACCAGTTATTCCATTGGTTGGTCATTTGCC<br>ATTGATGGCTAAATATGGTCCAGACTTGTTCCAAATTTTGGCCAAGAAGTACGG<br>TCCAATCTTCAGATTTCATATGGGTAGACAACCATTGGTTATCGTTGCTGATGCT<br>GAATTGTGTAGAGAAGTTGGTATCAAGAAGTTCAAGGACATCCCAAATAGATCA<br>ATGCCATCTCCAATTCAAGCTTGCCCATTGCATCAAAAGGGTTTGTTTTTTACCA<br>GAGATGCTAGATGGTCTACCATGAGAAACACCTTGGTTTCATTATACCAACCAT<br>CTCATTTGGCCTCATTGGTTCCAACTATGCAATCTTTTGTTGAAACCGCCACCAG<br>AAACATCGATCCATCTAAAGAAGAAGAGGACGTTACCTTCTCCAAGATTTCTTTG<br>GCTTTGGCTACTGATACAATTGCTCAAGCTGCTTTTGGTGTTAACTTCGGTTTGT<br>CTAAACCACAATCTAACTCCGATTCCGTTAACAAAATTGATGGTGGTGGTCAAG<br>ATAAGAACAACGATAACAAGAATGACGCTGTTTCTAAGTTCACCGATCAATACGT<br>TTACTCTGTTGCCCAAGTTAAGATGGATTTGACTGGTTCCTTGTCCATTATCTTG<br>GGTTTGTTGTTTCCAATCCTGCAAGAACCATTCAGACAGATCTTGAAAAGAATCC<br>CAGGTACTATGGATTGGAAGATTGAAAGAACTAACGACAACTTGTCTGGTAGGT<br>TGGACGAAATCGTTGAAAAGAGAATGAAGGATTCCGACAGAGGTTCTAAGGATT<br>TCTTGTCCTTGATTTTGAACGCCAGGGAATCTGAAAAGGTTTCCAAGAAAGTTTT<br>CACCCCAGATTACATTTCTGCCTTGACTTACGAACATTTGGTTGCTGGTTCTGCT |

| | |
|---|---|
| | ACTACTGCTTTTACTTTGTCTACTACCGTCTACTTGATCGCACAATATCCAGAAG<br>TCGAGAAGAAGTTGTTGGCTGAATTGGATGCATTTGGTCCAGCAGATCAAATGC<br>CAACTGCTCATGACTTGCAAAACAAGTTTCCATACGTTGACCAGGTCATCAAAG<br>AATCTATGAGGTTGTATCCAGTCTCTCCATTGATTGCTAGAGAAACCTCTTCCGA<br>AGTTGAAATTGGTGGTTACGTTTTGCCAAAAGGTACATGGGTTTGGTTCGGTGT<br>TGGTGTTATTGCTAAAGATCCAAAGAATTTCCCAGAGCCAAACAAGTTCAAGCC<br>AGAAAGATTTGATCCAAACTGCAAAGAAGAGAAAGAAAGACATCCATACGCTTT<br>CTTGCCATTTGGTATTGGTCCAAGAGCTTGTTTGGGTCAAAAGTTTGCCTTGCA<br>AGAAATCAAGTTGGCCTTGATTCACCTGTACCGTAAATACGTTTTTAGGCACTCT<br>CCAAACATGGAAAAGCCATTGGAATTTGAGTTCGGTATCATCTTGGATTTCAAGA<br>ACGGTGTTAAGTTGAGAGCTTTGAAGAGAACCCAAAAGTTCTAA |
| *FveMAX1a1* | ATGGGCATGGAATTCTTGATTACCAACGGTTCTTTGGTGTCTACCATTTTTACTG<br>TTTTGGCAGTTTTGGCTGGTGTCTTGGGTTACTTGTATGCTCCATATTGGGGTG<br>TTAGAAGAGTTCCAGGTCCAAGAACTATTCCATTTTTGGGTCATTTGCCTTTGTT<br>GGCTAAACATGGTCCAGATTTGTTCTCCGTTTTGGCTAAGCAATACGGTCCAAT<br>TTTCAGATTCCATATGGGTAGACAACCATTGATTATCGTTGCTGATGCTGAATTG<br>TGTAGAGAAGTCGGTATTAAGAAGTTCAAGGACATCCCAAACAGATCCATTCCA<br>TCTCCAATTTCTGCTTCTCCATTGCATCAAAAGGGTTTGTTCTTCACCAGAGATA<br>TTAGATGGTCTACCATGAGAAACACCATCTTGTCTGTTTACCAACCATCTTACTT<br>GGCTTCATTGGTTCCAACTATGCAGTCCTATATTGAATCTGCTACCCAAAACTTG<br>GACGACTCCTCTAACAAAGAAGATATCACCTTCTCCAACCTGTCTTTGAGATTG<br>GCTACTGATGTTATTGGTCAAGCTTCTTTCGGTGTTGATTTCGGTTTGTCTAAGC<br>CACAATCCATCTCTAACTCCATCAACAAGGTTGATAACGGTGAAGATAAGAACG<br>ATGATGAAGTCTCCGATTTCATCAACCAGCATATCTACTCTACTACCCAATTGAA<br>GATGGACTTGTCTGGTTCCTTGTCCATTATTTTGGGTTTGTTGGTCCCAATCATC<br>CAAGAACCTTTCAGACAAATCTTGAAGAGAATCCCAGGTACTATGGATTGGAAA<br>GTCGACAGAACTAATCAAAATTTGGCCGGTAGATTGGACCAAATCGTTATGAGA<br>AGAATGAAGGATTCCGACAGAGGTACTAAGAACTTCTTGTCCTTGATTATGAAC<br>GCCAGAGAATCTGAAGGTGTTGCTAAGTCTGTTTTCACCCCAGATTACATTTCT<br>GCTGTTACCTACGAACATTTGTTAGCTGGTTCTGCTACTACCTCTTTCACTTTGT<br>CATCTGCCGTTTACTTGATTTCTGGTCATCCAGAAGTTGAGTCTAAGTTGTTGGC<br>TGAAATTGATGGTTTTGGTCCACATGATCAAATTCCAACTGCTCCAGATCTGCAA<br>CATAAGTTTCCATACTTGGAACAGGTTATCAAAGAAGCCATGAGGTTCTATTTGG<br>TTTCACCATTGGTTGCTAGAGAAACCTCTAGAGAAGTTGAAGTTGGTGGTTACTT<br>ATTGCCAAAAGGTACTTGGGTTTGGTTAGCTTTGGGTGTTTTAGCAAAGGATCC<br>AAAGAATTTCCCAGAACCAGAAAAGTTCAAGCCAGAAAGATTTGACCCAAACGG<br>TAAAGAAGAAAAGCAAAGACATCCATACGCCTTCATTCCATTTGGTATTGGTCCA<br>AGAGCTTGCATCGGTCAAAAGTTTTCTCTGCAAGAGTTGAAGCTGTCATTGATC<br>CACTTGTACCGTAAATACGTTTTCAGACACTCTCCAAATATGGAAGCTCCATTGG<br>AATTGGAATTCGGTATCGTTTTGAACTTCAAGAAGGGTGTTAAGTTGACCGTCAT<br>TAAGAGAACCTGA |
| *FveMAX1a2* | ATGGGTATGGAATTCTTGATCACTAACGGTTCCTTGGTTAGTACCACCTTGACC<br>GTCTTGGCCGTTTTGGCTGGTGTTTTGGGTTACTTGTACGCTCCATACTGGGGT<br>GTTAGAAGAGTTCCAGGTCCTCGTACAATTCCGTTTTTGGGTCATTTGCCATTGT<br>TAGCTAAGCACGGTCCAGACTTATTCTCAGTTCTTGCGAAGCAATACGGTCCAA<br>TTTTCCGTTTCCACATGGGTAGACAACCATTGATTATTGTCGCTGATGCTGAATT<br>GTGTCGTGAAGTTGGTATCAAGAAGTTCAAAGACATTCCAAACAGATCTATTCCA<br>TCTCCAATTTCCGCTTCTCCACCACACCAAAAGGGTTTATTCTTCACCAGAGACA<br>CGAGATGGTCCACTATGAGAAACACTATTTTGTCAGTTTACCAACCATCTTACTT<br>GGCATCTTTGGTTCCAACCATGCAATCTTACATTGAATCTGCTACTCAAAACTTG<br>GACGATTCATCCAACAAGGAAGATATCACTTTCTCTAACTTGTCCTTGAGATTGG<br>CTACCGATGTTATCGGTCAAGCTGCTTTCGGTGTCGATTTCGGTTTGTCCAAAC<br>CACAATCCATCTCCGACTCTATTAACAAGGTCGACAACGGTGAAGACAAGAACG<br>ATGATGAAGTTTCTGACTTCATTAACCAACACATCTACTCCACTACTCACCTCAA<br>GATGGACTTGTCCGGTAGCCTGTCCATCATATTGGGTTTGTTAGTCCCAATTAT<br>CCAAGAACCATTCAGACAAATGTTGAAGCGTATCCCAGGTACTATGGACTGGAA |

| | |
|---|---|
| | GGTTGACAGAACCAACCAGAAGCTAGCCGGTAGATTAGACCATATCGTTATGAG<br>AAGAATGAAGGATTCTGATAGAGGTACCAAGAACTTCTTGTCTCTAATTTTGAAC<br>GCTCGTGAATCTGAAGGTGTGGCTAAGAGTGTCTTCACCCCAGATTACATCTCC<br>GCTGTTACTTACGAACACCTTTTGGCTGGTTCTGCCACCACTTCTTTTACTTTGT<br>CTTCTGCTGTCTACCTCATCTCTGGTCATCCAGAAGTTGAATCCAAGTTGTTGG<br>CCGAAATCGACGGTTTCGGACCTCACGACCAAATCCCAACTGCTCACGATTTGC<br>AACACAAGTTCCCATATTTGGAACAAGTCATCAAGGAAGCTATGAGATTCTACTT<br>GGTCTCTCCATTGGTTGCCAGAGAAACCTCTAGAGAAGTCGAAGTCGGTGGCT<br>ATTTATTACCAAAGGGTACCTGGGTCTGGTTGGCCCCAGGTGTTCTAGCTAAGG<br>ACCCAAAGAATTTTCCTGAACCAGAAAAATTTAAGCCAGAAAGATTTGATCCAAA<br>TGGTAAGGAAGAAAAGCAAAGACATCCATACGCTTTCATCCCATTCGGTATCGG<br>TCCAAGAGCTTGTATTGGTCAAAAATTCTCCTTGCAAGAATTGAAACTATCTTTG<br>ATTCACTTGTACAGAAAGTACGTTTTCAGACACTCTCCAAACATGGACACTCCAT<br>TGGAATTAGAGTTCGGTATTGTCTTGAACTTCAAGAACTCCGTTAAGTTGACTGT<br>CATCAAGAGGACCTGA |
| *MdMAX1a1* | ATGATGAAGAGCTTAGAAGAGGAGGAGACCTTGCAAAAGATAGGCGGCATGGA<br>AGTTTTATTCGCAACCAGCGTCTGGCTTATAAGCACAATCTTTACAGGATTAGCG<br>ATTCTTGGAGGCATCCTGGGTTATCTTTATGGGCCTTACTGGGGAGTGCGTAGA<br>GTACCAGGTCCCCCTACCATACCATTGTTAGGGCATTTGCCTTTGCTTGCTAAA<br>TATGGACCCGACGTTTTTTCTGTCCTGGCAGAGCAGTACGGCCCAATTTTCAGG<br>TTCCACATGGGGAGACAGCCGCTGATTATCGTGGCCGATGCTGAATTGTGCAG<br>GGAGGTAGGGATAAAAAAATTCAAAGATATACCTAATCGTTCTATACCGGCCCC<br>CATTTCAGCTAGTCCATTGCATCAAAAAGGATTGTTCTTTACGCGTGACGCAAG<br>ATGGTCAACAATGAGAAACACAATTCTTAGCGTGTACCAACCTTCCCACCTGGC<br>CTCACTGGTCCCGACAATGCAGTCATTTATCGAATCAGCAACAGAGAAACTACA<br>TTCAAGCAAAGAAGAAGAGATAACATTCTCCACTCTGTCACTTGGGTTGGCAAC<br>CGACGTATTGGGACAGGCCGCGTTCGGTGTCAATTTTGGATTGAGCAATACACA<br>TTCCATTGGCAAGTCCATCAAAGGGGAAGATAACGATGATGAAGTTAGCGATTT<br>CATTAACCAACATATTTACAGTACTACGCAATTGAAAATGGATTTAAGCGGCTCA<br>TTGTCAATTATTTTGGGCCTACTTGTTCCCATCTTGCAAGAGCCGTTCCGTCAGA<br>TCTTTAAGCGTATCCCTGGTACTATGGATTGGAAAGTTGAACGTACGAACCGTA<br>AGTTGAGCGGCAGACTTGATGGTATTGTGGCGAAGAGAATGCAAGACAGAGAG<br>AGGGGTTCCAAGGATTTTCTGTCAGTCATCATGAATGCAATGGAGAGCGAAGCC<br>GTATCTAAGAATGTCTTCACTCCTGATTATATCTCAGCTGTCACTTATGAACACC<br>TACTTGCTGGGAGCGCGACTACCGCCTTCACACTAAGCTCAATAGTATATTTGG<br>TAGCAGGCCACCCGGAGGTGGAGAAAAAACTTTTGGCAGAAATTGATGGATTC<br>GGCCCGCCAGATCAGATGCCTACAGCGCACGATCTACAGCATAAATTTCCCTAT<br>ATCGACCAGGTTATCAAAGAAGCTATGAGATTTTATATGGTGAGCCCACTGGTT<br>GCGAGGGAAACGTCTGGAGACGTTGAAATCGGAGGATACTTACTTCCGAAAGG<br>AACCTGGGTGTGGCTTGCGCTGGGTGTGCTAGCAAAAGACCCCAAGAATTTTC<br>CAGAGCCGGACAAGTTTCGTCCGGAGCGTTTCGATCCAAACTGTCAGGAACTA<br>AAACAACGTCACCCCTACGCTTTTATACCGTTTGGTATCGGCCCGAGATCCTGT<br>ATTGGCCAAAAATTTTCATTGCAAGAACTTAAGCTGAGTCTTATTCATTTGTACA<br>GAAAATATGTGTTCAGACACAGTCCCGACATGGAGAAGCCATTGGAATTAGAGT<br>ACGGTATTGTACTTAATTTCAAAAAGGGAGTTAAACTGCGTGTAATTAAGAGAAC<br>TTGA |
| *MdMAX1a2* | ATGGAATTGCAATTGTTGTTCACTAACGGTTTCTTGGTATCACCATCCACTTTGT<br>TTACAGCCCTAGCCATGTTCGCTGGTGTCTTGTCTTACCTATATGGTCCATACTG<br>GAGAGTCAGAAGAGTTCCAGGTCCACCAGTTATCCCCCTGTTGGGTCATTTGCC<br>ATTGTTGGCTAAGTACGGTCCAGATGTCTTCTCTGTCTTGGCCAAGCAGTACGG<br>TCCATTGTTCAGATTTCACATGGGTAGACAACCTTTGATTATTGTCGCTGATGCA<br>GAATTGTGTAGAGAAGTCGGTATCAAGAACTTCAAGGATATCAACAATAGATCTA<br>TTCCATCTCCAATTGCTGCTTCCCCATTGCACCAAAAGGGTTTATTCTTCACCAG<br>AGACGCTAGATGGTCTACTATGAGAAACACTATCCTATCCTTGTACCAACCATCT<br>CATTTGGCTTCTTTGGTCCCAACCATGCAATCTTTCATCGAAAGTGCTACTCAAA<br>AGTTGGACTCCTCTAAGGAAGCTGAAGACATCACTTTCTCTAACTTGTCCTTGA |

| | |
|---|---|
| | GATTGGCCACTGATGTTATTGGTCAAGCTGCCTTCGGTGTTAACTTTGGTTTGT<br>CGAAGTTGGAATCCATCAACGAATCCATTGACAAGATTGATTCTAAGGACAACA<br>ACGATGATGACCAAGTTTCTGACTTTATTAACCAACACATCTACTCTACCACTCA<br>ATTGAAGATGGACTTGTCTGGTTCTTTTTCTATCATTCTTGGTTTGTTGGCTCCT<br>ATCTTACAAGAACCATTCCGTCAAGTTTTAAAGAGAATTCCAGGTACCATGGACT<br>GGAAGGTTGAGAGAACCAACAGAAAGTTAAGTGGTCGTCTGGACGAAATTGTT<br>GTTAAGAGAATGAAGGATTCCGAAAGAGGTTCTAAGGATTTCTTATCCTTAATCA<br>TGAACGCTAGAGAATCTGAAACTGTTGCTAAAAACGTTTTCACCCAAGACTACAT<br>CTCTGCCGTTACTTACGAACACTTGTTGGCTGGTTCCGCTACCACCTCTTTCAC<br>CTTGTCCTCCACCGTCTACTTAGTGGCCGGTCACCCAGAAGTCGAACGTAAATT<br>GTTGGCTGAAATTGATGGTTTCGGCCCTCCAGATCAAATGCCAACTGCTTACGA<br>TTTGCAACACAAATTCCCATATATTGACCAAGTCATCAAGGAAGCTATGAGATTC<br>TACTTAGTCTCCCCATTGGTTGCTCGTGAAACTTCCAGAGAAGTTGAAATCGGT<br>GGTTACTTGTTACCAAAGGGTACTTGGGTTTGGCTCGCCCTCGGTGTCTTAGCT<br>AAGGACCCAAACAATTTCCCAGAGCCAGACACCTTCAGACCAGAAAGATTCGAC<br>CCTGACTGTAACGAAGAAAAGCAAAGACATCCATACGCTTTCATCCCATTTGGT<br>ATTGGTCCAAGAGCTTGTATCGGTCAAAAATTCTCCTTGCAAGAATTGAAGTTGT<br>CTTTGATTCACTTGTACAGAAAATACGTTTTCAGACACTCTCCAAACATGGAAAA<br>GCCATTGGCTTTGGAATACGGCATCGTTTTGAACTTCAAGAATGGTGTTAAGGT<br>CAGAGTTGTCAAGCGTTGA |
| *MdMAX1b1* | ATGATTGGTTCTATTGAAATGTTATTCACCATCTTCATGGTCTTAGCTATCTTGG<br>GAGGTGTTTTGGGTTACTTGTACTGGCCTTACTGGGGTATGAGAAGAGTCCCAG<br>GTCCACCTACTATTCCATTGGTCGGTCACATCCCATTGTTAGCCAAATACGGTC<br>CAGATGTCTTCTCCGTTTTGGCTAAGGAATACGGTCCAATCTTTAGATTCCACAT<br>TGGTAGACAACCACTAGTCATTGTCGCTGACGCTGAATTATGTCGTGAAGTTGG<br>TATCAAGAAGTTCAAGGACATGAAAAACAGAAGCATCCCATCCCCACTCGCTGC<br>TTCTCCATTGCACCAAAAGGGTTTGTTTTTGACCAGGGATGCTAGATGGGCTAC<br>AATGAGAAACACTATTTTGTCCATGTACCAACCATCTCACTTGGCTAATTTGGTT<br>CCAACCATGCAATCTTTCGTTGAATGTGCTACTCAAAAGTTAGCTTCGACCGCT<br>GAAGAAGAAGAAGATATCACTTTCTCTGACATCTCTTTGCAATTGACCACTGACG<br>TAATCGGTCAAGCTGCTTTCGGTGTTAACTTTGGTTTGTCCAAGCCACAATCCTT<br>GGGTGAATCTGTCGAAAAAATTGACGGTAAGTACAACAACGATGATGAAGTCTC<br>CGATTTCATCAACCAACACATGTACTCCACTGCTCAACTCAAATTGGACTTCAGT<br>GGTTCCTTATCTATCATCTTGGGTCTTCTGGTCCCAATTCTACAATGGCCATTCT<br>GGCAAATGTTGAAGAGAATTCCAGGTACCATGGACAGAAAGATTGAAAGATCTA<br>ACTGGCATTTGGTCTCTAGATTGGATGCCGCTGTTGAAAAGAGAATGAAGGACT<br>CTGGTAGAGGTTCTCGTGATTTCTTGTCATTGATATTGAACGCTCGTGAATCCG<br>GTCGTGTTTCCAAGAACGTTTTCACCCAAGACTACATTTCTGCTTTGGCTTACGA<br>ACACTTACTAGCCGGTTCCGCTACCACCGCCTTCACTCTCTCTTCTGTTGTCTA<br>CTTGGTTGCTGGTCATCCAGAAGTTGAAAAGAAGTTGTTGGCTGAAATCGATGG<br>TTTCGGTCCACCAGACCAAATGCCAACTGCCACTGACTTGCAACACAACTTCCC<br>ATACTTGGACCAAGTTGTTAAGGAAGCTATGAGATTCTACATGGTTTCTCCATTG<br>GTCGCCAGAGAAACTTCTAAGGAAATTGAAATCGGGGGTTACTTGTTGCCAAAG<br>GGTACCTGGGTTTGGTTGGCATTGGGTGTCTTGGCCAAGGATCAAAAGAATTTC<br>CCAGAACCAGAGAAGTTCAGACCAGAAAGATTCGACCCATCTTGTAAGGAAGAA<br>AAGCAAAGACACCCTTACGCTTTCATCCCATTTGGTTTAGGTCCAAGAACTTGC<br>ATTGGTCAAAAATTTGCTGTCCAAGAAATCAAGTTGGCTTTGATCCACTTATATA<br>GAAAGTACGTTTTCAGACATTCCCCATTCATGGAAGTTCCTTTGGAATTGGAATA<br>TGGTATTGTTTTGAAGTTCAAGAACGGTGTCAAGTTGAGAGTTATTAAGAGAACC<br>TGA |
| *MdMAX1b2* | ATGATTGGTTCCATTGAATTGTTGTTCAACATTTTCATGGTCTTGATCGTTTTGG<br>GTGGTATTTTGGGTTACTTGTACGGTCCATACTGGAGAGTTAGAAGAGTTCCAG<br>GTCCTCCAACCATCCCTTTGGTCGGTCACATTCCATTGATGGCTAAATATGGCC<br>CAGACATCTTCTCTGTCTTTGCTGAAGAATACGGTCCTATTTTTCGTTTCCACAT<br>TGGTAGACAACCATTGATTATCGTTGCTGATGCTGAATTGTGTAGAGAAGTTGG<br>TATCAAAAAGTTCAAGGATATCAAGAACAGATCCATCCCACCACCATTGGCCGC |

| | |
|---|---|
| | TTCTCCATTGCACCAAAAGGGTTTATTTTTGACAAGAGACGCTAGATGGGTTACT<br>ACCAGAAACACTATCCTAAGCGTTTACCAACCATCTCATTTAGCTAACTTGGTTC<br>CAACTATGCAATCTTTCGTTGAATGTGCCACTGAAAAATTGGATTCCATGGCCAA<br>GGTTGAGGAAGATATCACTTTCTCCGCCATCTCTTTGCGTTTAGCCACCGACGT<br>CATCGGTCAAGCTGCTTTCGGAGTTAACTTTGGTTTATCTAAGCCACAATCCGT<br>CGGTGAAAGTGTCGAAAAATTCGATGGTAAGGACAATAATGATGACGAAGTCTC<br>TGATTTCATCAACCAACATATCTACAGTACTACTCAAGTCAAGATGGACTTCTCC<br>GGGTCCTTGTCCATCATTTTAGGTTTGCTGGTCCCAATCCTACAATGGCCATTCT<br>GGCAAATGTTGAAGAGAATTCCAGGTACCATGGACGGTAAGATCGAACACAACA<br>ACAGACACTTGACCTCTAGATTAGGTGCTGTGGTTGAAAAGAGAATGAAGGACT<br>CTAGAAGAGGTTCTAAAGATTTCTTGTCTGTTATTTTGAACGCCCGTGAATCCGG<br>TCGTCTATCCAAGAACGTTTTCACCACCGACTACATTTCTGCTTTGGCTTACGAA<br>CACTTATTGGCTGGTTCTGCTACCACTGCCTTTACTTTGTCTTCCGCCGTCTACT<br>TGGTTGCGGGTCACCCAGAAGTCGAAAAGAAGTTGTTAGCTGAAATTGACGGTT<br>TCGGTCCACCTGACCAAATGGCTACCGCTTCTGACTTGCAACACAAGTTCCCAT<br>ACTTGGATCAAGCTAGATTGTTGAAATTCCAATGTTGTATTGAAGAATCATCTTC<br>TTTGGTCATAAAGGAAGCTATGAGATTCTACATGGTTTCCCCACTAGTTGCTCGT<br>GAAACTTCCAGAGATGTTGAAATTGGTGGTTACATCTTGCCAAAGGGTACCTGG<br>GTCTGGTTGGCTTTGGGTGTCTTGGCTAAGGATCAAAAGAACTTCCCAGAACCA<br>GAAAAGTTCAGACCAGAAAGATTCGACCCATCTTGTAAGGAAGAAAAGCAAAGA<br>CATCCATACGCTTTCATTCCGTTCGGTCTGGGTCCAAGATCTTGCATCGGTCAA<br>AAGTTCGCTATCCAAGAAATCAAGTTGGCATTGATTCACTTATACAGAAAGTACG<br>TCTTTAGACACTCCCCATATATGGAATCTCCACTCGAATTGGAATTCGGTATTGT<br>CTTAAAGTTCAAGAACGGTGTTAAGTTGAGAGTTATCAAGAGAAAGTGA |
| *GhiMAX1* | ATGGCGCTGGTTCACGAGACAATACAGAGCAGTGTGGGTGCGGAAGGATTCTG<br>GAGCTATTATCCATACGGCACGACCGTAAGCATAATCATTACGGTGCTAGCCAT<br>GGTCTTAGGCTATCTGTACGGCCCCTACTGGAAGGTACGTCGTGTACCTGGGC<br>CACCCACGATTCCTTTAGTCGGTCACCTGCCACTAATGGCTAAGTATGGGCCG<br>GATGTTTTCTCCGTGCTAGCAAAGCAATACGGGCCGATCTTTCGTTTTCACATG<br>GGCAGGCAACCATTAATTATTGTAGCCGATGCTGAATTGTGTAAAGAGGTCGGA<br>ATTAAGAAGTTTAAGGACATCCCTAATCGTTCTATTCCCAGCCCCATCGCCGCC<br>AGTCCACTGCATCAAAAAGGGTTATTCTTCACGAGGGACGCGAGGTGGAGTAC<br>GATGAGGAACACCATTCTGAGTGTCTACCAGCCTTCTCACTTAGCGTCCTTAGT<br>TCCTACGATGCAGAATTACATCGAATCCGCGACGGAAAATTTGCATAGTAGCAA<br>ACAGGATATTGTTTTCAGCAATTTAAGCCTAAAGCTTGCCACAGACGTCATTGG<br>GCAGGCAGCTTTTGGGGTGAATTTCGGCCTATCTAAACCGCAATCCATAAACGA<br>ATCTATTAGAAACGTCGACAAGCAAGGAAGCCAAGATGATGAGGTGTCAGATTT<br>CATTAACCAACATATCTACTCTACGACTCAGCTTAAAATGGATCTGTCCGGATCT<br>GTATCTATCATTCTAGGCTTATTAGTACCAATTTTACAAGAGCCATTCAGGCAGA<br>TCCTTAAAAGGATTCCTGGGACCATGGATTGGAAAGTAGAGAGGACGAACAAAA<br>AGCTGTCCGGCAGACTAGATGAGATCGTGAGTAACAGAATGAAGGATAAGAAC<br>CGTGGTAGTAAAGATTTCCTGTCACAGATACTTTCTGCCAGAGAATCTGAAAAT<br>GTCGCAAAGAACGTCTTTACCCCGGACTACATCTCAGCAGTCACATATGAACAT<br>TTATTGGCCGGCAGTGCCACGACATCATTCACATTAAGTTCTATTGTTTACTTAG<br>TGGCTGGGCATCCGGAAGTGGAAAAGAAATTGGTTGCGGAGATTGACGGGTTT<br>GGGCCCCACGATCAGGTTCCGACTGCCTACGATCTACAACATAAGTTTCCTTAC<br>CTAGACCAAGTTATCAAGGAAGCCATGAGATTCTACATTGTGTCACCGTTAGTC<br>GCGCGTGAAACCTCCAAGGAGGTTGAAATTGGTGGGTACCTTTTACCTAAAGG<br>GACCTGGGTGTGGCTGGCAGTGGGGGTTCTGGCCAAAGATCCTAAAAATTTTC<br>CCAACCCGGACAAGTTTATCCCCGAAAGGTTTGATCCAAATTGCGAGGAAGAAA<br>AGCAGCGTCACCCGTACGCCCTAATTCCCTTCGGGATCGGCCCTAGGGCCTGT<br>GTCGGAAGAAAATTTTCACTGCAGGAGATAAAATTAAGTCTTATCCACTTATACA<br>GGAAGTACACTTTCCAACATTCTAGTACAATGGAGAACCCCCTTGAATTAGAATA<br>TGGCATAGTACTGAATTTTAAACATGGCGTGAAATTGACGGTCACGAAACGTAC<br>CTAA |

| | |
|---|---|
| *HaMAX1* | ATGGAGAAATCTATGATTACAGAAGTGGTGGGGCTTGTCAACTTTACGAGTTCC<br>TTCTTCACGATCGTAGCTCTTTTAGCTGGGGTTTTAGTTTATTTTTACCAGCCCA<br>TATGGGGAGTGAGGAAAGTTCCAGGCCCACCGACTATTCCATTACTAGGCCAC<br>TTGCCTTTACTAGCACAGCATGGCCCGGACCTTTTCGCCGTACTAGCAAAAAGG<br>TATGGGCCAATCTACAGGTTTCACATGGGGAGGCAGCCACTTGTAATAGTAGCC<br>GATGCTGAGCTATGCCGTGAAGTCGGCATTAAGAAATTCAAGGACATCCCTAAC<br>AGGTCAATCCCCTCTCCGATATTGGCTAGCCCCCTGCACAGAAAAGGGCTGTTT<br>TGGATACGTGATAGTAGATGGTCAACGATGAGGAACACGATTGTTAGCGTTTAT<br>CAGCCCTCTTACTTGGCGAAGTTAGTGCCCATGATGCAAAGTTTCATCGAGACG<br>ACGTCTCAGAATATCCCGACGGATGACCAAGACATTGTATTTAATGAGTTTTCTC<br>TGAAAATGGCTACCGATGTCATTGGTAAAGCGGCCTTCGGTTTCGATTTCGGGC<br>TATCCAACCCTAACAGCCATCAAAATAACGACAAACACACAGAGAGTTTCATCAA<br>GCAGCACATATACTCCACCACAATGTTGAAGATGGACCTATCTGGTTCATTCTC<br>AATAATTCTAGGTTTATTGCTGCCTATCCTGCAGGAACCAGTTCGTCAGATTTTA<br>AAAAGGATACCCTACACCATGGACTGGAAGATAGAAAGGACTAACAAAAATCTA<br>AACTCCCAATTAGATCAAATCGTAGTAAACAAGATGAAGGAGGAGCAGAGAGGT<br>AGCAAGGACTTCTTGAGTCTTATTCTGAACGCAAGAGAATCCGAGACAGTAAGT<br>AAGAACCTATTTACGCCAGACTATCTGAGTTCTATCGCTTACGAACACCTACTAG<br>CTGGTTCCGCCACCACTTCATTCACTCTATCATCTTGCGTCTACTTAATCTCTGG<br>GCACCCGGAGGTGGAGAAGAAGCTGTTGAAGGAGATTGACGAATTTGGACCCC<br>CAGATCAAGTCCCGACTGCGGATGACCTTCAGAGTCGTTTCCCCTACCTTGACC<br>AAGTAGTTAAGGAGGTAATGAGATTCTATGTCGTCTCTCCCTTAATAGCTCGTGA<br>GACAAGCACGCAGGTAGAGATCGGTGGATATATCTTACCAAAAGGTACGTGGG<br>TATGGTTAGCGGTTGGCGTTCTTGCTAAAGACCCTAAAAATTTCCCGGATCCTG<br>AAAAGTTCAAGCCCGAGAGGTTCGACCCAAACTGTGATGAGGAGAAGCGTAGG<br>CATCCATATGCTTTCATCCCTTTCGGAATCGGGCCGAGAGTCTGTATCGGTCAA<br>AAGTTCTCTTTACAAGAAATCAAACTGACGTTGATACACCTGTACCAAAGGTATG<br>TGTTCAGACATTCTCCAAACATGGAGTCTCCGTTAGAATTCGACTTTGGCATCGT<br>CTTAAACTTCAAAAATGGGGTCAAAGTGAGAGCGATTAAGAGAACGTAG |
| *MaMAX1a* | ATGGAAATGGGTTCCACGACTATTACATTGGGGTGGGCAGAGTATGCGTTCAC<br>GCTTGTAGCCCTATTAGTTGGCTTTTTGATGTATCTGTATGCCCCGAGCTGGGG<br>TGTTAGGAAAGTACCCGGTCCTCCAACGATCCCACTTCTGGGACACTTGCCCTT<br>ATTAGCAAAGCACGGCCCAGACGTCTTCTCCGTGTTAGCGCAGACGTACGGGC<br>CAGTCTTTAGATTCCACATGGGAAGGCAACCTATGGTAATTGTGGCGGACCCC<br>GAGTTATGCAAGCAAGTTGGAATTAAGAAATTCAAGAGTGTGCCGAATCGTAGT<br>CTTCCTACCCCTATATCAGGGTCTCCTTTGCATCAAAAAGGATTATTCTTTACCA<br>GGGATTCACGTTGGTCCGCTATGAGGAACACAATAATTTCATTATATCAACCTAC<br>GCATCTGGCAGGGTTGATCCCAACGATGCAATCATACATCGACTCAGCTACCAG<br>GCACCTTTCATCCACCCAGAAAGACGACGTGACGTTTAGCGATTTGTCTCTGCG<br>TTTGGCTACTGATGTCATAGGAGAGGCAGCTTTCGGCGTTGATTTCGGCCTAAG<br>TGGCGGCTCACCCCCCGATGGGGAACGTTCAAAAGATGGCGAGGTTTCCGAGT<br>TTATTAAGGAACACATCTACTCTACTACTTCACTGAAGATGGATCTTTCAGGCTC<br>TTTCTCCATAGTGTTAGGTCTTTTGGTACCTGTATTACAAGAGCCTGTTAGGCAA<br>CTACTGAAGAGGATACCTGGTACCGCAGATTGGAAGATACATCAGACGAACCTA<br>AAGTTATCAAAGCGTCTGGAAGAGATCGTGGCGAAAAGAGCCAGCGAGCGTGC<br>CAGGGGTAGCAAAGACTTCCTGTCCGCCATACTTAATGCAAGGGACAGCGATG<br>GTGCCAGTAGAAAGTTGTTCACCTCAGACTATATCTCTGCTCTGGCATATGAGC<br>ACCTACTGGCAGGCTCTGCAACAACGTCTTTTACCTTAAGCTCAGTCCTTTACTT<br>GGTATCCGAACATCCTGAAGTGGAGCAGAAATTACTTCGTGAAATTGACGGATT<br>TGGCCCAAGTGACCTTATTCCTACATTCGATGATCTTCAGCACGAATTTCCCTAT<br>CTTGATCAAGTTATCAAAGAGGCGATGAGGTTTTATACAGTTAGCCCACTGGTG<br>GCACGTGAGACGAGTCAGCCCGTCGAGGTGGGCGGGTTCCTACTACCGAAAG<br>GTACGTGGGTCTGGATGGCCCCGGGTGTGCTAGCTAAAGACCCCAAGCATTTC<br>CCCGAACCGCACCTATTTAGGCCCGAGAGATTCGACCCGGCGTGTGATGAGGA<br>AAAACAGAGACACCCATACGCCCATATTCCATTTGGGATTGGTCCTCGTGCGTG<br>CATTGGACAAAAATTCTCCTTGCAGGAAATAAAGCTAGCGGTTATTCATTTATAC |

| | |
|---|---|
| | CGTCGTTATGTTTTCAGGCATTCTCCTAGCATGGAATCACCTTTGGAGTTCCAAT ACGGTGTAGTACTGAATTTCAAACATGGCGTCAAACTTCGTGTGATCCGTCGTT CAGAAGCATTTCACGTAATTATTTGA |
| *MaMAX1b* | ATGGAGGACAGGGTCCTGGAAATCGTGGAGTCTGTGGAGTCCTTGGTCCGTTC CTGCGCACCGTTCTTCTTAGCGTCCATGGCCCTGCTTTTGGGATTTTTAGTCTAT TTCTACGCCCCATATTGGAGGGTTAGAAGGGTACCAGGTCCTCCCACGACTTTC CCTTTGGGTCACATACCGTTTCTAGCGAAACACGGCCCGGATATTCTGAGGGTT TTCGCAAAAAACTATGGACCAATATTTAGGTTCCATTTGGGTAGGCAACCGTTG GTGATTGTTGCCGATGCGGAACTATGCAGGAAGGTCGGTATTAAGAATTTCAAG GATATACGTAACAGGTCTAGCCCCAGCCCTGCTACAGGATCCCCTCTGCTGCA GAATGGATTATTTCTTCTGAGGGACAGCCGTTGGACTTCCACCAGAAACATCAT AACATCTCTATATCAGCCCGCGCACTTAGCATCTTTGATCCCAACAATGCATCAC TACGCTACGTCTTTTTGCCATACTATCAGTACGATACAACGTAATCGTGAGGAC GTTCCATTCAGTGAACTGTCCTTAAGGCTGGCGATAGACATAATTGGCAAGACG GCATTTGGGATAGAATTCGGCTTGCTGAACGATGATGACGACGATGACGATGG ATCTTGCTTCCTGAGGCAGCATACATATGCAATCTCCTCATTAAAGATGGATCTA AGCGGATCTCTGAGCACAGCGCTGGGTTTGATTGCGCCGGTTTTACAGAACCC GTGTAGGGAGATTTTTAAGAGGATTCCTGGCGCAGCGGACTATAAATTGCACCA GATGAATCAACAACTGTGCGACAGAATTGACGCAATAATTGCAAAGCGTTCCAG CGAGATGACTAGAGAGAGTAAAGATTTTTTAGCAGCTTTACTGAACTCCAGGCA AACGCGTCTTGCTGAAAACCTACTGACGGACTCATACGTTCGTGCCCTAGTCTA CGAACATCTTATTGCAGGCACTAAAACGACTGCTTTCACTCTTGCGATGACGGT ATATCTAGTCTCTCGTCACCCGGATGTGGAGAAAAAACTAGTAGATGAGATTGA CCGTTTCGGCCCAAGAGACTTGATACCCACTTTCGACGATCTGCATAGCAAATT CCCTTATTTAGACCAAGTGATAAAGGAATCCATGAGGATGTATACAGTCTCCCC ATTGGTCGCAAGGGAAACTTCTCAGCAGGTTGAGATTGGAGGCTATGTCTTACC GAAAGGGACATGGGTCTGGCTAGCCCTGGGTGTTGTTGCAAAGGATTCTAAGC AGTTCCCGGCTCCAGACGTATTTAGGCCGGAAAGATTTGATCCCGCTCGTGAT GAGGAGAAACGTCGTCATCCCTATGCTCACATACCGTTCGGTATTGGCCCCCG TGCATGCATTGCGCAGAAATTCGCCATTCAAGAGGTGAAATTGGCCCTTATACA ACTATATAGGCACTACGTGTTCCGTCGTTCCCCAGAAATGGAACTACCCCCGGA ATTCCAGTATGGGCTGATACTGTCTTTTAAAAGAGACATAATGCTAAGAGCTATA AAACGTGCGAATGATTAG |
| *AmtMAX1* | ATGGCGTTTATGGAAGGTTTTCTACTACCCTTATTTCGTGTTACCAGCCCCTACT TTAGCATAAGGAACTTGTTCGAGAGATGGGTTCCTTTGATATGCACCGTCATTG CGATAGTCGTCGGCATTATATACGTATACTTTTCTGCGCCATCATGGAGAGTCA GAGCCGTGCCAGGTCATCCGACAAGGTGGTTCCTTGGGCATCTTCACCTGTTG GCAAAGCATGGCCCAGGTGTCTTCAGTGTTCTTGCGAAAGAATATGGTCCTATC TATCGTTTCCATATGGGTAGGCAACCACTGGTTATCATAGCAGATGCAGAGTTA TGTCGTGAAGCGGGCATAAAGAAGTTCAAAGACCTACCTAACAGAAGCATACCA TCCCCAATTTCTGCGAGTCCACTACACCTGAAGGGACTGTTCTTTAGCCGTGGA TCAAGATGGTCTATAATGAGGAATACAATAACCTCACTTTATCAGCCGAGCCACT TGGCTAGCCTTATCCCAACGATGCATTTTTTCATTGCTTCCGCGAGTAGTATGTT GGAATCCCATCTACAGAGAAAAGAGGATGATGTTAATTTCAGCGAGCTGACGCT AAAGTTAACGACAGACGTAATTGGCAGAGCAGCTTTTGGGGTGAACTTCGGCTT GACCGGATCATCCTCAGCTCAATCCGGTCAGAACACACTTAAGAAGGATGAAAA CTCTGAGGACGGCTATATCTCTAAATTCCTGAATCAACACATTTATTCCACTACA TCTTTGAAGATGGACTTAACCGGCACTTTCAGTATAATACTTGGCCTTTTCTTCC CTATCTTGCAAAATCCTGTCAGGTGGCTTCTGTGTAGAATTCCTGGAACTGCTG ATAGGGAACACGAACTTGCGAATAAAGAATTATCCCAGAGAGTAGATGAAGTTG TCGAGAAGCGTTGTAAAGAGATCGAGGGCGGAAGTTCAAATATCGACCTATTAA CCGCCGTGCTAAACGCGCAGGAATCATACAGAGCCATCAGCAAGAAAATTTTTA CACCTGAATGCGTATCAAGCTTGGCTTACGAGCATCTTCTGGCCGGAAGCGCC ACAACGTCATTCACGATGGCGGCTACCGTTTACCTAGTGTCAGCGCACCCTGAA GTTGAGGAAAAGCTATTACGTGAGATCGATACTTTTGGGCCCCCAAATAAGAGG CCAGACGCAGAAGACCTAAGGCTTAATTTCCCATATTTAGATATGGTAGTCAAG |

| | |
|---|---|
| | GAAGCGATGCGTTTCTTCACTGTGAGCCCTTTAGTTGCCAGGGAGGCCGAGCG<br>TGACGTACAAATCGGGGGGTACTTACTGCCAAAGGGGACGTGGATTTGGCTGG<br>CCTTGGGTGTACTAGCAAAGGATCCTTCAAATTTCCCCGAGCCCGAGAAGTTCA<br>AGCCGGAGAGATTTGACCCCCAAGGGGAAGAGGAAAAAAAGAGGCACCCGTAT<br>GCACATATTCCTTTTGGTATAGGACCCAGGGCATGTGTTGGAGGAAAATTCTCA<br>ATTCAGGAGATTAAACTAACACTGATATACTTATATCAGCGTTACATTTTTAGGC<br>ACTCACCGAATATGGAATCACCACTTGAGTTCGACTATGGCGTTGTTTTAAATTA<br>CAAGCATGGCGTGAAACTTCGTATTTTGAAGAGGTCACGTGTGTAG |
| *PgMAX1* | ATGGCCTCTCTATGTGGACTACTGACGATTTTTAGTACAGAGACTGACAGGTTT<br>ATTTCAACACAGGACCAATTCATGAATACGACGACTATACTGATTTGCGTCTTTA<br>TATTGGCCGCAGCCTCAATTACCGCTTGGATCTACCTTGCTATTCCAACGTGGA<br>AGGTTCGTCGTGTACCATCTCCCCCGGCCTTCTGGCTACTGGGCCACCTGCCT<br>CTTCTAGCGAAACACGGTCCTGAAGTGTTTATTCAGTTGGCTAGAAAGTACGGT<br>CCCATCTACAGGTTTAACATAGGACGTCAGCCGTTGGTCATTATTGCCGATGCT<br>GATCTTTGCCGTGAGGTTGGAATAAAGAAATTCAAGCAATTCTCTAACCGTTCTA<br>TCCCTTCTCCGATCGCTTCAAGTCCCCTTCATCAGAAAGGACTTTTCTTTACGAG<br>GGATTCAAGGTGGAGTTCAATGCGTGGTGCAATTCAACCCTTATATCAGACTGG<br>TAGGATTAGTAATCTGCTACCTGTCATGGAGAGGGTGGTCTGTGTTTTGAAGCG<br>TAAATTGGCAGCCAAAGAAAAAACTGATGACATAGACTTTTCCGAGCTGCTGCT<br>GCGTGTAGCAACTGACATTATAGGCGAAGCAGCTTTTGGGGAGAGGTTCGGCT<br>TGACCGAGGAGACAACAGCCATTAGTTCTTCTAACCCGGCAGAAGTTTCAGAAT<br>TTATAAAACAACATGTTTACAGCACCTCTTCACTGAAGATGGACCTGAATGGTAC<br>TTTTTCCATTCTTGCGGGGATTCTATTCCCCATCGCCCAGGAACCTTTTAGACAA<br>ATCCTTAGCCGTATACCCGGCACTGGGGATTGGAAGGTGTGCATTAATAATCGT<br>CGTCTTACACATCGTCTAAACGCTATCGTAGAAAAAAGGAAAAAAGACGTGGTA<br>GGAAAAGAGAAGAGAATGGATTTCTTAAGCACGGTTACAGGGTCTAAGTTTTCC<br>CGTGAACTGTTTTCTGAAGAGTACATATCAGCGCTGACATATGAACACCTTTTG<br>GCGGGATCCGCCACAACGTCATTCACAATATCAGTAATACTGTATTTGGTGTCC<br>GCCCATCCAGACGTAGAATCTAAGCTGCTAAGGGAGATCGACGAATTCGGCCC<br>GCCGGATCGTAACCCAGCTGCTGAAGACTTGGACATTAAGTTTCCTTACTTGAC<br>CCAAGTCATCAAAGAGGCCATGCGTTTTTACACCGTATCCCCTTTGGTTGCTCG<br>TGAGGCGTCTGAGCCCGTTCAGATTGGGGGGTACACTTTGCCTAAGGGAACGT<br>GGGTGTGGATGGCTCTGAATGCCTTGGCCAAGGACCCCCGTTACTTTCCCGAG<br>CCAGAGATGTTCAACCCGGAGCGTTTCGATCCTGAGTGTGAAGAAGAAAAGAAT<br>CGTCATCCCTACGCGAATTCACCTTTCGGAATTGGTCCGCGTGCATGCATTGGG<br>ATGAAATTTGCCTTCCAAGAGATTAAAGTCGTCCTTATACATTTGTACCAACTAT<br>ATACATTCGACCATTCCCCTGCTATGGAGAATCCCCTAGAGTTCCAATTCGGAA<br>TCGTTGTCTCTGTGAAGTACGGGATCCGTTTGAGACTGAGACATCGTAGAGCG<br>CAATCTCCAGTGTAA |
| *PpCYP722C* | ATGTTGAATCTATCCGAAGAAGGTTTATTATTCTTGGTCAGAAACTACTACGATG<br>TCTTCATTGTCGCCGTCTTCTCCATTGGTGTTACCTTCTTGGTTTCCAAGATTGC<br>TTGGAGAAGCTTGGACATGACTACTACCTCTTACAGAGGTGGTGGTATCCCAGG<br>TAGATTGGGTTTGCCATTCGTTGGTGAAACCTTGTCATTGTTATCCGCCACCTC<br>CTCTATCAAGGGTTGTTACGAATTCGTTCGTTTGCGTAGAATCTGGCACGGTAA<br>GTGGTTCAAGACCAGAATTTTCGGTCAAATCCATGTTTTCGTCCCATCCACTGAA<br>GGTGCCAGAGCTATCTTTTCCGACGATTTCGCCAAGTTCAACAAGGGCTATGTC<br>AAGTCCATGGCTGACTGTGTGGGTGAAAAGTCTCTCTTGTGTGTTCCACACGAA<br>GACCACAAGAGAATCAGACACTTGTTATCTGAACCTTTCTCTATGAATTCTTTGT<br>CTACTTTCGTCCAAAAGTTCGACAAGGTTTTGTGCCAAGAATTGAAGAAATTAGA<br>AGGTGGTGGTAAATCTTTCGTCGTCTTGGATTTTTCCATGAAGATTACTTTCGAT<br>GCTATGTGTAACATGTTGCTCAGTGTTACTGATGATTCTTTGTTGAGAAAGATCA<br>ACAAGGACTGTACTGCTGTTTCTGATGCCATGTTGACCTTCCCATACATGATTCC<br>TGGAACCAGATACTACAAGGGTATTAAGGCTCGTCGTAGATTGATGGAAACCTT<br>TAAGGATATCATCGGTAGAAGAAGATCTGGTAAGGAATCTGCTGAAGATTTTCT<br>ACAATCCATGTTGGAAAGAGACAGCCACCCACCAAACGAAAAGTTGCAAGATTC<br>TGAAATCATGGACAACTTGTTAACATTGATTATTGCTGGTCAAACCACTACCTCT |

| | |
|---|---|
| | GCTGCTATGATGTGGTCTGTTAAATTCTTGGACGAAAACAGAGAAGCTCAAGAG<br>AGACTTAGAGAAGAACAATTGTCCATCGCTAGAGCCAGACCAGACGGTGCTTCT<br>GCCACTTTGGAAGACTTCAAGAACATGCCATACTGTTTGAAAGTTGTCAAGGAA<br>ACTTTGCGTATCTCTAACGTCCTTTTGTGGTTCCCAAGAGTTGCTTTATCTGATT<br>GCACCATTGAAGGTTTCGAAATTAAGAAGGGTTGGCACGTTAACATCGACGCTA<br>CCTGTATCCACAACGACCCAGCTTTGTACGCTGAACCAATGCAATTCAACCCAT<br>CTAGATTCGATGAAATGCAAAAGCCATACTCCTTCATTCCATTCGGTTCTGGTCC<br>AAGAACTTGTTTGGGTATGAACATGGCTAAGGTCACCATGTTAGTTTTTCTACAT<br>AGATTGACTTCCGGTTACAGATGGACTGTTGACGACTTGGACACCTCCTTGGCT<br>AGAAACGCTCACATTCCACGTTTGCGTTCCGGTTGTCCAATCACTTTGAGAGCT<br>TTGTGA |

**Table S8.** Primers used for site-directed mutagenesis of PpMAX1c

| Primer name | Mutation | Sequence (5'-3') |
|---|---|---|
| PpMAX1cM1QI66SV-For | QI66SV | ACTTGTTCTCCGTTTTGGCCAAGCAATACGG |
| PpMAX1cM1QI66SV-Rev | QI66SV | TGGCCAAAACGGAGAACAAGTCTGGACCAT |
| PpMAX1cM2R108A-For | R108A | TCCCAAATGCTTCAATGCCATCTCCAATT |
| PpMAX1cM2R108A-Rev | R108A | GCATTGAAGCATTTGGGATGTCCTTGAA |
| PpMAX1cM3Q115A-For | Q115A | CTCCAATTGCTGCTTGCCCATTGCATCA |
| PpMAX1cM3Q115A-Rev | Q115A | GGCAAGCAGCAATTGGAGATGGCATTG |
| PpMAX1cM4C117S-For | C117S | TTCAAGCTTCTCCATTGCATCAAAAGGGT |
| PpMAX1cM4C117S-Rev | C117S | GCAATGGAGAAGCTTGAATTGGAGATGG |
| PpMAX1cM5R136A-For | R136A | CTACCATGGCTAACACCTTGGTTTCATTA |
| PpMAX1cM5R136A-Rev | R136A | AGGTGTTAGCCATGGTAGACCATCTAG |
| PpMAX1cM6L139I-For | L139I | GAAACACCATCGTTTCATTATACCAACC |
| PpMAX1cM6L139I-Rev | L139I | ATGAAACGATGGTGTTTCTCATGGTAG |
| PpMAX1cM7V159I-For | V159I | AATCTTTTATTGAAACCGCCACCAGAAA |
| PpMAX1cM7V159I-Rev | V159I | CGGTTTCAATAAAAGATTGCATAGTTG |
| PpMAX1cM8R164K-For | R164K | CCGCCACCAAGAACATCGATCCATCTAA |
| PpMAX1cM8R164K-Rev | R164K | CGATGTTCTTGGTGGCGGTTTCAACAA |
| PpMAX1cM9K179N-For | K179N | CCTTCTCCAACATTTCTTTGGCTTTGGC |
| PpMAX1cM9K179N-Rev | K179N | AAGAAATGTTGGAGAAGGTGATGTCCT |
| PpMAX1cM10A183K-For | A183K | TTTCTTTGAAGTTGGCTACTGATACAAT |
| PpMAX1cM10A183K-Rev | A183K | TAGCCAACTTCAAAGAAATCTTGGAGA |
| PpMAX1cM11T188I-For | T188I | CTACTGATATTATTGCTCAAGCTGCTTTT |
| PpMAX1cM11T188I-Rev | T188I | GAGCAATAATATCAGTAGCCAAAGCCAA |
| PpMAX1cM12A190G-For | A190G | ATACAATTGGTCAAGCTGCTTTTGGTGTT |
| PpMAX1cM12A190G-Rev | A190G | CAGCTTGACCAATTGTATCAGTAGCCAA |
| PpMAX1cM13K226D-For | K226D | CTGTTTCTGATTTCACCGATCAATACGTT |
| PpMAX1cM13K226D-Rev | K226D | CGGTGAAATCAGAAACAGCGTCGTTCTT |
| PpMAX1cM14T228I-For | T228I | CTAAGTTCATTGATCAATACGTTTACTCT |
| PpMAX1cM14T228I-Rev | T228I | ATTGATCAATGAACTTAGAAACAGCGTC |
| PpMAX1cM15Y231H-For | Y231H | CCGATCAACATGTTTACTCTGTTGCCC |
| PpMAX1cM15Y231H-Rev | Y231H | AGTAAACATGTTGATCGGTGAACTTA |
| PpMAX1cM16VA235TT-For | VA235TT | TTTACTCTACCACCCAAGTTAAGATGGATTTG |
| PpMAX1cM16VA235TT-Rev | VA235TT | TAACTTGGGTGGTAGAGTAAACGTATTGATC |
| PpMAX1cM17F254V-For | F254V | GTTTGTTGGTTCCAATCCTGCAAGAACCA |
| PpMAX1cM17F254V-Rev | F254V | GGATTGGAACCAACAAACCCAAGATAAT |
| PpMAX1cM18D281A-For | D281A | GAACTAACGCCAACTTGTCTGGTAGGTT |
| PpMAX1cM18D281A-Rev | D281A | ACAAGTTGGCGTTAGTTCTTTCAATCT |
| PpMAX1cM19H335Q-For | H335Q | CTTACGAACAATTGGTTGCTGGTTCTG |
| PpMAX1cM19H335Q-Rev | H335Q | CAACCAATTGTTCGTAAGTCAAGGCA |
| PpMAX1cM20T350V-For | T350V | TGTCTACTGTTGTCTACTTGATCTCTCAA |
| PpMAX1cM20T350V-Rev | T350V | AGTAGACAACAGTAGACAAAGTAAAAGC |
| PpMAX1cM21Q356G-For | Q356G | TGATCTCTGGTTATCCAGAAGTCGAGAA |
| PpMAX1cM21Q356G-Rev | Q356G | CTGGATAACCAGAGATCAAGTAGACGG |
| PpMAX1cM22Y357H-For | Y357H | TCTCTCAACATCCAGAAGTCGAGAAGAAG |
| PpMAX1cM22Y357H-Rev | Y357H | CTTCTGGATGTTGAGAGATCAAGTAGAC |
| PpMAX1cM23L368I-For | L368I | TGGAAGAAATTGATGCATTTGGTCCAGC |
| PpMAX1cM23L368I-Rev | L368I | ATGCATCAATTTCTTCCAACAACTTCT |
| PpMAX1cM24V390L-For | V390L | TTCCATACTTGGACCAGGTCATCAAAGA |
| PpMAX1cM24V390L-Rev | V390L | CCTGGTCCAAGTATGGAAACTTGTTTT |
| PpMAX1cM25L400F-For | L400F | CTATGAGGTTCTATCCAGTCTCTCCATT |
| PpMAX1cM25L400F-Rev | L400F | CTGGATAGAACCTCATAGATTCTTTGA |
| PpMAX1cM26P402M-For | P402M | GGTTGTATATGGTCTCTCCATTGATTGC |

| | | |
|---|---|---|
| PpMAX1cM26P402M-Rev | P402M | GAGAGACCATATACAACCTCATAGATT |
| PpMAX1cM27S413K-For | S413K | AAACCTCTAAGGAAGTTGAAATTGGTGG |
| PpMAX1cM27S413K-Rev | S413K | CAACTTCCTTAGAGGTTTCTCTAGCAA |
| PpMAX1cM28FG430LA-For | FG430LA | GGGTTTGGTTGGCTGTTGGTGTTATTGCTAAA |
| PpMAX1cM28FG430LA-Rev | FG430LA | CACCAACAGCCAACCAAACCCATGTACCTTT |
| PpMAX1cM29I435L-For | I435L | TTGGTGTTCTGGCTAAAGATCCAAAGAA |
| PpMAX1cM29I435L-Rev | I435L | CTTTAGCCAGAACACCAACACCGAACC |
| PpMAX1cM30N446E-For | N446E | CAGAGCCAGAGAAGTTCAAGCCAGAAAG |
| PpMAX1cM30N446E-Rev | N446E | TGAACTTCTCTGGCTCTGGGAAATTCT |
| PpMAX1cM31E462Q-For | E462Q | AAGAGAAACAAAGACATCCATACGCTTT |
| PpMAX1cM31E462Q-Rev | E462Q | GATGTCTTTGTTTCTCTTCTTTGCAGT |
| PpMAX1cM32P470A-For | P470A | CTTTCTTGGCTTTTGGTATTGGTCCAAGA |
| PpMAX1cM32P470A-Rev | P470A | TACCAAAAGCCAAGAAAGCGTATGGAT |
| PpMAX1cM33L479A-For | L479A | GAGCTTGTGCTGGTCAAAAGTTTGCCTTG |
| PpMAX1cM33L479A-Rev | L479A | TTTGACCAGCACAAGCTCTTGGACCAA |
| PpMAX1cM34G480A-For | G480A | CTTGTTTGGCTCAAAAGTTTGCCTTGCAA |
| PpMAX1cM34G480A-Rev | G480A | ACTTTTGAGCCAAACAAGCTCTTGGACC |
